# Phylogenomics clarifies *Balanophora* evolution, plastid reduction, and obligate asexuality origins

**DOI:** 10.1101/2025.03.29.646081

**Authors:** Petra Svetlikova, Huei-Jiun Su, Kenji Suetsugu, Filip Husnik

## Abstract

Holoparasitic plants are non-green plants that depend entirely on their host plants for essential resources, which often results in functional reduction and gene loss. However, the timing and extent of gene loss associated with parasitism remain unclear. Although *Balanaphara* is known to have highly reduced plastid genomes, only five species from a few geographically restricted regions have been studied. Here, we expanded *Balanaphara* sampling with seven species from twelve populations across Taiwan and Japan. We assembled their plastid genomes and transcriptomes and inferred multi-gene phylogenies from diverse plastid and nuclear markers. The uncovered relationships within the genus imply independent origins of obligate apomixis in island populations of several species. All the plastid genomes are highly reduced (14-16 kbp), gene-dense, completely syntenic, strongly AT-biased (87-88%), and with the same non-canonical genetic code (TAG->Trp) as previously reported in other *Balanaphara* species. To further understand the functional role of these non-photosynthetic plastids, we predicted the subcellular localization of nuclear-encoded proteins based on expression data. Strikingly, over 800 *Balanaphara* proteins were predicted to be plastid-targeted, suggesting a retained capacity for the biosynthesis of amino acids, fatty acids, riboflavin, lipoic acid, heme, isoprenoids, and components of the glycolysis and pentose phosphate pathways. Our findings show that the extreme plastid genome reduction in *Balanaphara* mainly occurred before the origin of the clade and primarily erased photosynthesis-related functions. Numerous highly expressed nuclear genes of *Balanaphara* still retain chloroplast transit peptides, and the resulting proteins are likely targeted to the non-photosynthetic organelle that remains metabolically active. Similar to other parasitic eukaryotes with non-photosynthetic plastids, such as apicomplexans, the loss of photosynthesis led to extreme plastid genome reduction without massive elimination of other photosynthesis-unrelated functions. Balanophoraceae thus emerges as a fascinating model for reconstructing the evolutionary changes associated with photosynthesis loss in land plants.

## Introduction

The Balanophoraceae family comprises exclusively parasitic plants that exploit their host plants for all essential resources (Hansen 1972; Yu et al. 2021; Cardoso and Braga 2024). Although closely related to mistletoes, Balanophoraceae are completely non-photosynthetic root holoparasites (Heide-Jorgensen 2008; Su et al. 2015). The family contains 13 genera, with *Balanophora* being the most frequently studied, comprising 25 species (POWO 2024). The highest diversity of *Balanophora* is in humid forests of the subtropical and tropical regions of Asia-Pacific (Figure S1). Southern China hosts the highest number of species, at least 15 (Kuijt and Hansen 2014; Yu et al. 2022).

Similar to non-photosynthetic single-celled eukaryotes, such as Apicomplexa or heterotrophic plants that once had photosynthetic ancestors, the genus *Balanophora* has experienced significant plastid reduction. The plastid genomes of *Balanophora*, along with those of several other parasitic and mycoheterotrophic plants, were proposed to be only one step from the complete organelle genome loss (Barrett and Davis 2012; Graham et al. 2017; Hadariová et al. 2018). Despite the loss of most plastid-encoded genes, the non-photosynthetic organelle is still present and, due to protein import from the cytoplasm, was predicted to be involved in the biosynthesis of compounds such as amino acids, heme, and fatty acids (Su et al. 2019; Chen et al. 2023). However, only a few *Balanophora* plastid genomes and transcriptomes have been assembled so far, limiting our understanding of the timing and evolution of the plastid reduction. *Balanophora* plastids retain *accD, clpP, ycf1, ycf2, trnE*, ribosomal protein-coding genes (*rpl2, rpl14, rps2, rps3, rps4, rps7, rpsll, rpsl2, rpsl4, rpsl8*, and *rpsl9*), and 3-4 rRNA genes as shown for *B. reflexa* and *B. laxiflora* (Su et al. 2019). These plastomes are not only about ten times smaller than those of photosynthetic plants but also extreme in their compactness, GC content, and codon usage bias. Moreover, the entire Balanophoraceae family is prone to genetic code reassignments: four genera within the family have been shown to use non-canonical genetic codes in their plastid genome. This alternative code, where the TAG/TGA codons have been reassigned from stop codons to tryptophan (Su et al. 2019; Ceriotti et al. 2021; Kim et al. 2023), occurs among extreme genomes, but it is the only such case described from land plants.

The relationships among most of the 13 Balanophoraceae genera have been tested with molecular data (Su et al. 2015; Ceriotti et al. 2021; Sanchez-Puerta et al. 2023) and found to correspond with the morphological classifications (Hansen 1980; Takhtajan 1997; Kuijt and Hansen 2015). According to these studies, *Balanophora* is sister to *Langsdorffia*, and together with *Thonningia*, they form the Balanophoroideae subfamily. Several molecular phylogenetic analyses have explored the taxonomy and speciation of *Balanophora* species. However, these studies were limited by sparse taxon sampling, with no more than ten and usually only five or six species included (Su et al. 2012; Yu et al. 2021; Yu et al. 2022; Kim et al. 2023). Based on these results and morphological observations, two major groups of *Balanophora* are currently recognized: the subgenus *Balania*, which includes *B. tobiracola*, the *B. harlandii* complex, and the *B. involucrata* complex; and the subgenus *Balanophora*, which includes *B. fungosa, B. subcupularis, B. appressa, B. parajaponica, B. japonica, B. laxiflora*, and *B. yakushimensis* (Su et al. 2012; Yu et al. 2022).

The reproductive biology of Balanophoraceae is also extraordinary but surprisingly understudied. *Balanophora* plants emerge above ground only during flowering when their ‘fungus-like’ inflorescence appears (Su et al. 2012; Thorogood and Santos 2020). Their inflorescence is considered a cluster of the most morphologically reduced flowers known (Eberwein et al. 2009) and bears either only unisexual flowers, or both female and male flowers. *Balanophora* species are predominantly sexual, with two species consisting solely of female individuals (*B. japonica* and *B. nipponica*) (Kuwada 1928; Su et al. 2012). These female-only species reproduce strictly asexually via apomixis (obligate apomixis). Several additional *Balanophora* species include female-only populations, although these species also occur in other areas where they reproduce sexually (Hansen 1972; Su et al. 2012; Yu et al. 2021).

Although apomixis (seed formation without fertilization) has been documented in at least 80 plant families and 300 genera (Carman 1997; Hojsgaard et al. 2014; Albertini 2019), obligate apomixis is very rare. Most cases of apomixis are facultative, with seeds produced through a combination of sexual reproduction and apomixis (Hogan 1983; Schmidt and Antlfinger 1992; Teppner 1996; Huang et al. 2009). Theoretical models predict that reliance solely on apomixis can lead to the accumulation of deleterious mutations and potentially negatively impact plant survival (Kimura et al. 1963; Muller 1964). This could explain why ancient obligate apomictic species are nearly nonexistent (Whitton et al. 2008). The only known isolated apomictic lineage is the monotypic genus *Houttuynia*, while all other angiosperm apomicts are confined to the tips of the tree of life, with no genera exclusively composed of obligate apomicts (Whitton et al. 2008). Thus, a detailed investigation of the multiple obligate apomictic Japanese and Taiwanese species could offer insights into how obligate apomictic species diversify.

Here, we sampled seven of the eight known *Balanophora* species from 12 populations in Taiwan and Japan, including six obligately apomictic populations (Figures 1 and 2), assembled their plastid genomes and nuclear transcriptomes, and inferred their phylogenetic positions. The assembled plastomes are completely syntenic with minimal gene rearrangements and a few gene losses. The topologies from different markers were largely congruent, confirming the relationships within Balanophoraceae and resolving the positions of the newly studied *Balanophora* species as well as the origins of apomictic species/populations. We further predicted *in silico* the subcellular localization of *Balanophora* proteins and provided new evidence that *Balanophora* plastids are still involved in many non-photosynthetic pathways, such as the biosynthesis of amino acids, fatty acids, and heme.

**Figure 1.**
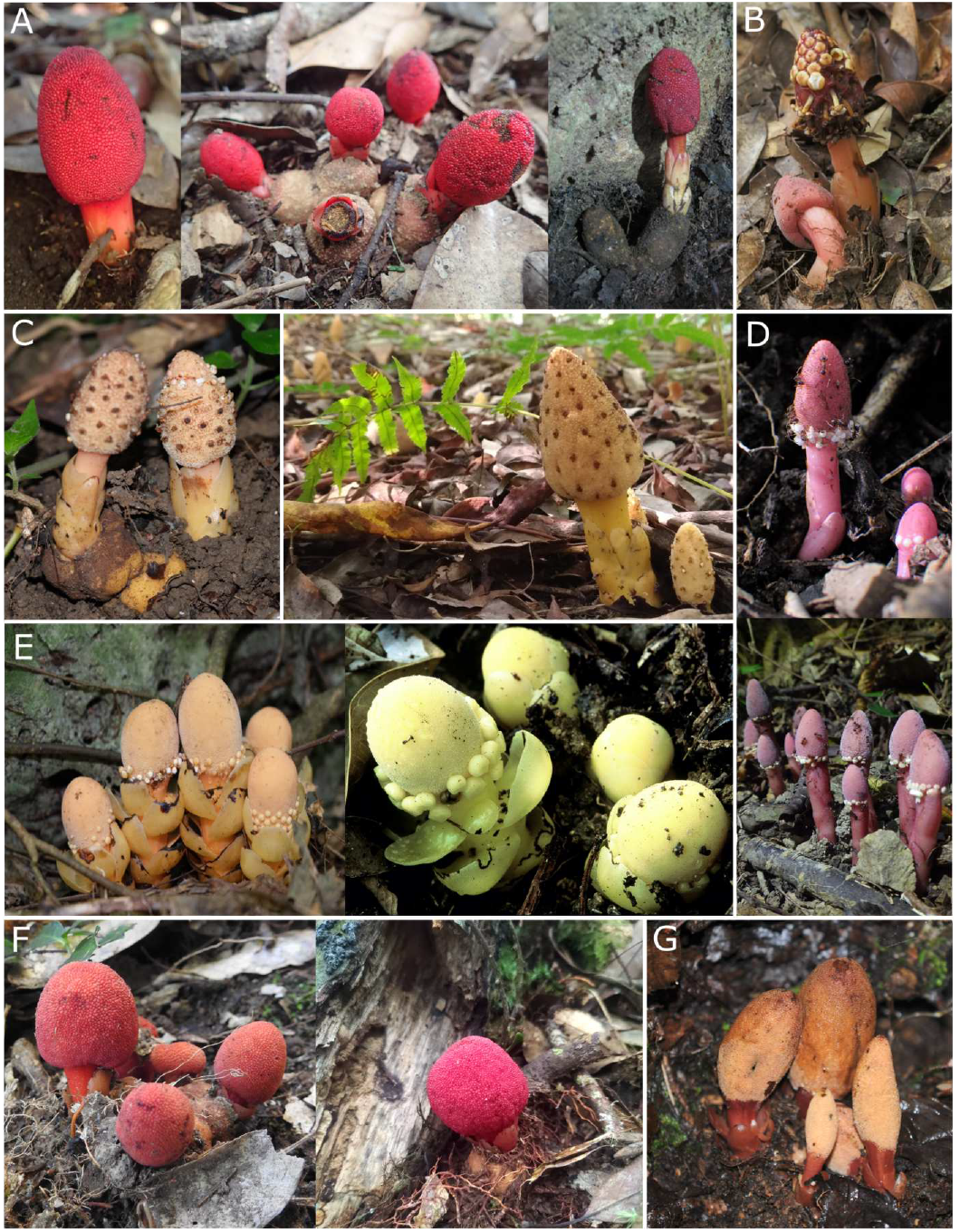
Sampled *Balanaphara* populations from Taiwan and Japan. **(A)** *B. japonica* (left and center: Kyushu, Japan; right: Taiwan), **(B)** *B. mutinoides* (Taiwan), **(C)** *B. tobiracola* (from left: Okinawa, Japan; Taiwan), **(D)** *B. subcupularis* (Kyushu, Japan), **(E)** *B. fungosa* ssp. *fungosa* (from left: Okinawa, Japan; Taiwan), **(F)** *B. yakushimensis* (from left: Kyushu, Japan; Taiwan), **(G)** *B. nipponica* (Honshu, Japan). For more information about sampling locations, see **Figure 2; Table S1**).

**Figure 2.**
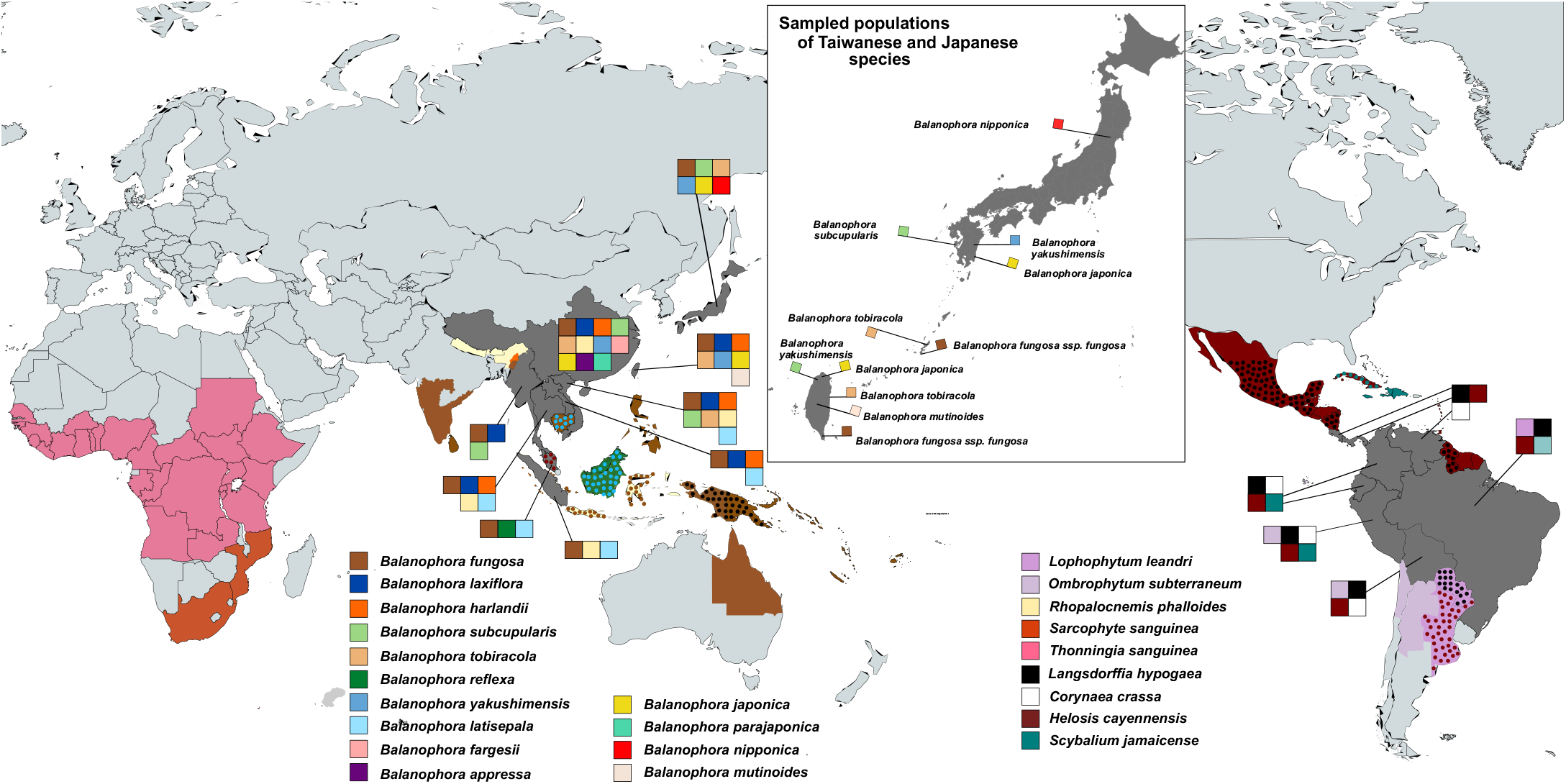
The world distribution of Balanophoraceae species with sequencing data available (and thus displayed on the phylogenetic trees in this study). Inset figure: sampling sites in Taiwan, Okinawa, and mainland Japan. Color coding is identical to **Figures 3, 4, and 5**. When only 2 species are presented in an area, the combination of small filled dots and background color was used.

## Results

### Plastid genomes of Japanese and Taiwanese *Balanaphara* species

The complete plastid genomes of six Japanese and five Taiwanese *Balanophora* species assembled here range from 14,259 to 16,290 bp in size, with a GC content of 12-13% (Figure S2; Table S2). They encode only 11-13 ribosomal protein-coding genes (*rpl* and *rps*), 3-4 other protein-coding genes (16S, 23S, 4.5S, and 5S rRNA), and 1 tRNA gene (*tRNA-Glu/trnE*) (Tables S3 and S4).

We newly report the plastomes of five *Balanophora* species not studied previously: *B. fungosa* ssp. *fungosa, B. japonica, B. mutinoides, B. nipponica*, and *B. tobiracola*. The plastomes of *B. tobiracola*, with 16,255-16,290 bp and 16 protein-coding genes, are the largest within the genus to date (Figure S2; Tables S2 and S3). Interestingly, the plastid genome of *B. mutinoides* lacks the *rpl2*+*rps19* gene fusion observed in the closely related *B. harlandii*. Comparison of the plastid genomes of Japanese and Taiwanese populations showed high similarity in plastome size, GC content, and the number of genes, with a high level of sequence identity (Tables S5 and S6). The *ycf1* gene appears to remain functional in transmembrane import, as its transmembrane domain is still detectable (SupplMat2).

All studied *Balanophora* plastomes contain genes with in-frame TAG codon(s), corroborating that all of them use the alternative genetic code described by Su et al. (2019), with the TAG codon reassigned from a stop codon to tryptophan. Depending on the species, we detected 1-8 in-frame TAG codons in 9-12 protein-coding genes (60-80% of all protein-coding genes). The exact numbers and positions of in-frame TAG codons within the individual genes are listed in Table S4.

All the plastid genomes were additionally compared to *de novo* genome assemblies generated in SPAdes-3.15.5 (Prjibelski et al. 2020) to rule out misassemblies resulting from the seed-based NOVOPlasty approach. The *de novo* plastome assemblies by SPAdes-3.15.5 and seed-based assemblies by NOVOPLAsty were identical for 5 out of 12 samples (Table S7). The remaining SPAdes assemblies were either not 100% identical with the NOVOPlasty assemblies or were identical but highly fragmented and incomplete. A short sequence comprising *rrn5* and the start of *ycf1* were missing from three SPAdes assemblies. These discrepancies can be caused by the differences in sequencing coverage affecting the assemblers in different ways but also by the presence of several versions of plastid genomes that mainly differ in highly variable regions (heteroplasmy).

### Phylogeny of the *Balanaphara* genus inferred from plastid genes

Analyses of all plastid-gene datasets (16S rRNA, 23S rRNA, 16S+23S rRNA, and 15 plastid protein-coding genes) using maximum likelihood (ML) and Bayesian inference methods yielded consistent topologies with four major clades of *Balanophora* (SupplMat1). The first clade contains *B. japonica, B. yakushimensis, B. nipponica*, and *B. laxiflora*. It is sister to clade 2, which comprises *B. fungosa* and *B. subcupularis*. These clades are sister to the third clade which includes *B. reflexa* and *B. latisepala*. The last clade contains species with the largest plastid genomes: *B. harlandii, B. mutinoides, B. fargesii*, and *B. tobiracola* (Figure 3).

**Figure 3.**
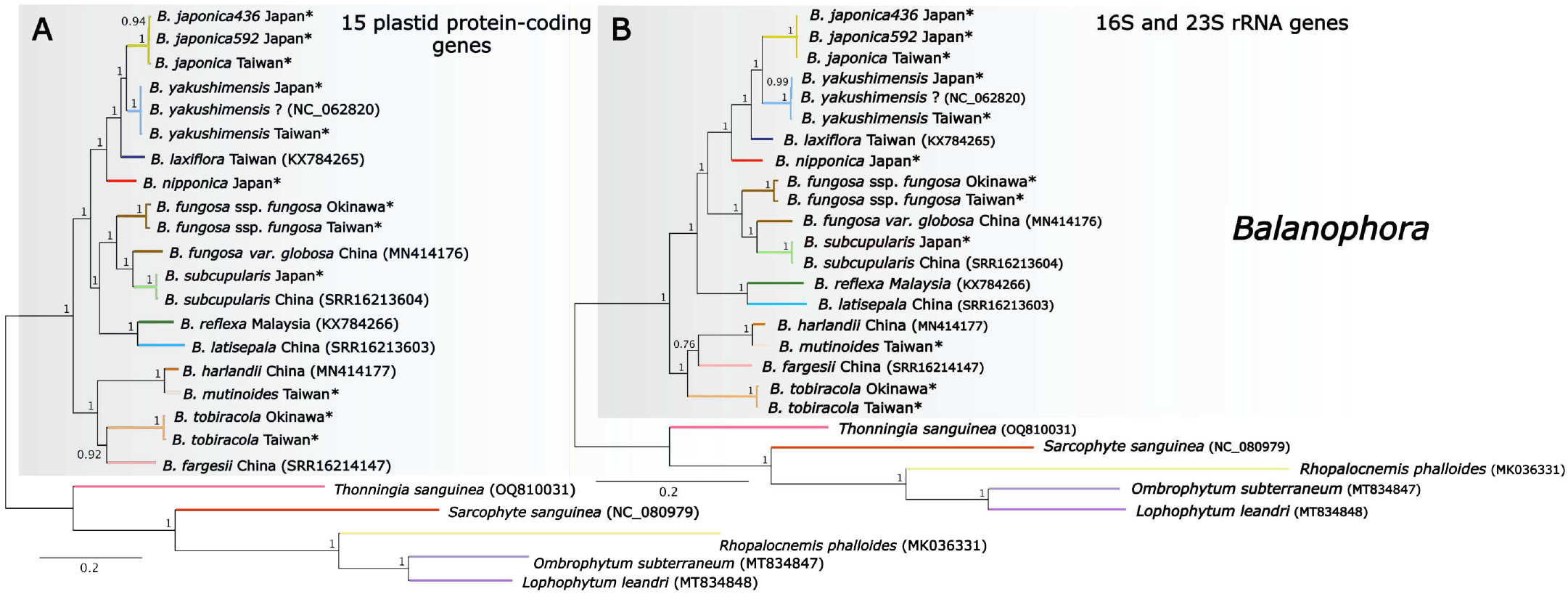
Phylogeny of Balanophoraceae inferred from plastid genes including all studied Taiwanese and Japanese *Balanaphara* (highlighted by asterisks). **(A)** Bayesian phylogeny inferred from 15 protein-coding plastid genes (LG+G model; 10,000,000 generations); **(B)** Bayesian phylogeny inferred from concatenated plastid 16S and 23S rRNA genes (GTR+G model; 10,000,000 generations). Posterior probabilities are displayed next to branches. Both alignments were trimmed by Gblocks before constructing the trees. Both phylogenetic trees are rooted. Protein-coding genes used: *accD, clpP, rpl2, rpl14, rpl36, rps2, rps3, rps4, rps7, rps11, rps12, rps14, rps18, rps19*, and *ycf1*. GenBank accession numbers for sequences obtained from NCBI are indicated in parentheses. The color coding is identical to the species distribution map in **Figure 2**.

All plastid-gene phylogenies (Figure 3; SupplMatlA-G and K-P) recovered a clade with *B. japonica* as sister to *B. yakushimensis*, with *B. laxiflora* as their sister, and with early divergence of *B. nipponica* within clade 1. Surprisingly, *B. subcupularis* is, with strong support, deeply nested in clade 2 and likely very closely related to *B. fungosa* ssp. *indica. B. reflexa* and *B. latisepala* were always clustered together in the same clade in all phylogenetic trees (Figure 3; SupplMatlA-G and K-P).

In the last clade, *B. harlandii* and *B. mutinoides* form a sister subclade to *B. tobiracola* and likely *B. fargesii*, although the position of *B. fargesii* is not stable based on 16S+23S rRNA datasets (Figure 3B; SupplMat1E-G). In agreement with their plastome sequence identities, Japanese and Taiwanese populations are inferred to be well-supported sister branches (Figure 3).

### Plastid gene loss across *Ba/anaphara*

*Balanophora* species have completely syntenic plastomes with minimal gene rearrangements and a few gene losses (Figure 4; Table S3). All known *Balanophora* plastomes, including those revealed here, lack *rpl16, rps8, matK*, and other tRNAs when compared to outgroups. Moreover, several independent gene losses were found within the genus (Figure 4 left). The *ycf2* gene has been lost once from all species in clade 2 (e.g. *B. fungosa* and B. *subcupularis*) and twice independently in clade 1, from *B. nipponica* and *B. yakushimensis*. The *rpl36* gene is missing from clades 1 and 3. In addition, *rpl14* was lost during the diversification of the *B. harlandii* and *B. mutinoides* branches. The only gene rearrangement observed in *Balanophora* plastomes is the inversion of *rps4* in *B. harlandii* reported by Kim et al. (2023).

**Figure 4.**
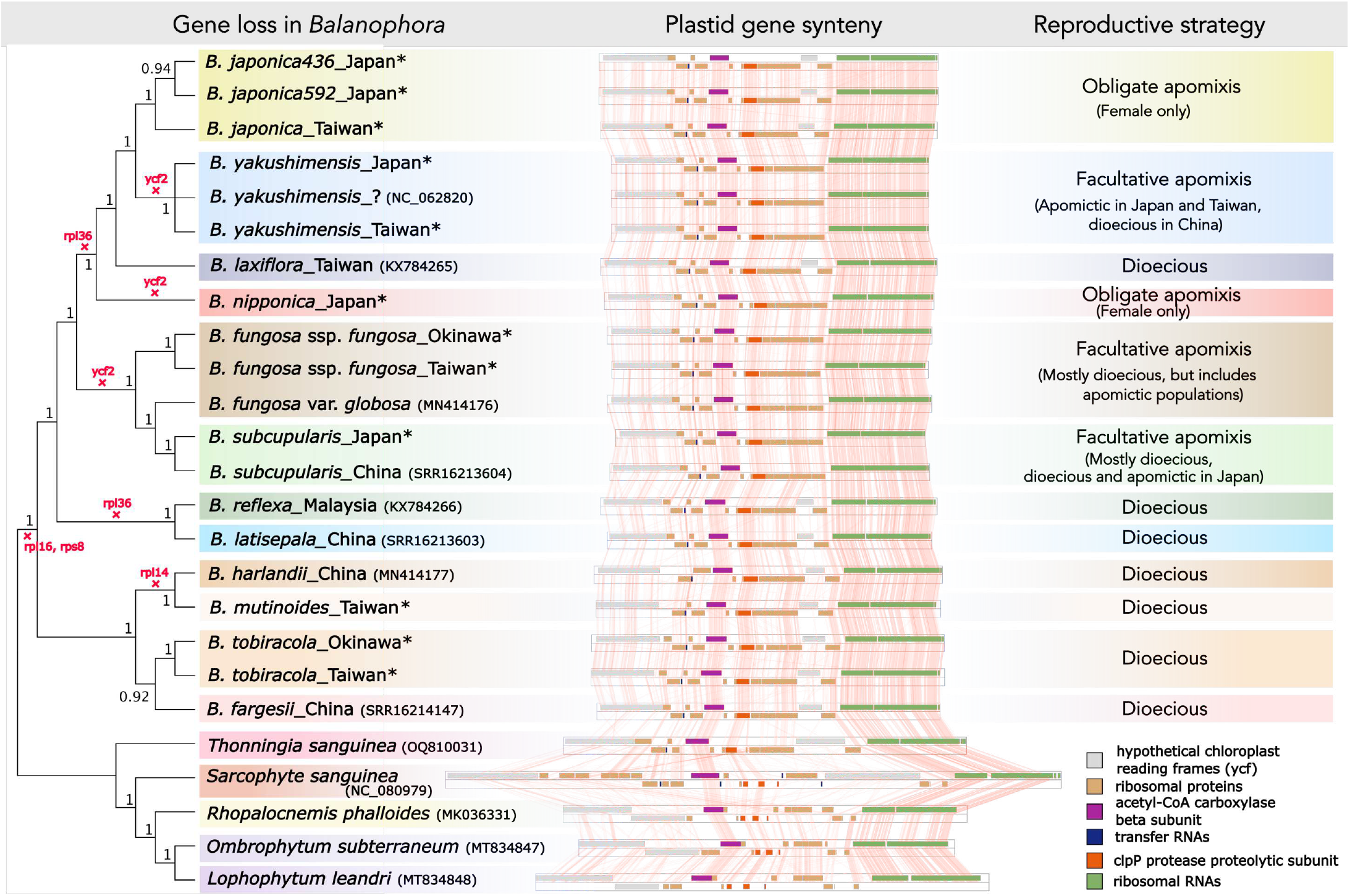
tBlastX alignment of whole plastid genomes of Balanophoraceae ordered according to their phylogeny and annotated with *Balanaphara* reproductive strategies. The cladogram was inferred from the Bayesian phylogenetic tree based on 15 protein-coding plastid genes (LG+G model; 10,000,000 generations; **Figure 3A**) and posterior probabilities are displayed next to nodes. Independent losses of specific plastid genes are highlighted in red. GenBank accession numbers for sequences obtained from NCBI are indicated in parentheses. The color coding is identical to the species distribution map in **Figure 2**.

### Phylogeny of the genus *Balanaphara* inferred from nuclear genes

The positions of most *Balanophora* species within the Balanophoraceae phylogenetic tree showed high congruence between the datasets and inference methods used (Figure 3 and 5; SupplMat1). The phylogeny based on 18S rRNA revealed relationships within *Balanophora* similar to those based on plastid genes, although the position and topology of the first clade could not be resolved-it is either sister to clade 2 or clade 3 (Figure 5A; SupplMatlH-J).

**Figure 5.**
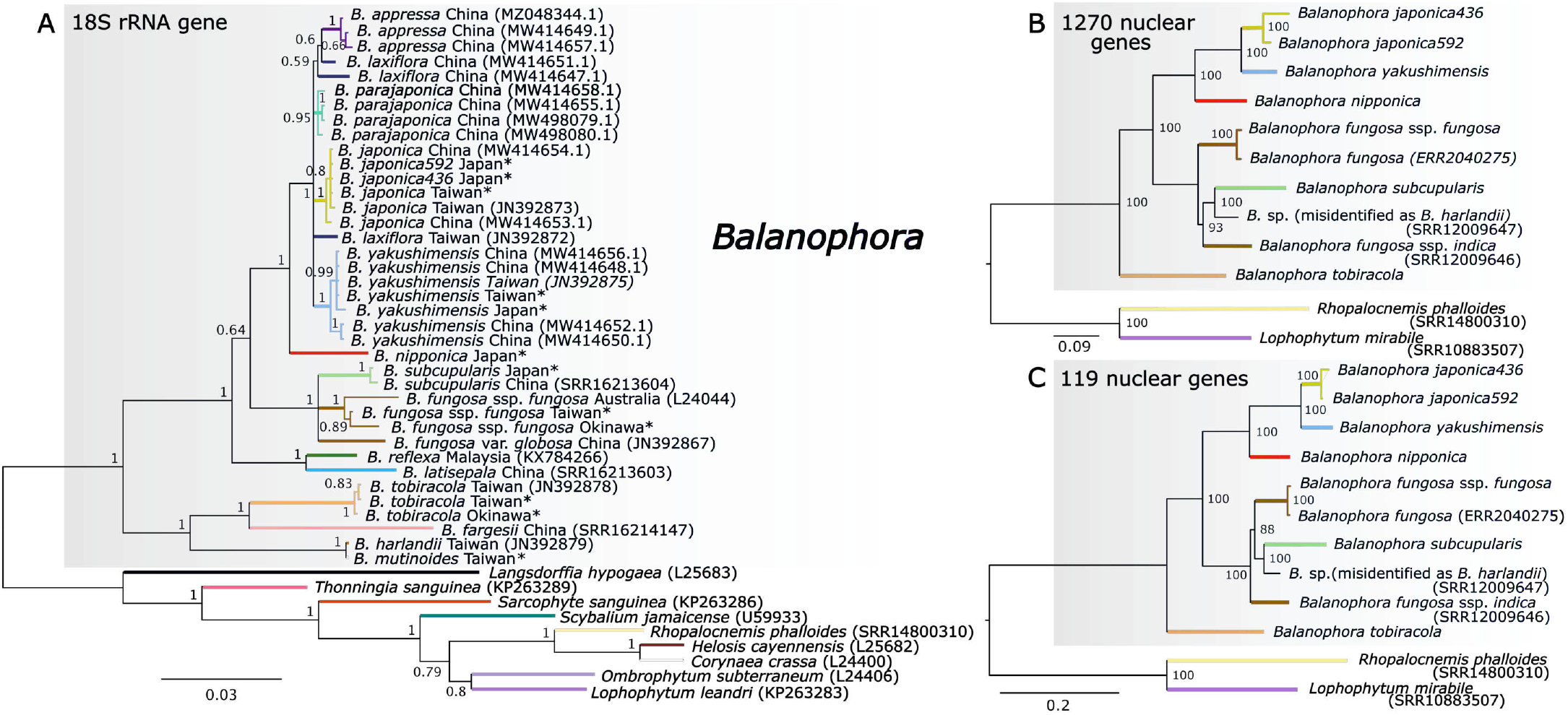
Phylogeny of Balanophoraceae inferred from nuclear genes. **(A)** Bayesian phylogeny inferred from the nuclear 18S rRNA gene (GTR+G model; 10,000,000 generations); **(B)** A consensus species tree inferred by IQTREE (1,000 bootstraps; JTT+F+R5) from 1,270 nuclear genes predicted from species transcriptomes and extracted by the STAG (Species Tree Inference for All Genes) method implemented in Orthofinder. **(C)** Maximum likelihood tree inferred by IQTREE (1,000 bootstraps; JTT+F+R3) from 119 single-copy nuclear genes predicted from species transcriptomes and recovered from at least 8 species. Publicly available transcriptomic data are accompanied by accession numbers. The color coding is identical to the species distribution map in **Figure 2**.

Multigene phylogenies inferred from nuclear transcriptomes (Figures 5B and 5C) confirmed well-resolved positions of closely related *B. japonica, B. yakushimensis, and B. nipponica* within clade 1 and their sister clade 2, including *B. fungosa* ssp. *fungosa, B. fungosa* ssp. *indica*, and *B. subcupularis*. However, we were not able to fully resolve the relationships within clade 2, as the topologies differed depending on the dataset used. In agreement with previous analyses, *B. tobiracola* is sister to all other species from the genus for which transcriptomic data are currently available. We also determined that the publicly available transcriptomic data referred to as *B. harlandii* (SRR12009647) were likely misidentified and instead derived from a sample belonging to the *B. fungosa* clade (Figure S3).

### Transcriptome assemblies

Our transcriptome assemblies of *Balanophora* showed a slightly lower level of completeness compared to the published *Balanophora* genomes (Figure 6; Table S8). However, they were still sufficiently complete for the extraction of phylogenetic markers. Out of 1,614 conserved Embryophyta genes, we detected 516-691 complete and single-copy genes from the transcriptomes and more than 800 genes from the high-quality genomes. The analyses revealed many (about 300-400) complete and duplicated genes in several transcriptomes (e.g., *B. japonica592, B. subcupularis*, and *B. tobiracola*; see Figure 6). These duplicates likely represent transcript isoforms, paralogous copies of these genes in some *Balanophora* genomes, or RNA contamination from the host. Only single-copy genes were used for downstream analyses, and the single-gene alignments were manually inspected to reduce the effects of these issues.

**Figure 6.**
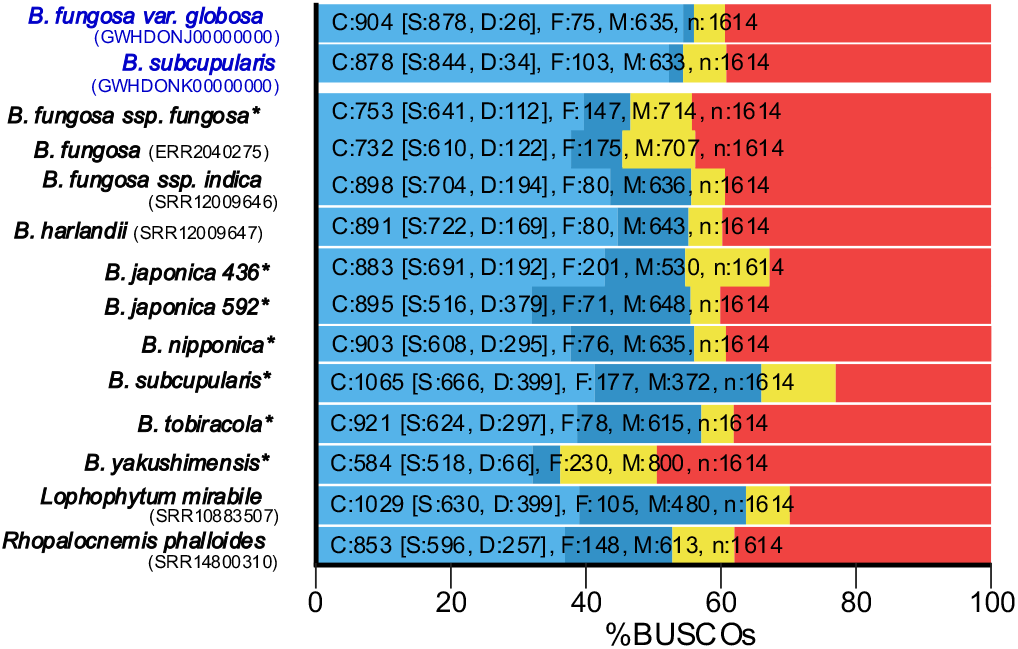
BUSCO completeness of transcriptomes generated in this study (highlighted by asterisks) and re-assembled from publicly available transcriptomic data (accession numbers in parentheses). Two high-quality *Balanophora* genomes were included as a reference in the analysis (in blue), *B. fungosa* var. *globosa* (GWHDONJ00000000) and *B. subcupularis* (GWHDONK00000000) published by Chen et al. (2023). C: complete transcripts including single-copy (S) and duplicated (D) transcripts; F: fragmented transcripts; M: missing transcripts.

### Plastid metabolic pathways predicted from transcriptomic data

We found a surprisingly high number of proteins predicted *in silico* to be targeted to the Balanophoraceae plastids from the host cytoplasm, ranging from 897 to 3,134 (1,794 on average; Table S9). When we clustered these protein sequences at 98% and 95% sequence identities, 5-24% (16% on average) and 8-37% (20% on average) of the sequences were identified, likely respresenting protein isoforms or recent paralogs. According to EggNOG results (Figures 7, S4, and S5), the proteins are involved in the essential and non-essential amino acid biosynthesis, fatty acid biosynthesis, riboflavin, lipoic acid, heme, and isoprenoid biosynthesis (MEP), glycolysis, and pentose phosphate pathway. The completeness of the pathways is, in general, similar across species. When counting only proteins with high-confidence plastid translocons to exclude false positives or dual-targeted proteins (e.g. mitochondrial proteins), this conservative subset contains 366 to 1,457 proteins (805 on average; Table S9).

**Figure 7.**
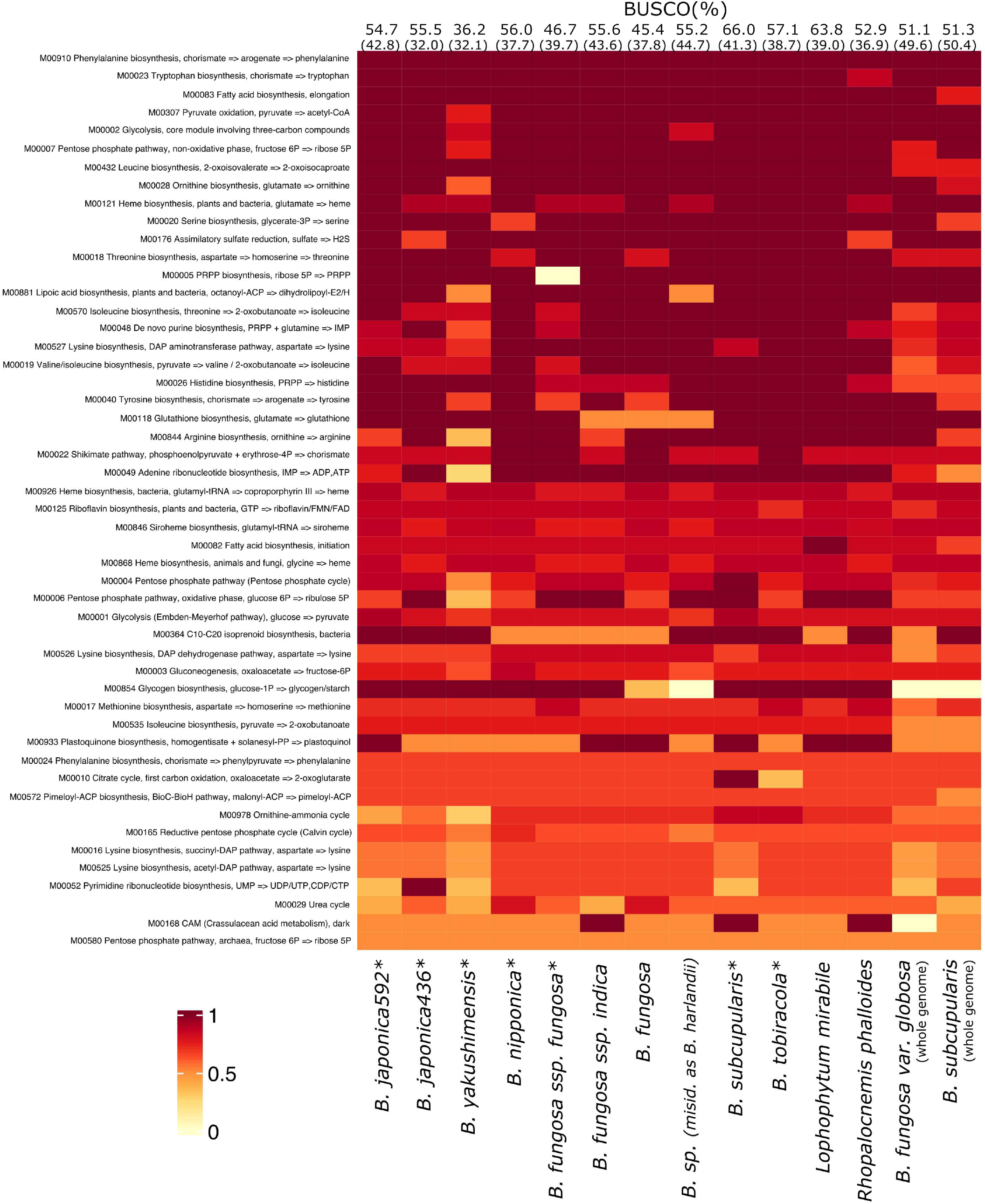
Completeness of 50 most complete plastid biosynthetic pathways inferred from predicted proteomes of both newly generated data (highlighted by asterisks; Table S8) and previously published data. (*B. fungosa spp. indica* (SRR12009646), *B. fungosa* (ERR2040275), *B*. sp. (misidentified as *B. harlandii*; SRR12009647), *L. mirabile* (SRR10883507), *R. phalloides* (SRR14800310), *B. fungosa var. globosa* (GWHDONJ00000000), and *B. subcupularis* (GWHDONK00000000)). *B. fungosa var. globosa* and *B. subcupularis* show completeness of plastid biosynthetic pathways inferred from proteomes predicted from whole genome data. BUSCO completeness scores **(Figure 5)** of the transcriptomes are indicated in percentages above the figure. Complete transcripts including single-copy and duplicated transcripts and only single-copy transcripts (in parenthesis) are displayed. The samples are ordered according to their phylogeny.

## Discussion

### Phylogenomics confirms the relationships within Balanophoraceae and highlights the importance of taxon sampling for taxonomically challenging species

The relationships among the genera of Balanophoraceae generally agree with the recently published phylogenies of the family based on the diverse datasets containing no more than 22 concatenated genes (Su et al. 2015; Kim et al. 2023; Sanchez-Puerta et al. 2023). The analysis of nuclear *l8S rRNA* (Figure S6) confirmed four subfamilies within the family as suggested by Sanchez-Puerta et al. (2023). Balanophoroideae comprising *Balanophora* + *Langsdorffia* + *Thonningia*, is sister to a clade comprising three other subfamilies: Scybalioideae (with *Scybalium*), Lophophytoideae (with *Lophophytum* + *Ombrophytum*), and Helosidoideae (with *Rhopalocnesmis* + *Helosis* + *Corynaea*). The relationships within these latter subfamilies are congruent with the previous molecular phylogenies. In agreement with the phylogeny of Sanchez-Puerta et al. (2023), which was inferred from the nuclear *rDNA* operon and mitochondrial *matR* sequences, the position of *Sarcophyte* could not be resolved with the nuclear *18S rRNA gene*. According to our results, *Sarcophyte* is either sister to *Thonningia* or falls within the Lophophytoideae-Helosidoideae clade. Its placement as sister to all other genera in Balanophoraceae, as suggested by Kim et al. (2023), may be an outgroup attraction caused by aligning protein-coding sequences in nucleotides (thereby disrupting the triplet codon structure) and inferring the phylogeny with nucleotide substitution models.

The relationships within the genus *Balanophora* recovered in this study generally agree with the published phylogenies (e.g., Su et al. 2012; Yu et al. 2022). Additionally, our findings contribute to a deeper understanding of the speciation and taxonomy of *Balanophora*. We provide the first molecular data on the only Japanese endemic *Balanophora, B. nipponica*, and reveal its phylogenetic position. *B. nipponica* is an early-diverging member of clade 1 (Figures 3, 5, and S6; also including *B. appressa, B. japonica, B. laxiflora, B. parajaponica*, and *B. yakushimensis*). *B. nipponica* was previously considered a synonym of *B. japonica* (Hansen 1972). However, our pairwise genetic distance analysis (Table S7) reveals a significantly greater divergence between *B. nipponica* and *B. japonica* than that observed within *B. japonica* populations. Furthermore, the genetic distance between *B. nipponica* and other species is notably distinct. These findings, together with morphological studies (Watanabe and Akuzawa 1982; Murata 1990), strongly support the recognition of *B. nipponica* as a separate species.

We also confirmed that *B. subcupularis* is closely related to *B. fungosa* as suggested previously by Yu et al. (2022). According to our multigene phylogenies, it is a sister species to *B. fungosa* var. *globosa* (Figures 3, 5A, and S6) and/or *B. fungosa* ssp. *indica* (Figures 5B and 5C). However, we could not fully resolve relationships within this clade, since no plastome data from *B. fungosa* ssp. *indica* and no transcriptomic data from *B. fungosa* var. *globosa* are currently available.

Our phylogenetic results and comparison of whole plastome sequences indicate that *B. fungosa* subspecies and *B. mutinoides* (now part of the *B. harlandii* complex) might represent different species rather than subspecies variation.

*B. fungosa* var. *globosa* shares higher nucleotide identity in its plastome with *B. subcupularis* than with other *B. fungosa* subspecies, *B. fungosa* ssp. *fungosa* (Tables S5 and S6). *B. fungosa* ssp. *indica* also appears quite distinct from *B. fungosa* ssp. *fungosa* based on our nuclear multigene phylogenies (Figures 5B and 5C). Another example is *B. mutinoides* from Taiwan, which has been previously proposed as a separate species within the *B. kawakamii* lineage in the *B. harlandii* complex (Yu et al. 2022). The genetic differences between *B. mutinoides* and *B. harlandii* observed here indeed exceed the range observed between populations (Tables S5 and S6). This new classification is supported by differences in their morphology (leaf arrangement along the stem) and phenology (Yu et al. 2022). Our multigene phylogeny not only corroborates the findings of previous studies (Su et al. 2012; Yu et al. 2022) in placing *B. japonica* separately from the *Balania* subgenus, but also reveals morphological classification discrepancies within the section *Polyplethia* of the *Balanophora* subgenus. Although both *B. laxiflora* and *B. latisepala* were assigned to section *Polyplethia*, both plastid and nuclear *18S rRNA* phylogenies suggest that *B. laxiflora* is more closely related to *B. japonica* and *B. yakushimensis*. Moreover, *B. latisepala* and *B. reflexa* were originally placed in different sections under the *Balanophora* subgenus (Hansen, 1972), yet they share common traits, including 4-5-merous male flowers, polyporate pollen, and dioecious reproduction.

Our transcriptomics phylogenies examining the evolution of Balanophoraceae confirms that trees inferred from hundreds of nuclear genes complement and validate phylogenies inferred from limited gene sets. We advocate for the broader use of phylotranscriptomics with even richer Balanophoraceae taxon sampling to better resolve the positions of taxa such as *Sarcophyte* and species from the *B. harlandii* and *B. laxiflora* clades that could not be resolved here due to the unavailability of data.

### The evolution of obligate apomixis potentially enables *Balanaphara* to colonize islands

The new phylogeny of Balanophoraceae based on the highest number of *Balanophora* species so far allowed us to trace the evolution of reproductive strategies in the genus (Figure 4 right). Most *Balanophora* species show dioecious sexual reproduction (Stebbins 1941; Hansen 1972; Su et al. 2012), indicating that dioecy is an ancestral trait and that apomixis is a later-acquired reproductive strategy in *Balanophora* (Yu et al. 2021). To date, only *B. japonica*, and *B. nipponica* have been reported as obligately apomictic (Murata 1990; Su et al. 2012). Other *Balanophora* species include obligately apomictic populations, such as *B. fungosa* ssp. *indica* var. *globosa* in Java, *B. fungosa* ssp. *indica* in India and the Indo-Chinese subcontinent, *B. elongata* var. *ungeriana* in Java, *B. elongata* in Sumatra and Java, and *B. yakushimensis* in Taiwan and Japan (Hansen 1972; Su et al. 2012; Yu et al. 2021). According to Yu et al. (2021), there are at least two independent origins of apomixis in *Balanophora*. Alternatively, the ability of *Balanophora* species with hermaphroditic flowers to set fruit via facultative apomixis may have been acquired much earlier in the evolution of *Balanophora*, potentially representing a single origin. The evidence that not only *B. yakushimensis* produces solely female individuals, but other *Balanophora* species (e.g., *B. subcupularis* and *B. tobiracola*) also exhibit apomixis (i.e., they reproduce via apomixis yet also engage in dioecious reproduction; Suetsugu and Hashiwaki 2024; Suetsugu 2025) corroborates this hypothesis (Figure 4 right).

The capacity for apomixis in *Balanophora* species with hermaphroditic flowers may have functioned as a preadaptation for the evolution of obligate apomixis in some species. Given that facultatively apomictic species already possess the capacity for apomictic reproduction, the genetic changes required to induce obligate apomixis may be relatively simple and might not necessitate loss of function in the sexual pathway (Van Der Kooi and Schwander, 2014). While obligate asexuality is rare in plants, several obligately apomictic taxa, such as *Corunastylis* and *Boechera*, appear to have transitioned from facultative to obligate apomixis (Sorensen et al. 2009; Aliyu et al. 2010). A similar pathway is plausible for *Balanophora* based on current evidence (e.g., Suetsugu and Hashiwaki 2024; Suetsugu 2025).

Based on the data available, obligate apomictic species likely originated once or twice in the evolution of *Balanophora*: first, in *B. nipponica* (or its ancestor) and second, in the ancestor of *B. japonica* in Japan and Taiwan. The second independent origin of obligate apomixis remains unclear. Based on the phylogeny and pairwise genetic distance data from Yu et al. (2021), *B. japonica* and *B. parajaponica* are extremely similar in morphology, and their genetic distance falls within the range observed among populations of, for example, *B. laxiflora* or *B. yakushimensis*. This indicates that *B. japonica* and *B. parajaponica* are potentially conspecific; if so, only *B. nipponica* is the sole obligate apomict at the species level.

What is clear is that *Balanophora* populations with obligate apomixis occur exclusively on islands, Taiwan and/or Japan. Both regions have been isolated from their respective land masses for millions of years, making them valuable for studying ecological gradients, plant distribution patterns, and species adaptation to different ecological niches. The Japanese archipelago, a major component of the continental islands in East Asia, has been proposed as a refugium harboring Tertiary relict floras (Milne and Abbott 2002). Its current biota have been shaped by a sequence of migrations, extinctions, and speciations in response to paleogeographical isolation and/or connections (Millien-Parra and Jaeger 1999). Balanophoraceae is one of the oldest parasitic lineages, originating in the Cretaceous ca. 100 Mya (Nauman et al. 2013). Japanese flora is rich in *Balanophora* species and marks the northernmost extent of their distribution. From an evolutionary perspective, these species exemplify an “out-of-the-tropics” diversification. However, the biogeography of the genus including their dispersal to or vicariance in Japan remains unexplained. The Ryukyu Arc may have acted as a stepping-stone chain allowing the dispersal of some species via land bridges from the continent through Taiwan to the Ryukyus during the Pleistocene (Osozawa et al. 2012) but the exact level of population connectivity of *Balanophora* across the Asia-Pacific remains to be investigated.

On the other hand, Taiwan is situated at the tectonic boundary between the Eurasian Plate and the Philippine Sea Plate. Its flora represents a transitional zone, exhibiting characteristics of both the Eastern Asiatic and Malesian regions, influenced by its unique geographical and historical context (Takhtajan 1986). Of the six *Balanophora* species currently recognized in Taiwan, two exhibit probable obligate apomixis, as only females have been observed (Su et al. 2012).

The capacity for apomixis may have contributed to the extended habitats of *Balanophora* species. This aligns with Baker’s law proposing increased occurrence of uniparental reproduction in habitats that are colonized through long-distance dispersal (Baker 1955; Keller et al. 2024). Uniparental reproduction enabled by apomixis allows populations to be established by single individuals, making apomictic species more effective colonizers than their sexually reproducing relatives (Hojsgaard and Hörandl 2015). Consequently, apomixis has been associated with colonization success in environments with low carrying capacities, such as high-latitude and high-altitude regions (Stebbins 1959; Richards 2003; Hörandl 2008).

### *Balanaphara* plastids contain highly reduced genomes, yet they are still highly metabolically active in most other pathways usually present in chloroplasts

The extent of plastome reduction in studied *Balanophora* species (Figures S2) agrees with previously assembled plastomes of *Balanophoraceae* (Schelkunov et al. 2019; Su et al. 2019; Cerrioti et al. 2021; Kim et al. 2023; Figure 4 and Tables S2 and S3), confirming that the extensive gene loss occurred in their common ancestor. *Balanophora* plastomes are completely syntenic, the smallest, and most gene-dense within the family, yet they retain more genes than the plastomes of *Lophophytum, Ombrophytum*, and *Rhopalocnemis* (Tables S2 and S3). We confirmed the presence of the 5S rRNA gene, previously suggested by Su et al. (2012) and Kim et al. (2023), located between the 4.5S rRNA and *ycf1* genes in all studied plastomes. Interestingly, two *Balanophora* clades (clade 1 with *B. nipponica* and *B. yakushimensis* and clade 2 with *B. fungosa*; Figure 4) have independently lost the *ycf2* gene, which was recently revealed as an indispensable part of the Ycf2-FtsHi chloroplast protein import motor (Liang et al. 2024a). This complex collaborates with the TIC (Translocon at the Inner Chloroplast membrane) complex on preprotein translocation into chloroplasts, and its structure is conserved among many land plants (Liang et al. 2024b). As *ycf2* is missing in several *Balanophora* species and several mycoheterotrophic plants (e. g. Lam et al. 2015; Ravin et al. 2016), the exact composition and functioning of the protein import machinery should be studied in the future to determine whether the Ycf2-FtsHi and TIC complexes are present, partly modified, or lost (as shown for Poaceae; Liang et al. 2024b).

Loss of photosynthesis in primary plastids has occurred many times independently in angiosperms, but it is rare in other land plants (Hadariová et al. 2018). A similar extent of plastid reduction as observed in *Balanophora* can be found in unrelated lineages of full heterotrophs, including mycoheterotrophic and other parasitic plants (Table S3; Graham et al. 2017; Wicke and Naumann 2018). The plastomes of 12 mycoheterotrophic Thismiaceae are 14-18 kbp in size and contain a subset of genes similar to those in *Balanophora* (Garrett et al. 2023). Other examples of holoparasitic plants with convergent plastid reduction are found in the fully parasitic Hydnoraceae. The plastomes of five *Hydnora* and three *Prosopanche* species are almost twice as large as *Balanophora* plastomes, yet they share 19 genes out of their 23-26 unique genes (Jost et al. 2022). They still retain *rps8, rpl16*, and 2-5 additional tRNA genes missing from *Balanophora*, while the *clpP* (clp protease) gene, which is typically among the last to be lost during the final stages of plastid reduction (Hadariová et al. 2018), has already been erased.

The transition to heterotrophic plastids has also occurred multiple times in many unrelated protists with secondary and tertiary plastids, and the most extreme cases of plastid reduction are found in parasites (Hadariová et al. 2018; Mathur et al. 2019). Some of the most fascinating examples of species with reduced plastomes or complete plastid genome losses without or with organelle loss have been reported from Myzozoa (Janouškovec et al. 2015). As noted above, the plastomes of Balanophoraceae are exceptionally small and AT-biased (Table S2). The only comparable plastomes are two apicoplast genomes of the parasitic apicomplexan *Rhytidocystis* spp., which are about the same size as the *Balanophora* plastome and contain only 10 and 11 genes left with the lowest GC content (9 and 11.6%) reported to date (Mathur et al. 2021). Moreover, one of these species and other Myzozoa (Kwong et al. 2019; Mathur et al. 2019; Muñoz-Gómez et al. 2019) use an alternative genetic code where UGA (a stop codon in the standard genetic code) encodes tryptophan in their plastids, a feature also found in *Lophophytum* and *Ombrophytum* (Cerrioti et al. 2021).

Although establishing the exact number of plastid-targeted proteins is highly method and organism-dependent, chloroplasts of photosynthetic plants such as *Arabidopsis* contain approximately 3,000 proteins imported from the cytoplasm (Jarvis 2008; Rolland et al. 2012; Nakai 2018). In non-photosynthetic tissues of green plants (e.g., amyloplasts, chromoplasts, or root plastids), the number of imported proteins often varies based on the method used, but is usually reduced to slightly over 1,000 (Christian et al. 2023). The most reduced plastid proteomes in apicomplexan parasites or chlorophytes, such as *Helicosporidium* or *Polytomella*, still contain approximately 100-400 imported proteins (Pombert et al. 2014; Barylyuk et al. 2020; Fuentes-Ramírez et al. 2021) required for core functions such as Fe-S cluster assembly, and the biosynthesis of fatty acids and isoprenoids. Notable exceptions are *Cryptosporidium* (Apicomplexa) and *Hematodinium* (Dinoflagellata), which completely lack plastid organelles (Abrahamsen et al. 2004; Gornik et al. 2015). Direct proteomic data for heterotrophic plants are limited but it is hypothesized that as long as the plastid organelle is present, its nuclear-encoded proteome would range from several hundred to 1-2 thousand proteins. Among holoparasitic plants, the only proposed case of complete plastid genome loss is from *Rafflesia* spp., although the organelle remains visible in TEM micrographs (Molina et al. 2014). Even if we assume that *Rafflesia* contains the most reduced plastid proteome among plants, it likely relies on a similar number of nuclear-encoded proteins imported as in apicomplexans or chlorophytes to fulfill its core functions.

Our in silico protein targeting predictions with conservative filtering suggest that *Balanophora* plastid proteomes are not as functionally reduced and are more similar to plastids in non-photosynthetic tissues of photosynthetic plants. On average, we predicted over 800 nuclear-encoded plastid-targeted proteins across our *Balanophora* transcriptomes. We particularly focused on biosynthetic pathways, and the consistent presence of these specific genes across species (in agreement with BUSCO completeness) suggests a conserved role for the plastid among the *Balanophora* clade (Figure 7). Based on these results, the main metabolic pathways in the plastid include the biosynthesis of essential and non-essential amino acids, fatty acids, riboflavin, lipoic acid, heme, and isoprenoids, as well as glycolysis and the pentose phosphate cycle. However, since our data come from transcriptomes of only one tissue type (flowers that contain ‘chromoplasts’), we cannot rule out that the function of the non-photosynthetic *Balanophora* plastid is more differentiated or reduced when present in the tuber inside the host tree roots.

Interestingly, several pathways show consistently lower yet similar completeness (Figure 7), while some proteins contain dual localization signals (Table S9), suggesting not only a division of some processes across multiple compartments but also functional redundancy in mitochondrial and plastid proteomes (Xu et al. 2013). We thus hypothesize that the non-photosynthetic plastid interacts tightly with the mitochondrion - a relationship that should be tested by subcellular imaging of *Balanophora* cells. Cup-shaped plastids were previously proposed to be present in *Balanophora* in the vicinity of a large number of morphologically diverse mitochondria-like organelles (Gedalovich-Shedletzky and Kuijt 1990). The retention of functionally multi-use proteins that are dual-targeted to both endosymbiotic organelles (Duchêne et al. 2005) was recently linked to photosynthetic activity (DeTar et al. 2024). However, such an analysis for *Balanophora* will depend on a careful reconstruction of the mitochondrial proteome and single-gene phylogenies, which are currently hampered by numerous genes of both foreign (HGTs) and unclear origin in the extremely complex mitogenomes and mitoproteomes of *Balanophoraceae* (Sanchez-Puerta et al. 2017; Roulet et al. 2020; Zhou et al. 2023). A detailed look at other plastid and mitochondrial maintenance proteins such as DNA and RNA polymerases, ribosomal proteins, tRNAs, aminoacyl-tRNA transferases, translation-related factors, or TIC/TOC and TIM/TOM complex proteins in non-photosynthetic *Balanophora* could in the future make it a useful model for the discussion on functional constraints driving organelle proteomes in heterotrophic plants.

## Materials and Methods

### Plant sampling, nucleic acid extractions, and sequencing

All *Balanophora* species were sampled from natural populations in mainland Japan, Okinawa, and Taiwan (Figures 1 and 2; Table S1). The map of sampled populations and distribution maps of Balanophoraceae (Figure 2) and *Balanophora* (Figure S1) were created online with mapchart.net. Young intact above-ground tissue, including inflorescence and bracts, was collected and stored at −70°C. DNA and RNA were extracted from the inflorescence using the NucleoBond HMW DNA (MACHEREY-NAGEL, Duren, Germany) and MasterPure Complete DNA & RNA Purification (LGC Biosearch Technologies, Teddington, UK) kits. Prior to enzymatic lysis, the plant tissue was ground with liquid nitrogen. DNA and RNA concentrations and purity were determined using the Qubit 4 Fluorometer and NanoDrop One (Thermo Fisher Scientific, Waltham, MA, USA). PCR-free DNA and cDNA sequencing libraries were prepared with the NEBNext Ultra II DNA and NEBNext Ultra II RNA (NEB) Library Preparation kits for Illumina. Sequencing with 150 bp paired-end (PE) reads was carried out at the OIST SQC facility on the Illumina NovaSeq6000 and NovaSeqX platforms, and with 300 bp PE reads on the Illumina MiSeq sequencer at VYM Genome Research Center, National Yang-Ming University, Taiwan (Tables S7 and 8). The quality of raw sequencing data was assessed by FastQC version 0.12.1 and an initial taxonomic assessment was performed using phyloFlash v3.4.2 (Gruber-Vodicka et al. 2020).

### Plastid genome assemblies

Plastid genomes were assembled by NOVOPlasty v4.3.1 (Dierckxsens et al. 2016) and annotated in Prokka v1.14.6 (Seeman 2014). The genome annotations were further manually refined with NCBI BLASTP v2.14.1 (Altschul et al. 1990) searches against NR (non-redundant protein database) in Artemis v18.1.0 (Carver et al. 2012) and Geneious Prime 2023.2.1 (Kearse et al. 2012). The locations of ribosomal RNA and transfer RNA genes were predicted by RNAmmer 1.2 (Lagesen et al. 2007) and tRNAscan-SE (Chan and Lowe 2019), and confirmed via genome alignments with known *Balanophoraceae* plastomes. The ycf1 gene has been drastically reduced in size and was proposed to be involved in transmembrane import (de Vries et al. 2015). We thus checked its N-terminal transmembrane domain region with TMHMM (https://services.healthtech.dtu.dk/services/TMHMM-2.0/). The pairwise genetic distances between *Balanophora* plastid genomes were analyzed in MEGA 11 (Tamura et al. 2021) using the Jukes-Cantor model (Jukes and Cantor 1969) with 4 categories of nucleotide substitution rates. Finally, the circular plastid genomes were visualized by OGDraw (Greiner et al. 2019). All the plastid genomes were additionally compared to *de novo* genome assemblies generated in SPAdes-3.15.5 (Prjibelski et al. 2020) using the default settings to rule out misassemblies resulting from the seed-based NOVOPlasty approach.

### Assembly of 18S rRNA nuclear gene sequences

The full-length 18S rRNA nuclear gene sequence was directly assembled from raw reads using the phyloFlash v3.4.2 pipeline (Gruber-Vodicka et al. 2020). For two Japanese samples, *B. yakushimensis* and *B. fungosa* ssp. *fungosa*, this approach did not result in full-length sequences. Instead, we localized the contigs of the 18S rRNA gene in the draft genome assemblies using Barrnap v0.9 (Seemann 2022) and NCBI BLAST v2.14.1 (Altschul et al. 1990), and assembled the gene for each species in Geneious Prime 2023.2.1 (Kearse et al. 2012). Due to intraspecific variation, these two species contain several versions of the 18S rRNA gene that differ in highly variable regions. Only the most abundant variant was therefore selected for downstream analyses.

### Transcriptome assemblies

The transcriptomes of all studied Japanese species and other species from the Balanophoraceae family, for which transcriptomic data are publicly available in the NCBI SRA database (Sayers et al. 2022), were assembled using SPAdes-3.15.4 with the *-rna* option (Prjibelski et al. 2020). Transcriptome completeness was evaluated by BUSCO v5.7.1 (Manni et al. 2021) against the database of 1,614 conserved core genes from Embryophyta (embryophyta_odb10). The BUSCO summary results were visualized using the generate_plot.py BUSCO script. BUSCO scores were also calculated for the whole genomes of *B. fungosa* var. *globosa* and *B. subcupularis* (Chen et al. 2023), and these values were used as maximum completeness references. Proteomes were predicted from the transcriptomes using TransDecoder v5.5.0 [https://github.com/TransDecoder/TransDecoder].

### Phylogenetic analyses

In addition to phylotranscriptomics (see below), we analyzed five different datasets separately: plastid-encoded 16S rRNA, 23S rRNA, 16S+23S rRNA, and 15 protein-coding genes, plus the 18S rRNA nuclear gene dataset, using both maximum likelihood (ML) and Bayesian inference methods. The plastid protein-coding dataset comprised 12 genes of ribosomal proteins (*rpl2, rpl14, rpl36, rps2, rps3, rps4, rps7, rpsll, rpsl2, rpsl4, rpsl8*, and *rpsl9*) and three other genes (*ycf1, accD*, and *clpP*). The ribosomal RNA genes were aligned individually using the MAFFT v7.490 E-INS-i algorithm and concatenated in Geneious Prime 2023.2.1 (Kearse et al. 2012). We analyzed the rRNA datasets by IQ-TREE v1.6.12 (Nguyen et al. 2015) in ModelFinder mode and MrBayes v3.2.6 with the GTR+G model for 10,000,000 generations. The convergence of two independent runs and burn-in were checked in Geneious Prime 2023.2.1 (Kearse et al. 2012).

The protein-coding plastid genes were translated to amino acids and individually aligned using the MUSCLE v3.8.31 algorithm in SeaView v5.0.5 (Gouy et al. 2010). The alignments were inspected for in-frame stop codons (TAG and TGA) and re-assigned to tryptophan (W) according to their previously determined alternative genetic code (Su et al. 2019). The individual gene alignments were concatenated in Geneious Prime 2023.2.1 (Kearse et al. 2012). The concatenated protein-coding gene alignment was analyzed by IQ-TREE v1.6.12 (Nguyen et al. 2015) with a standard model selection method set to MFP (ModelFinder Plus) and by MrBayes v3.2.6 with the LG+G model for 10,000,000 generations. In addition, heterogeneous models LG+C20+F+G, LG+C40+F+G, and LG+C60+F+G were used in IQ-TREE v1.6.12.

Finally, we also inferred phylogenetic trees using the same settings as above from datasets trimmed by Gblocks v0.91b under a less stringent selection, allowing smaller final blocks, gap positions within the final blocks, and less strict flanking positions (Castresana 2000).

### Plastid genome alignment across the Balanophoraceae phylogeny

The whole plastid genomes of the species studied here, along with all other species from Balanophoraceae with available plastomes, were aligned and visualized using a custom script developed in Processing 4.3 (Reas and Fry 2006) [https://github.com/filip-husnik/genome-plots-processing]. Subsequently, the whole genome alignment was mapped on the topology inferred from the trimmed 15 plastid-encoded protein-coding gene alignment.

### Phylogenetic analyses based on transcriptomic data

To test the topologies of the phylogenetic trees derived from plastid gene datasets, we conducted additional phylogenetic analyses using nuclear genes from our transcriptomic data. The orthologous genes across all species were identified with OrthoFinder v2.5.4 (Emms and Kelly 2015). Multiple sequence alignments and gene trees for each orthogroup were inferred in OrthoFinder using the “*-M msa*” option. From the OrthoFinder results, we selected 119 single-copy orthologous genes present in at least eight out of twelve species and concatenated the alignments using Phyutility (Smith and Dunn 2008). These concatenated alignments were trimmed by trimAll v1.4.1 (Capella-Gutiérrez et al. 2009) using automatic settings. Additionally, we inferred a species tree from 1,270 nuclear genes using the STAG (Species Tree Inference for All Genes) method as implemented in OrthoFinder (Emms and Kelly 2018). Both datasets were analyzed using IQ-TREE v1.6.12 (Nguyen et al. 2015) with the model selection method set to MFP.

### Prediction of metabolic pathways in the plastid

Nuclear-encoded plastid-targeted proteins were predicted from the proteomes by Deeploc 2.0 (Thumuluri et al. 2022). Orthologous sequences were assigned and functionally annotated by a custom pipeline [https://github.com/ECBSU/genomics-scripts/tree/main/Functional-annotation/]. The scripts use eggNOG 5.0 (Huerta-Cepas et al. 2019) with the EggNOG MMseqs2 database for Viridiplantae, compare the outcome against standalone KEGG databases, and provide KEGG module completion and BRITE completion scores (reflecting the completeness of each biosynthetic pathway within the predicted plastid proteomes). To identify the number of protein isoforms and paralogs, all proteins predicted to be targeted to the plastid were also clustered in CD-HIT using 98% and 95% sequence identities as cut-offs (Li & Godzik 2006).

## Acknowledgments

We thank Hiromu Hashiwaki, Shuichi Kurogi, Hiroaki Yamashita, Akira Ichinowatari, and Yuki Ueno for assistance with plant sampling, and Arno Hagenbeek and Akito Shima for their help with data analyses. We are grateful for the help and support provided by the Sequencing Section (SQC) and Scientific Computing and Data Analysis (SCDA) sections of Core Facilities at OIST. We also thank the Environmental Science & Informatics Section of Core Facilities at OIST for obtaining the permissions required for plant sampling in Okinawa. The AI tools (ChatGPT and Grammarly) were only used for grammar corrections and minor text improvements. FH was supported by the JSPS KAKENHI grant 23K14256 and the HFSP Early Career Grant (RGEC29/2024). KS was financially supported by PRESTO (JPMJPR21D6) from the Japan Science and Technology Agency. HJS was supported by funding from the National Science and Technology Council, Taiwan (109-2311-B-845-001).

## Author Contributions

All authors designed and performed the research, contributed to the interpretation of the results, and wrote the manuscript. PS, HJS, and FH analyzed the data. HJS, KS, and FH supervised the project.

## Data Availability

Raw reads and plastid genome assemblies were deposited in the NCBI under the BioProject number (PRJNA1199576). Other data, such as plastome annotations and gene alignments, are available via Figshare (10.6084/m9.figshare.28588421).

## Supplementary Figures

**Figure S1.**
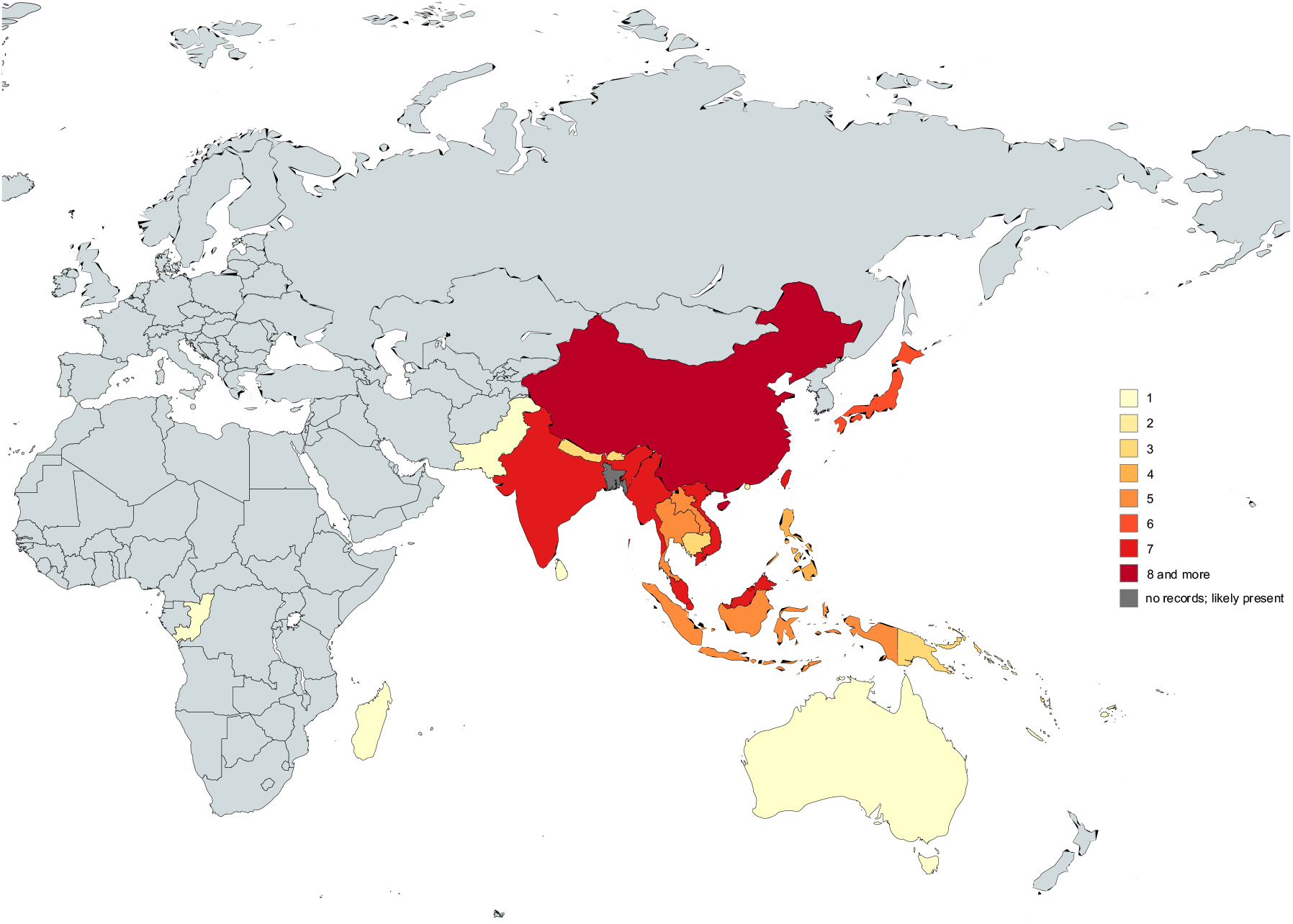
*Balanaphara* species diversity in the world. The number of species is indicated by a heatmap. Light gray corresponds to areas with no *Balanaphara* occurrence.

**Figure S2.**
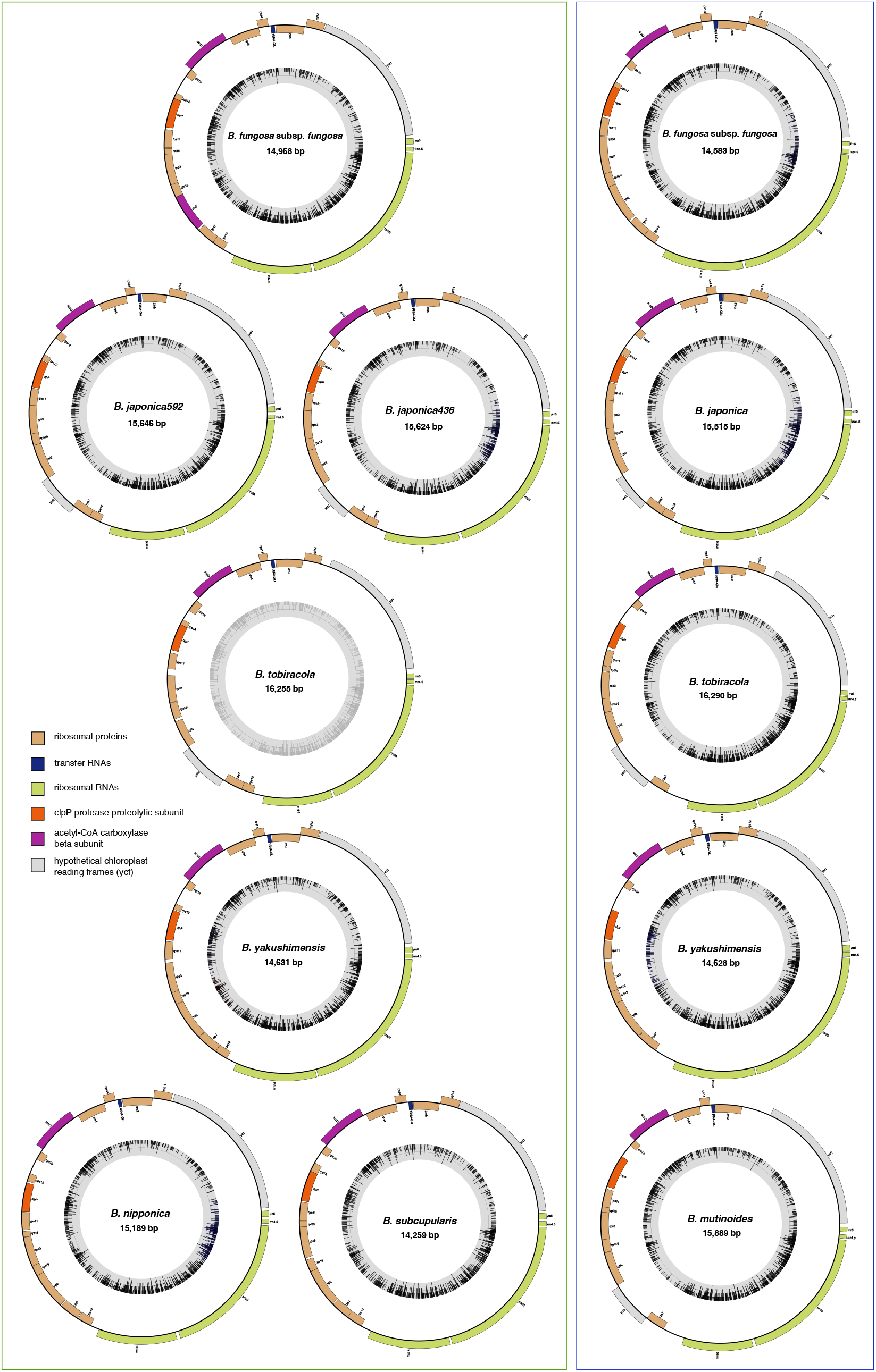
Circularized plastid genomes of the studied Japanese (left) and Taiwanese (right) *Balanaphara* species assembled in this study.

**Figure S3.**
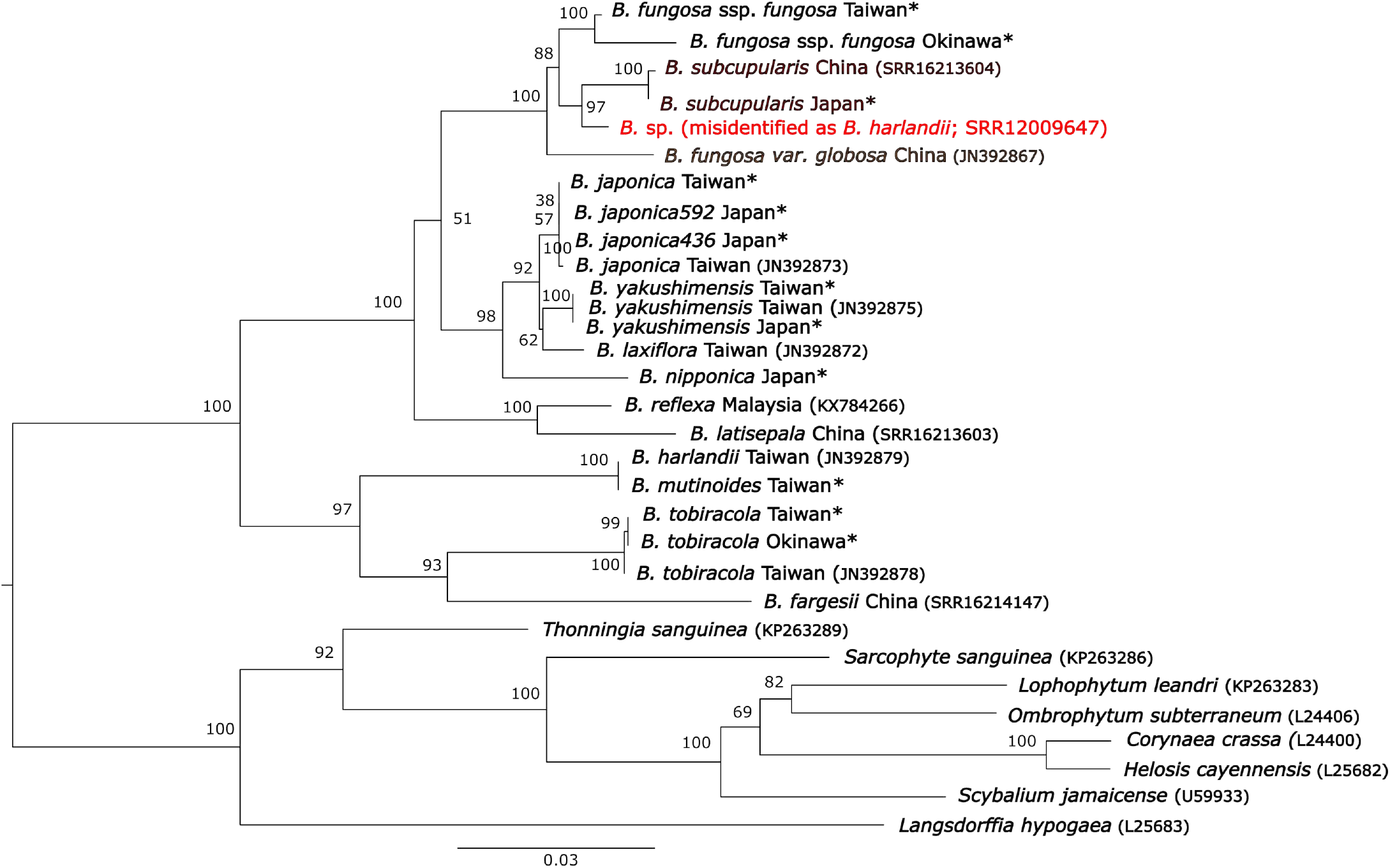
Maximum likelihood phylogeny of Balanophoraceae inferred by IQTREE from the nuclear 18S rRNA gene (GTR+F+I+G4 model; 1,000 bootstraps). The position of the misidentified *Balanaphara* from NCBI is in red. The phylogenetic tree is rooted, and bootstrap values are displayed next to branches. Sequences newly generated in this study are highlighted by asterisks. GenBank accession numbers for previously published public data are indicated in parentheses. The alignment was trimmed by Gblocks before constructing the tree.

**Figure S4.**
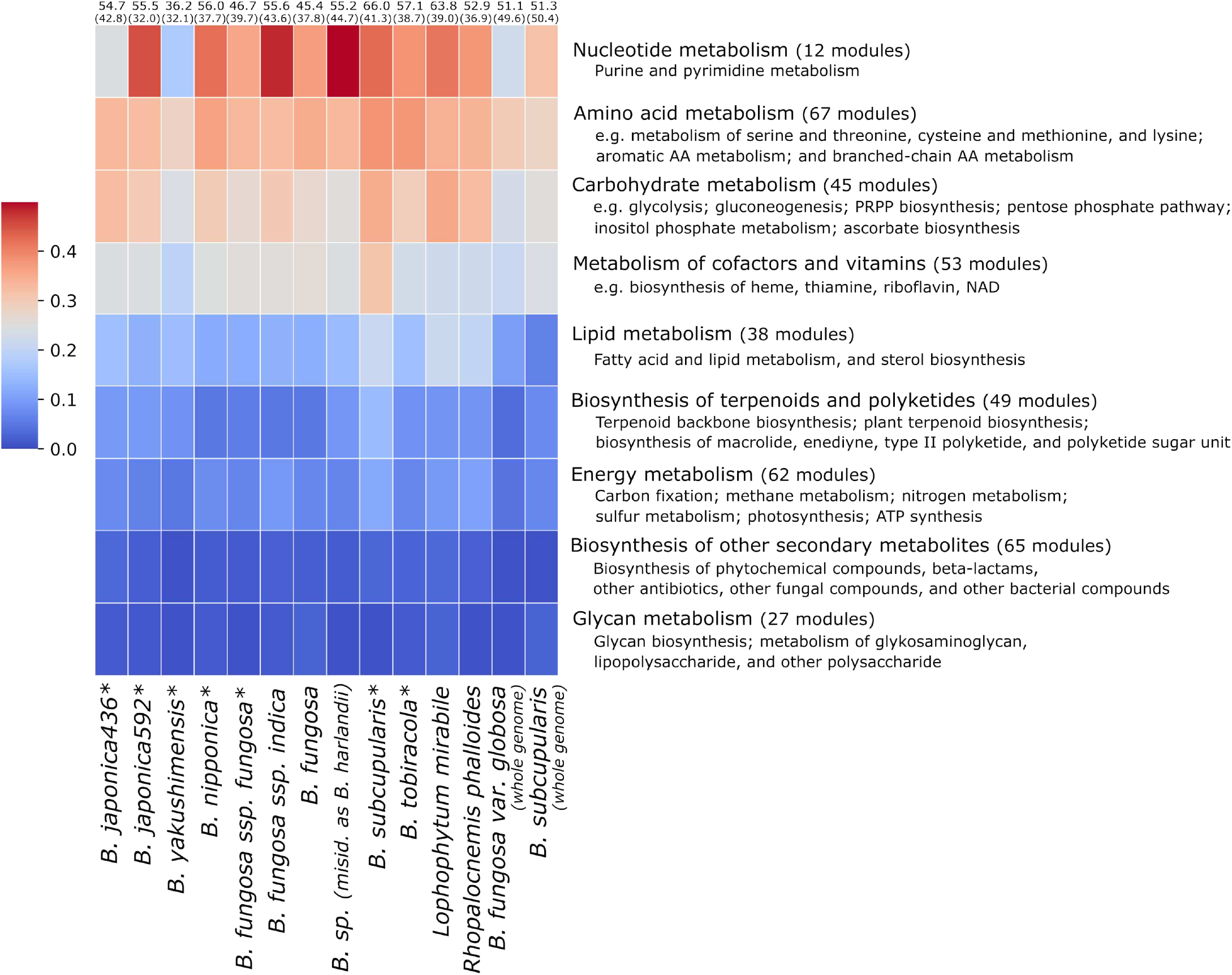
Average completeness of all plastid biosynthetic pathways from nine KEGG pathway-module categories. The plot was inferred from predicted proteomes of both newly generated data (highlighted by asterisks) and previously published data (*B. fungasa spp. indica* (SRR12009646), *B. fungasa* (ERR2040275), *B*. sp. (misidentified as *B. har/andii*; SRR12009647), *L. mirabi/e* (SRR10883507), *R. pha//aides* (SRR14800310), *B. fungasa var. g/abasa* (GWHDONJ00000000), and *B. subcupu/aris* (GWHDONK00000000)). *B. fungasa var. g/abasa* and *B. subcupu/aris* show completeness of plastid biosynthetic pathways inferred from proteomes predicted from whole genome data. For all KEGG modules and their completeness see Figure S5. BUSCO completeness scores of the transcriptomes are indicated as percentages above the figure. Complete transcripts including single-copy and duplicated transcripts and only single-copy transcripts (in parentheses) are displayed. The samples are ordered according to their phylogeny.

**Figure S5.**
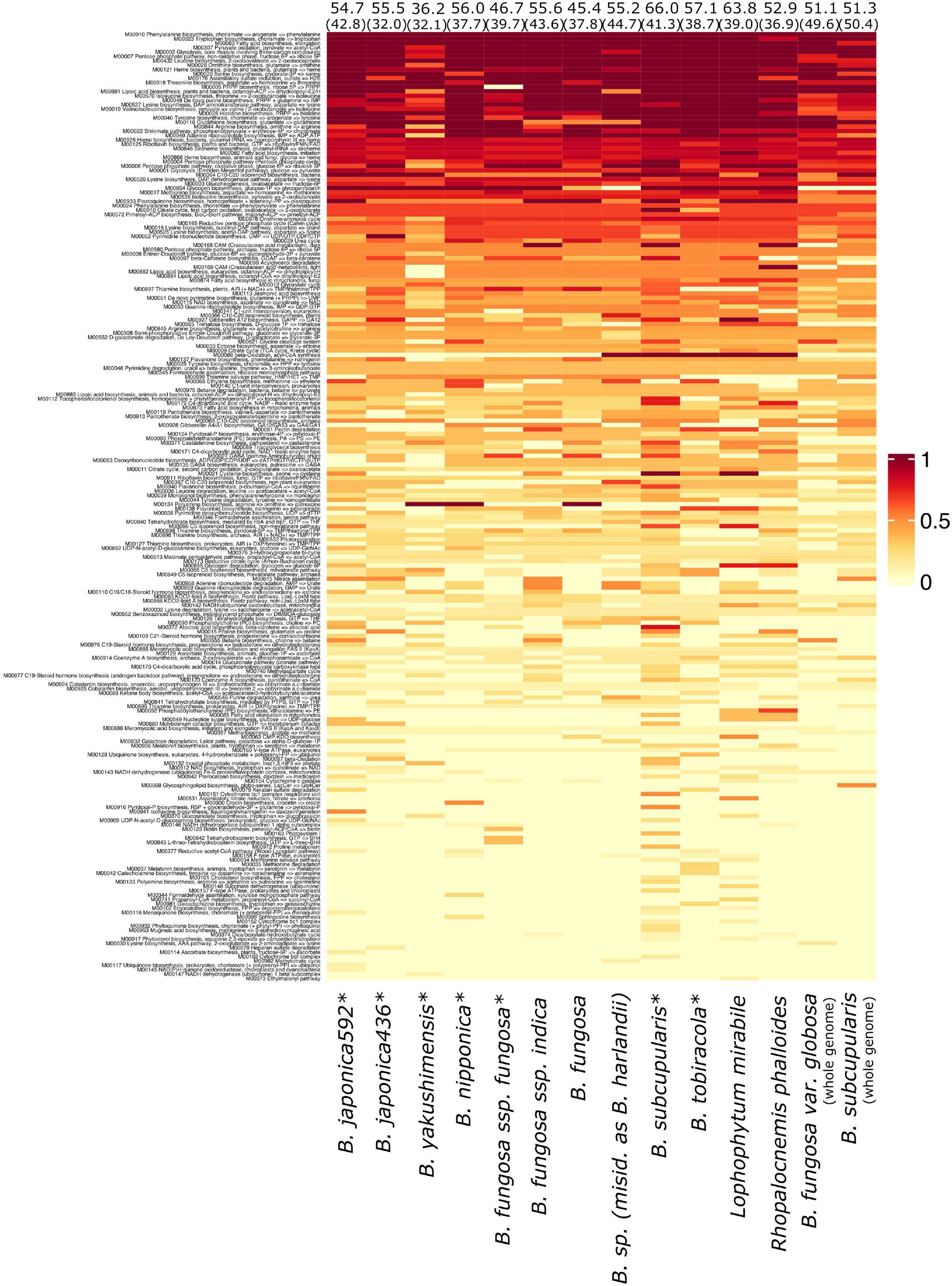
Completeness of all plastid biosynthetic pathways inferred from predicted proteomes of both newly generated data (highlighted by asterisks) and previously published data. (*B. fungasa spp. indica* (SRR12009646), *B. fungasa* (ERR2040275), *B*. sp. (misidentified as *B. har/andii*; SRR12009647), *L. mirabi/e* (SRR10883507), *R. pha//aides* (SRR14800310), *B. fungasa var. g/abasa* (GWHDONJ00000000), and *B. subcupu/aris* (GWHDONK00000000)). *B. fungasa var. g/abasa* and *B. subcupu/aris* show completeness of plastid biosynthetic pathways inferred from proteomes predicted from whole genome data. The samples are ordered according to their phylogeny.

**Figure S6.**
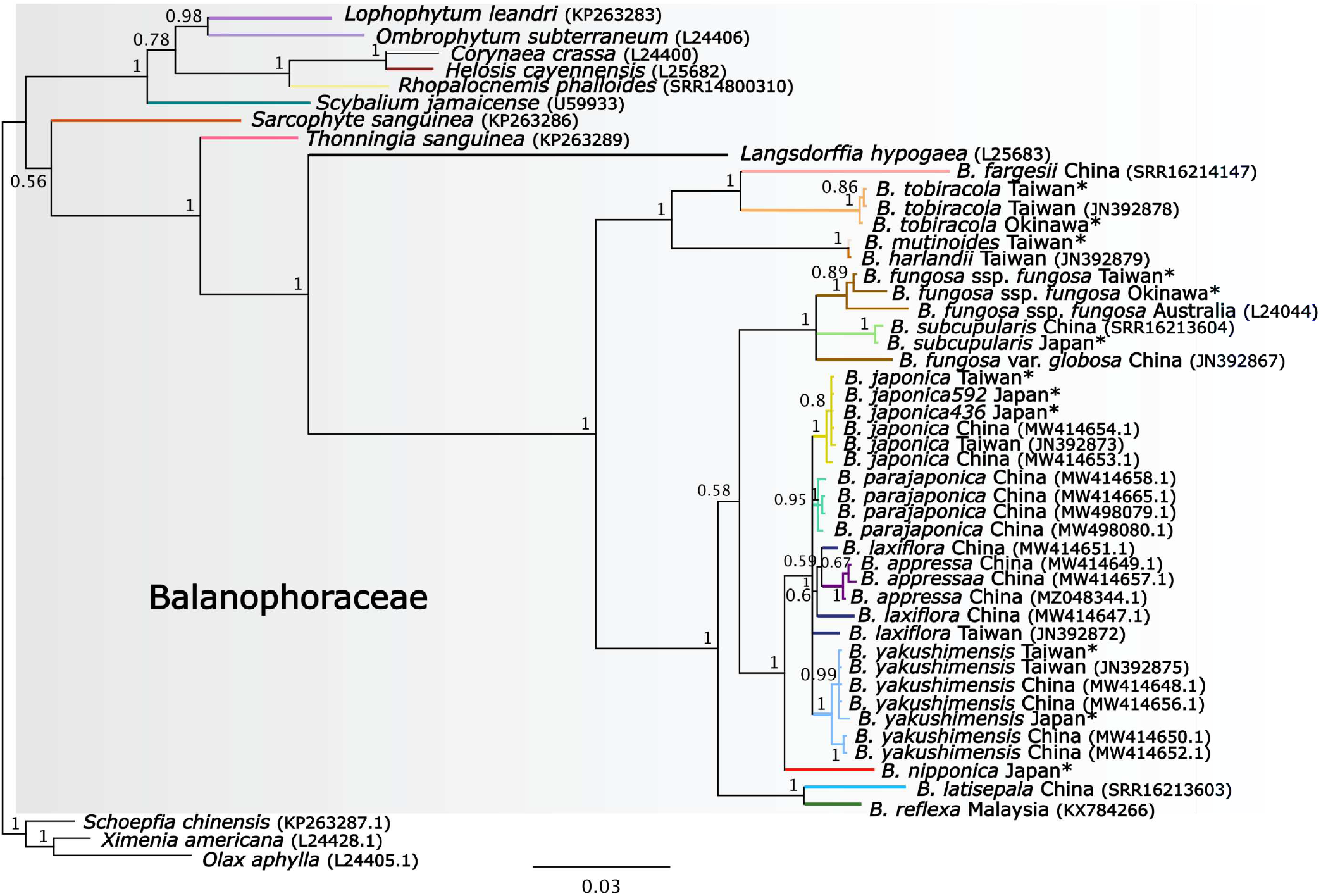
Bayesian phylogeny of Balanophoraceae inferred from the nuclear 18S rRNA gene (GTR+G model; 10,000,000 generations). Posterior probabilities are displayed next to branches. The dataset was trimmed by Gblocks before constructing the trees. The phylogenetic tree is rooted by three species from Santalales. GenBank accession numbers for sequences obtained from NCBI are indicated in parentheses.

