## Supplemental figures and tables for "Phylogenomics clarifies *Balanophora* evolution, plastid reduction, and obligate asexuality origins"

#### Supplementary figures and tables

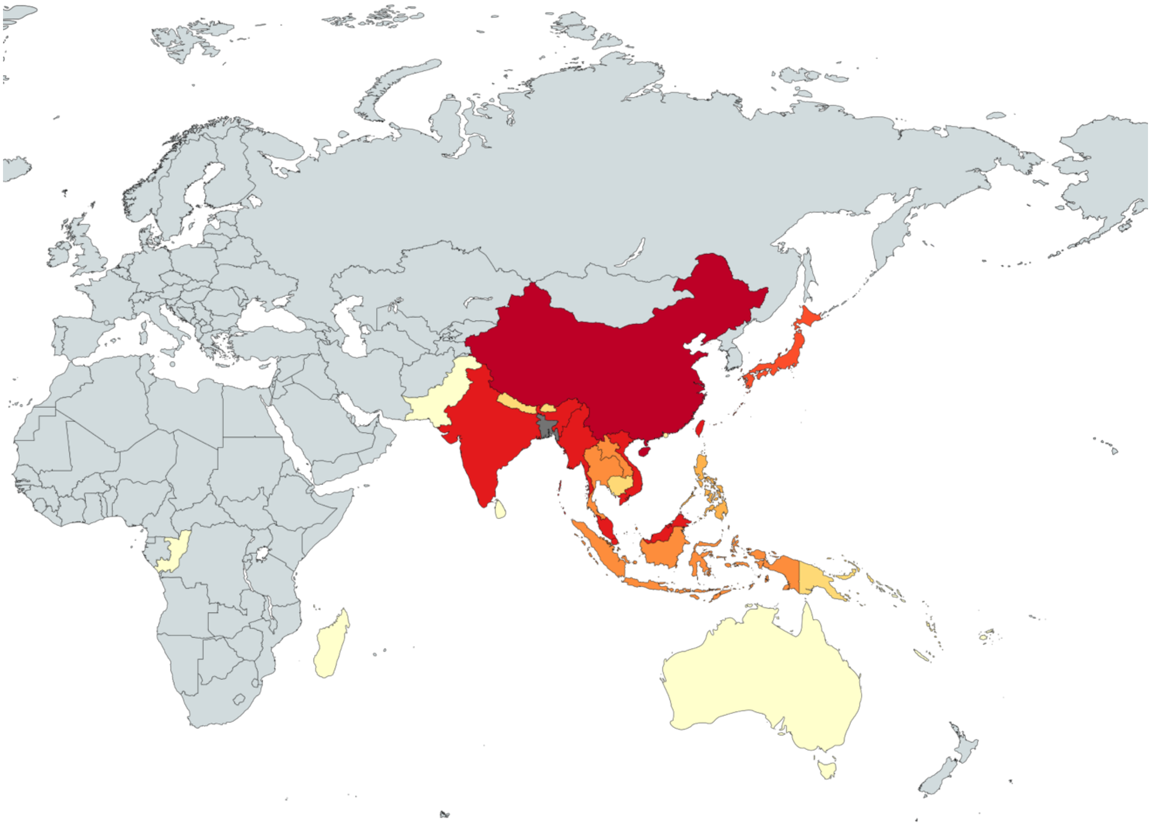

**Figure S1. *Balanophora* species diversity in the world.** The number of species is indicated by a heatmap. Light gray corresponds to areas with no *Balanophora* occurrence.

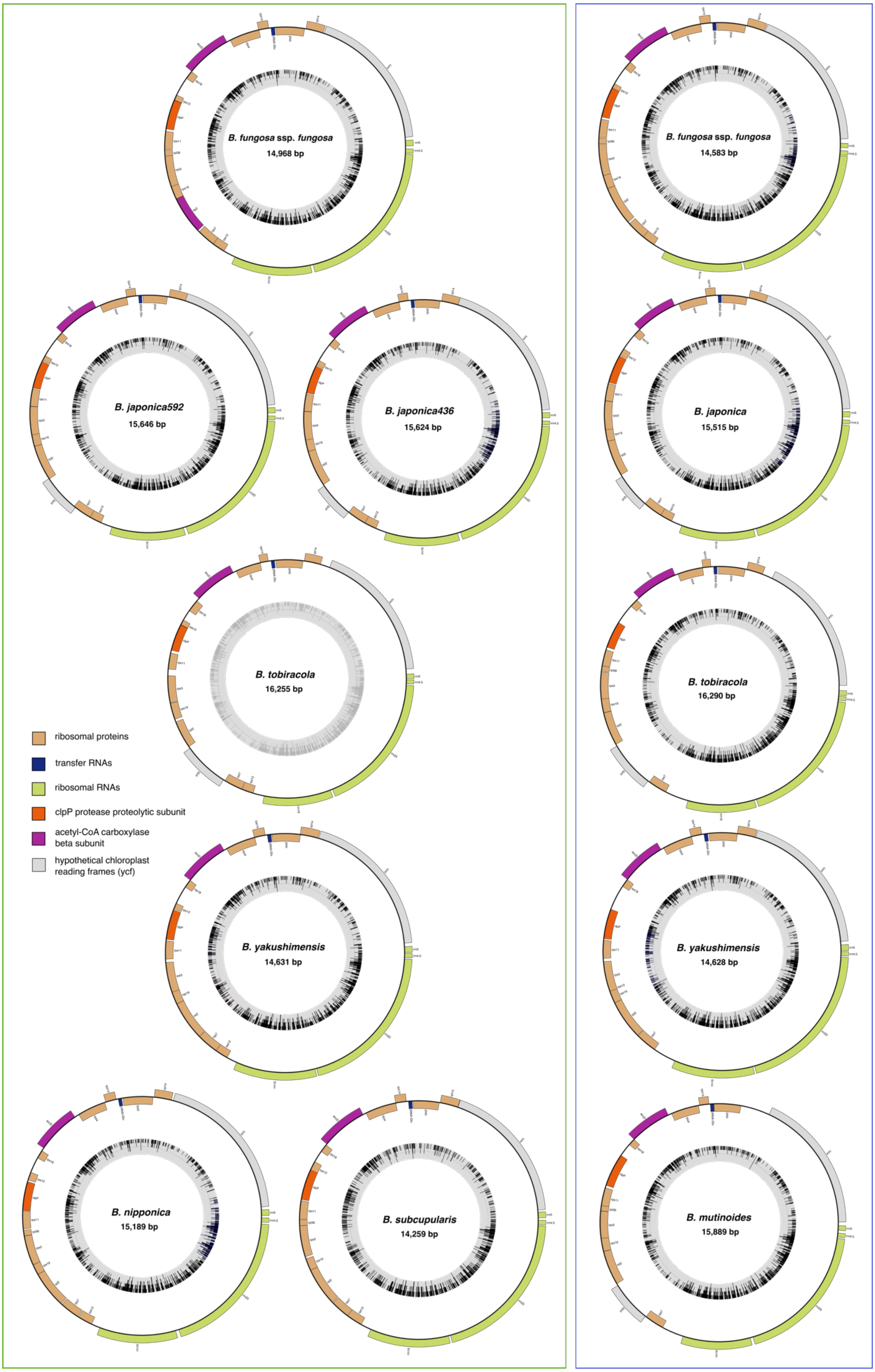


**Figure S2. Circularized plastid genomes of the studied Japanese (left) and Taiwanese (right) *Balanophora* species assembled in this study.**

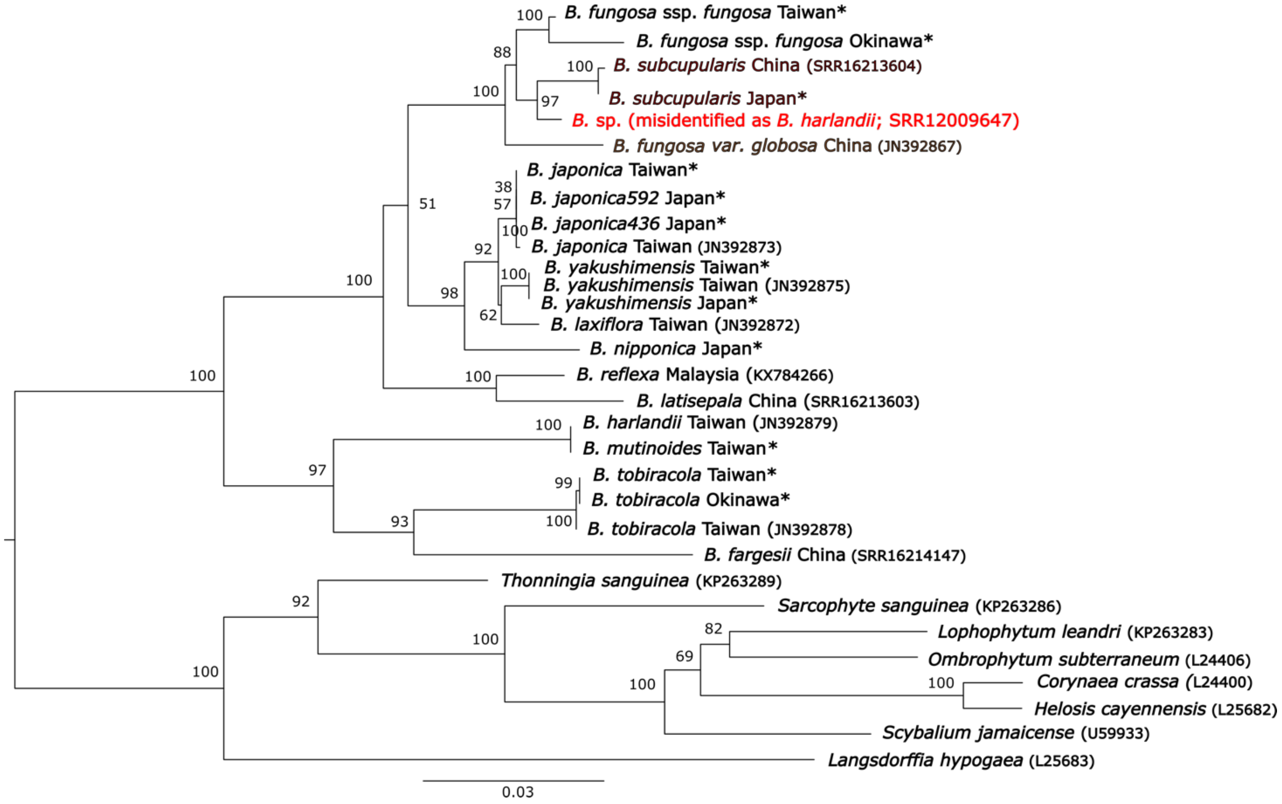


**Figure S3. Maximum likelihood phylogeny of Balanophoraceae inferred by IQTREE from the nuclear 18S rRNA gene (GTR+F+I+G4 model; 1,000 bootstraps).** The position of the misidentified *Balanophora* from NCBI is in red. The phylogenetic tree is rooted, and bootstrap values are displayed next to branches. Sequences newly generated in this study are highlighted by asterisks. GenBank accession numbers for previously published public data are indicated in parentheses. The alignment was trimmed by Gblocks before constructing the tree.

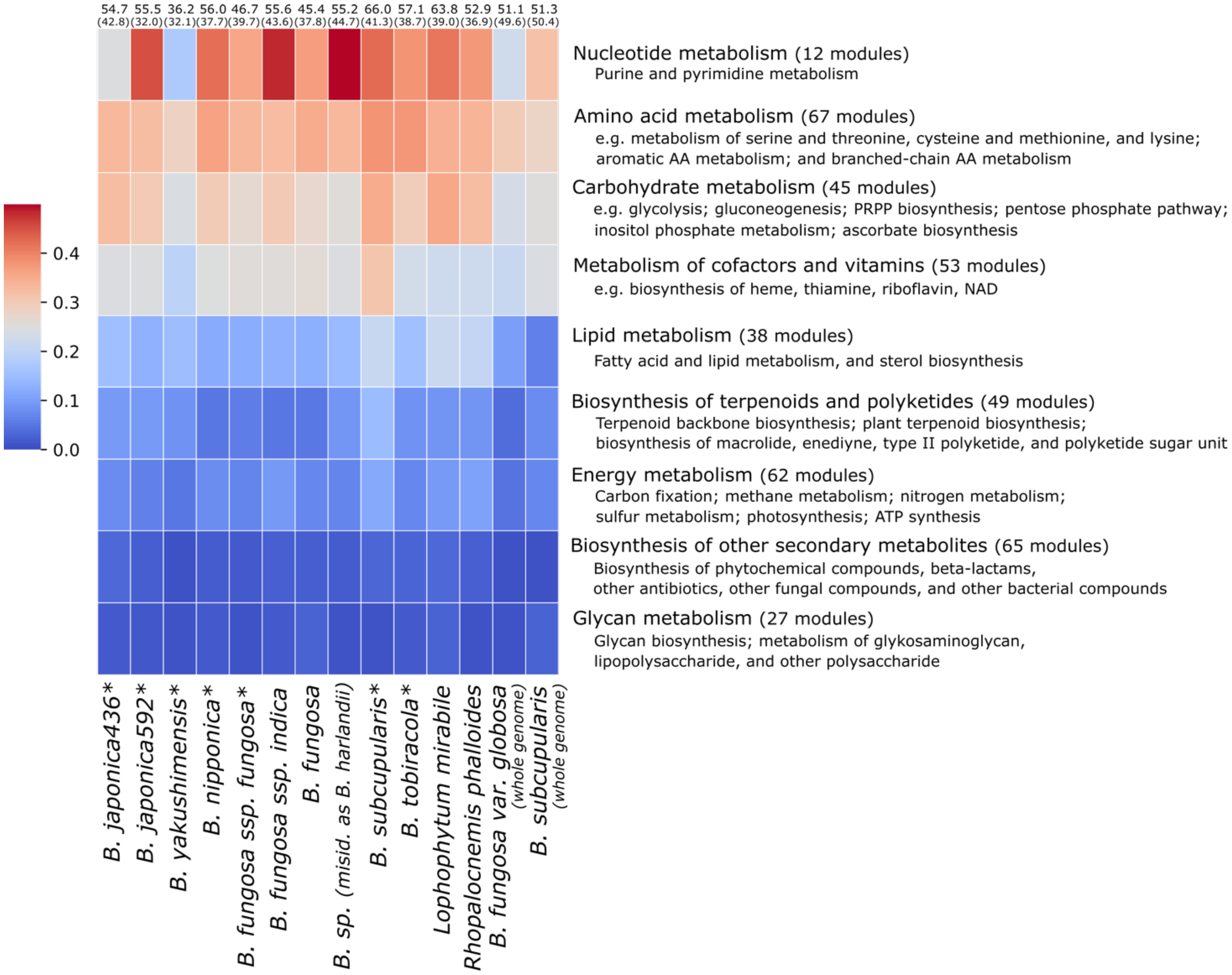


**Figure S4. Average completeness of all plastid biosynthetic pathways from nine KEGG pathway-module categories.** The plot was inferred from predicted proteomes of both newly generated data (highlighted by asterisks) and previously published data (*B. fungosa spp. indica* (SRR12009646), *B. fungosa* (ERR2040275), *B.* sp. (misidentified as *B. harlandii*; SRR12009647), *L. mirabile* (SRR10883507), *R. phalloides* (SRR14800310), *B. fungosa var. globosa* (GWHDONJ00000000), and *B. subcupularis* (GWHDONK00000000)). *B. fungosa var. globosa* and *B. subcupularis* show completeness of plastid biosynthetic pathways inferred from proteomes predicted from whole genome data. For all KEGG modules and their completeness see Figure S5. BUSCO completeness scores of the transcriptomes are indicated as percentages above the figure. Complete transcripts including single-copy and duplicated transcripts and only single-copy transcripts (in parentheses) are displayed. The samples are ordered according to their phylogeny.

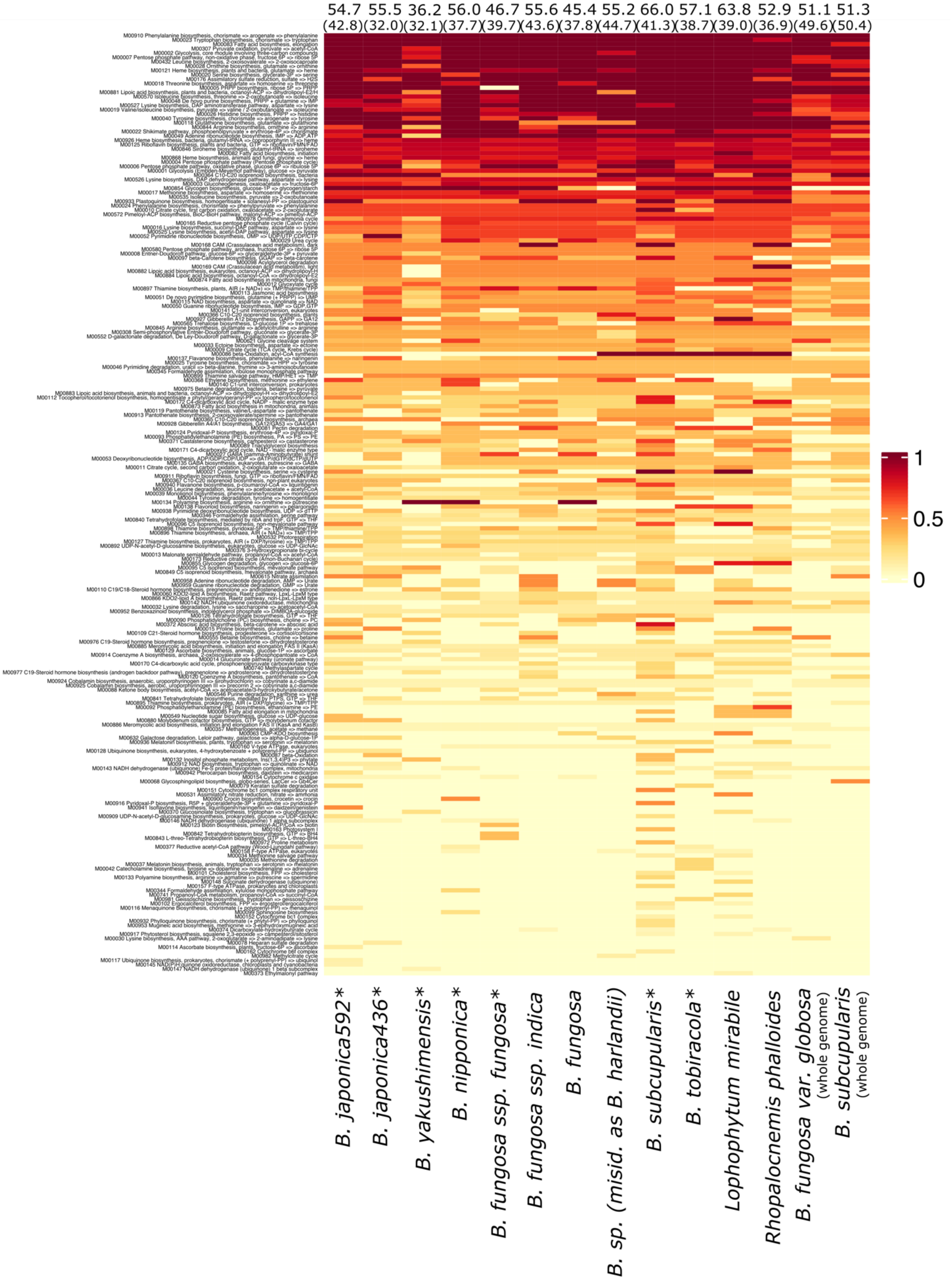


**Figure S5. Completeness of all plastid biosynthetic pathways inferred from predicted proteomes of both newly generated data (highlighted by asterisks) and previously published data** (*B. fungosa spp. indica* (SRR12009646), *B. fungosa* (ERR2040275), *B.* sp. (misidentified as *B. harlandii*; SRR12009647), *L. mirabile* (SRR10883507), *R. phalloides* (SRR14800310), *B. fungosa var. globosa* (GWHDONJ00000000), and *B. subcupularis* (GWHDONK00000000)). *B. fungosa var. globosa* and *B. subcupularis* show completeness of plastid biosynthetic pathways inferred from proteomes predicted from whole genome data. The samples are ordered according to their phylogeny.

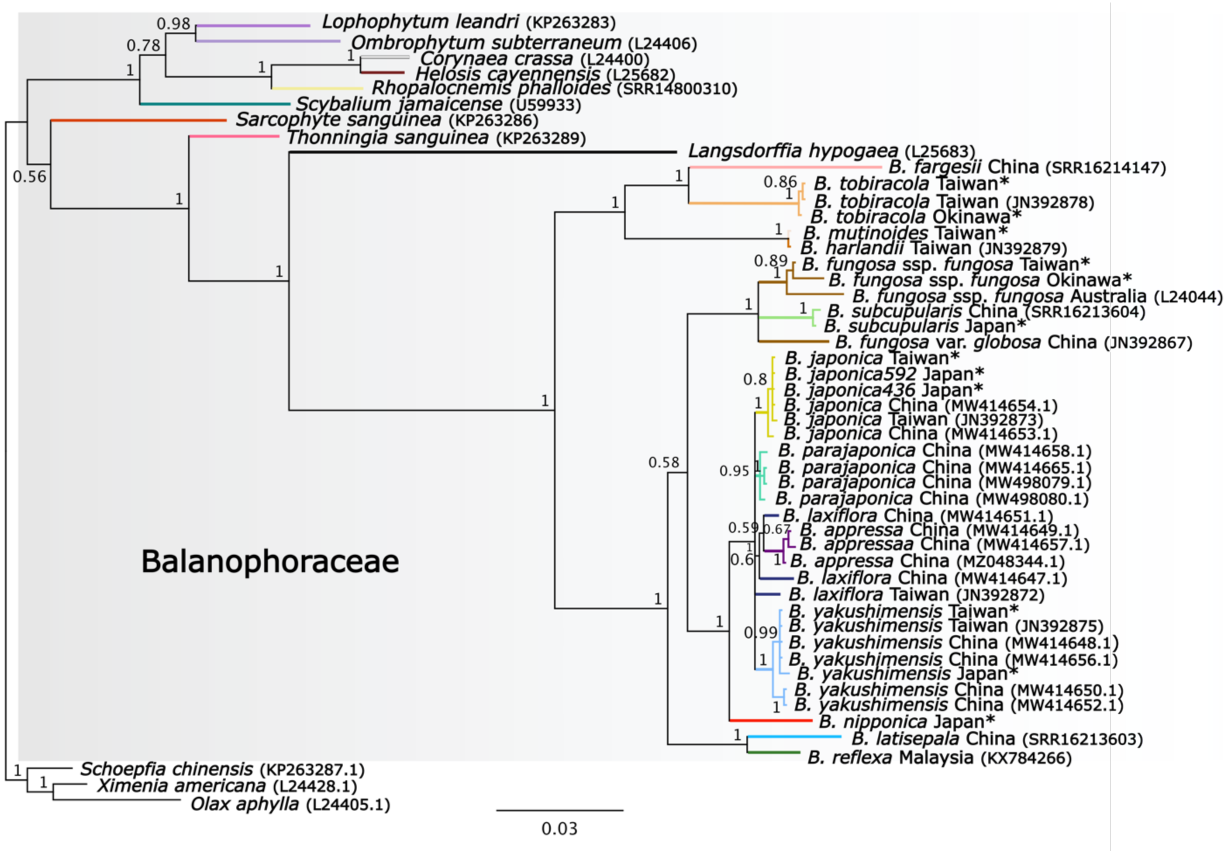


**Figure S6. Bayesian phylogeny of Balanophoraceae inferred from the nuclear 18S rRNA gene (GTR+G model; 10,000,000 generations).** Posterior probabilities are displayed next to branches. The dataset was trimmed by Gblocks before constructing the trees. The phylogenetic tree is rooted by three species from Santalales. GenBank accession numbers for sequences obtained from NCBI are indicated in parentheses.

**Table S1. The sampling sites of *Balanophora* species in mainland Japan, Okinawa, and Taiwan.**

| Sample | Locality |
|---|---|
| *B. fungosa ssp. fungosa JAP* | Okinawa, Japan |
| *B. fungosa* ssp. *fungosa TWN* | Pintung, Taiwan |
| *B. japonica592JAP* | Kyushu, Japan |
| *B. japonica436 JAP* | Kyushu, Japan |
| *B. japonica TWN* | Taipei, Taiwan |
| *B. nipponica JAP* | Honshu, Japan |
| *B. mutinoides TWN* | Nantou, Taiwan |
| *B. subcupularis JAP* | Kyushu, Japan |
| *B. tobiracola JAP* | Okinawa, Japan |
| *B. tobiracola TWN* | Hualian, Taiwan |
| *B. yakushimensis JAP* | Kyushu, Japan |
| *B. yakushimensis TWN* | Taoyuan, Taiwan |

**Table S2. The plastid genome features of the Balanophoraceae species, including genome size, GC content, and the number of protein-coding genes, rRNA genes, and tRNA genes.** Plastid genomes assembled in this study are accompanied by JAP for Japanese populations and TWN for Taiwanese populations.

| species | plastid genome features | | | | |
|---|---|---|---|---|---|
| | size (bp) | GC content (%) | protein-coding genes | rRNA genes | tRNA genes |
| *Balanophora fargesii* | 15,813 | 12.5 | 16 | 4 | 1 |
| *Balanophora fungosa* ssp. *fungosa JAP* | 14,968 | 12.5 | 15 | 4 | 1 |
| *Balanophora fungosa* ssp. *fungosa TWN* | 14,583 | 12.7 | | | |
| *Balanophora fungosa* var. *globosa* | 15,130 | 12.6 | | | |
| *Balanophora harlandii* | 16,056 | 12.9 | 15 | 4 | 1 |
| *Balanophora japonica* 436 JAP | 15,624 | 12.1 | 15 | 4 | 1 |
| *Balanophora japonica* 592 JAP | 15,646 | 12.0 | | | |
| *Balanophora japonica TWN* | 15,515 | 12.1 | | | |
| *Balanophora latisepala* | 14,951 | 11.6 | 15 | 4 | 1 |
| *Balanophora laxiflora* | 15,505 | 12.2 | 15 | 4 | 1 |
| *Balanophora mutinoides TWN* | 15,889 | 12.9 | 15 | 4 | 1 |
| *Balanophora nipponica JAP* | 15,189 | 13.0 | 15 | 4 | 1 |
| *Balanophora reflexa* | 15,507 | 11.6 | 15 | 4 | 1 |
| *Balanophora subcupularis JAP* | 14,259 | 12.9 | 15 | 4 | 1 |
| *Balanophora subcupularis* | 14,309 | 12.8 | 15 | 4 | 1 |
| *Balanophora tobiracola JAP* | 16,255 | 12.3 | 16 | 4 | 1 |
| *Balanophora tobiracola TWN* | 16,290 | 12.3 | | | |
| *Balanophora yakushimensis JAP* | 14,631 | 12.6 | 14 | 4 | 1 |
| *Balanophora yakushimensis TWN* | 14,628 | 12.6 | | | |
| *Balanophora yakushimensis* | 14,624 | 12.7 | | | |
| *Lophophytum leandri* | 20,892 | 11.6 | 12 | 3 | 0 |
| *Ombrophytum subterraneum* | 17,313 | 14.0 | 13 | 3 | 0 |
| *Rhopalocnemis phalloides* | 18,622 | 13.2 | 13 | 3 | 1 |
| *Sarcophyte sanguinea* | 28,372-28,384 | 19.1 | 19 | 4 | 4 |
| *Thonningia sanguinea* | 18,560-19,013 | 20.4-21.2 | 17 | 4 | 1 |

**Table S3. Presence/absence of plastid genes in 13 species from the Balanophoraceae family.** Pseudogenes are in light yellow. + gene fusion; - abnormal anticodon loop. Plastid genomes assembled in this study are accompanied by JAP for Japanese populations and TWN for Taiwanese populations.

| | | | | | Protein synthesis | | | | | | | | | | | | | | | | | | | | | | | | | | | | Essential | | | | |
|---|---|---|---|---|---|---|---|---|---|---|---|---|---|---|---|---|---|---|---|---|---|---|---|---|---|---|---|---|---|---|---|---|---|---|---|---|---|
| species | taxonomic group | type | size (bp) | GC content (%) | infA | matK | rpl2 | rpl14 | rpl16 | rpl20 | rpl22 | rpl23 | rpl32 | rpl33 | rpl36 | rps2 | rps3 | rps4 | rps7 | rps8 | rps11 | rps12_1 | rps12_2 | rps14 | rps15 | rps16 | rps18 | rps19 | rrn4.5 | rrn5 | rrn16 | rrn23 | ycf1 | ycf2 | accD | clpP | trnE |
| *Cuscuta gronovii* | Convolvulaceae | HP | 86,744 | 37.7 | | | | | | | | | | | | | | | | | | | | | | | | | | | | | | | | | |
| *Cuscuta obtusiflora* | Convolvulaceae | HP | 85,286 | 37.8 | | | | | | | | | | | | | | | | | | | | | | | | | | | | | | | | | |
| *Phelipanche purpurea* | Orobanchaceae | HP | 62,891 | 31.1 | | | | | | | | | | | | | | | | | | | | | | | | | | | | | | | | | |
| *Conopholis americana* | Orobanchaceae | HP | 45,673 | 33.9 | | | | | | | | | | | | | | | | | | | | | | | | | | | | | | | | | |
| *Balanophora fargesii* | Santalales | HP | 15,813 | 12.5 | | | | | | | | | | | | | | | | | | | | | | | | | | | | | | | | | |
| *Balanophora fungosa* ssp. *fungosa JAP* | Santalales | HP | 14,968 | 12.5 | | | | | | | | | | | | | | | | | | | | | | | | | | | | | | | | | |
| *Balanophora fungosa* ssp. *fungosa TWN* | Santalales | HP | 14,583 | 12.7 | | | | | | | | | | | | | | | | | | | | | | | | | | | | | | | | | |
| *Balanophora fungosa* var. *globosa* | Santalales | HP | 15,130 | 12.6 | | | | | | | | | | | | | | | | | | | | | | | | | | | | | | | | | |
| *Balanophora harlandii* | Santalales | HP | 16,056 | 12.9 | | | + | | | | | | | | | | | | | | | | | | | | | + | | | | | | | | | |
| *Balanophora japonica* 436 JAP | Santalales | HP | 15,624 | 12.1 | | | | | | | | | | | | | | | | | | | | | | | | | | | | | | | | | |
| *Balanophora japonica* 592 JAP | Santalales | HP | 15,646 | 12.0 | | | | | | | | | | | | | | | | | | | | | | | | | | | | | | | | | |
| *Balanophora japonica TWN* | Santalales | HP | 15,515 | 12.1 | | | | | | | | | | | | | | | | | | | | | | | | | | | | | | | | | |
| *Balanophora latisepala* | Santalales | HP | 14,951 | 11.6 | | | | | | | | | | | | | | | | | | | | | | | | | | | | | | | | | |
| *Balanophora laxiflora* | Santalales | HP | 15,505 | 12.2 | | | | | | | | | | | | | | | | | | | | | | | | | | | | | | | | | - |
| *Balanophora mutinoides TWN* | Santalales | HP | 15,889 | 12.9 | | | | | | | | | | | | | | | | | | | | | | | | | | | | | | | | | |
| *Balanophora nipponica JAP* | Santalales | HP | 15,189 | 13.0 | | | | | | | | | | | | | | | | | | | | | | | | | | | | | | | | | |
| *Balanophora reflexa* | Santalales | HP | 15,507 | 11.6 | | | | | | | | | | | | | | | | | | | | | | | | | | | | | | | | | - |
| *Balanophora subcupularis JAP* | Santalales | HP | 14,259 | 12.9 | | | | | | | | | | | | | | | | | | | | | | | | | | | | | | | | | |
| *Balanophora subcupularis* | Santalales | HP | 14,309 | 12.8 | | | | | | | | | | | | | | | | | | | | | | | | | | | | | | | | | |
| *Balanophora tobiracola JAP* | Santalales | HP | 16,255 | 12.3 | | | | | | | | | | | | | | | | | | | | | | | | | | | | | | | | | |
| *Balanophora tobiracola TWN* | Santalales | HP | 16,290 | 12.3 | | | | | | | | | | | | | | | | | | | | | | | | | | | | | | | | | |
| *Balanophora yakushimensis JAP* | Santalales | HP | 14,631 | 12.6 | | | | | | | | | | | | | | | | | | | | | | | | | | | | | | | | | |
| *Balanophora yakushimensis TWN* | Santalales | HP | 14,628 | 12.6 | | | | | | | | | | | | | | | | | | | | | | | | | | | | | | | | | - |
| *Balanophora yakushimensis* | Santalales | HP | 14,624 | 12.7 | | | | | | | | | | | | | | | | | | | | | | | | | | | | | | | | | |
| *Lophophytum leandri* | Santalales | HP | 20,892 | 11.6 | | | | | | | | | | | | | | | | | | 1 gene | | | | | | | | | | | | | | | |
| *Ombrophytum subterraneum* | Santalales | HP | 17,313 | 14.0 | | | | | | | | | | | | | | | | | | 1 gene | | | | | | | | | | | | | | | |

| | | | | | |
|---|---|---|---|---|---|
| *Rhopalocnemis phalloides* | Santalales | HP | 18,622 | 13.2 | 1 gene; – |
| *Sarcophyte sanguinea* | Santalales | HP | 28,372 -28,384 | 19.1 | |
| *Thonningia sanguinea* | Santalales | HP | 18,560 -19,013 | 20.4-21.2 | – |
| *Osyris alba* | Santalales | HeP | 147,253 | 37.7 | |
| *Schoepfia jasminodora* | Santalales | HeP | 118,743 | 38.1 | |
| *Viscum album* | Santalales | HeP | 128,921 | 36.4 | |
| *Pilostyles aethiopica* | Apodanthaceae | HP | 11,348 | 24.2 | |
| *Pilostyles hamiltonii* | Apodanthaceae | HP | 15,167 | 22.7 | |
| *Cytinus hypocistis* | Cytinaceae | HP | 19,400 | 29.9 | |
| *Cynomorium coccineum* | Cynomiriaceae | HP | 45,519 | 30.0 | |
| *Epipogium aphyllum* | Orchideaceae | MH | 30,650 | 32.8 | |
| *Epipogium roseum* | Orchideaceae | MH | 19,047 | 30.6 | |
| *Thismia tentaculata* | Thismiaceae | MH | 16,031 | 27.1 | |
| *Sciaplhila densiflora* | Triuridaceae | MH | 21,458 | 39.9 | |
| *Hydnora visseri* | Hydnoraceae | HP | 27,233 | 23.4 | |

**Table S4. The attributes of all protein-coding genes in Balanophoraceae, including individual gene length, GC content, and the positions of in-frame TAG codons within the genes.** Plastid genomes studied here and previously published plastid genomes are accompanied by JAP/TWN and accession numbers, respectively.

| protein-coding gene | gene attributes | *B. fargesii* (SRR16214147) | *B. fungosa* ssp. *fungosa* JAP | *B. fungosa* ssp. *fungosa* TWN | *B. fungosa* var. *globosa* (MN414176) | *B. harlandii* (MN414177) | *B. japonica* 436 JAP | *B. japonica* 592 JAP | *B. japonica* TWN | *B. latisepala* (SRR16213603) | *B. laxiflora* (KX784265) | *B. mutinoides* TWN | *B. nipponica* JAP |
|---|---|---|---|---|---|---|---|---|---|---|---|---|---|
| *accD* | Length (bp)<br>GC content (%)<br>In-frame TAG<br>(TGA)[TGG]codons | 915<br>17.37<br>19;204 | 915<br>17.15<br>19;204 | 912<br>17.32<br>19;204 | 900<br>17.33<br>19;204 | 908<br>17.29<br>19;204 | 894<br>17.22<br>19;204 | 894<br>17.22<br>19;204 | 888<br>17.22<br>19;204 | 888<br>17.45<br>20;204 | 894<br>17.56<br>19;204 | 900<br>17.33<br>19;<br>204 | 900<br>18.0<br>19;204 |
| *clpP* | Length<br>GC content (%)<br>In-frame TAG<br>codons | 597<br>18.42<br>22;172 | 597<br>19.59<br>22;78;<br>172 | 615<br>19.18<br>22;78;<br>172 | 609<br>19.04<br>22;78;<br>[172] | 642<br>19.47<br>22;78;<br>172 | 597<br>19.93<br>22;78;<br>172 | 597<br>19.93<br>22;78;<br>172 | 597<br>19.93<br>22;78;<br>172 | 588<br>18.53<br>22 | 600<br>19.66<br>22;78;<br>172 | 717<br>19.8<br>22;78;<br>172 | 597<br>19.93<br>22;78;<br>172 |
| *matK* | Length<br>GC content (%)<br>In-frame TAG<br>codons | | | | | | | | | | | | |
| *rpl2* | Length (bp)<br>GC content (%)<br>In-frame TAG<br>codons | 738<br>14.76<br>247;28<br>4 | 759<br>14.22<br>35;247<br>;284 | 762<br>14.17<br>35;247<br>;284 | 768<br>14.71<br>35;247<br>;284 | 1026+<br>13.35<br>247;28<br>4 | 765<br>13.98<br>35;247<br>;284 | 756<br>14.15<br>35;247<br>;284 | 756<br>14.15<br>35;247<br>;284 | 765<br>13.2<br>247;28<br>4 | 750<br>14.26<br>35;247<br>;284 | 741<br>15.24<br>247;28<br>4 | 762<br>14.69<br>35;247<br>;284 |
| *rpl14* | Length (bp)<br>GC content (%)<br>In-frame TAG<br>codons | 354<br>11.01<br>x | 348<br>10.63<br>x | 348<br>10.63<br>x | 351<br>10.54<br>x | | 357<br>10.64<br>111 | 357<br>10.64<br>111 | 357<br>10.64<br>111 | 342<br>11.11<br>x | 357<br>10.36<br>111 | | 354<br>11.01<br>x |
| *rpl16* | Length (bp)<br>GC content (%) | | | | | | | | | | | | |
| *rpl36* | Length (bp)<br>GC content (%) | 114<br>11.4<br>x | 117<br>11.11<br>x | 117<br>11.11<br>x | 168<br>10.11<br>x | 117<br>11.96<br>x | | | | | | 117<br>11.11<br>x | 117<br>10.25<br>x |
| *rps2* | Length (bp)<br>GC content (%)<br>In-frame TAG<br>codons | 537<br>5.4<br>33;110 | 573<br>3.66<br>x | 558<br>3.94<br>x | 531<br>4.89<br>110 | 543<br>6.99<br>16;110 | 522<br>4.4<br>18;210 | 522<br>4.4<br>18;210 | 522<br>4.4<br>18;210 | 543<br>4.05<br>18;110 | 558<br>4.83<br>18;210 | 579<br>6.56<br>16;110 | 618<br>5.33<br>18 |
| *rps3* | Length (bp)<br>GC content (%) | 615<br>5.52<br>x | 636<br>4.4<br>x | 636<br>4.55<br>x | 618<br>5.82<br>x | 597<br>5.52<br>x | 612<br>5.88<br>x | 612<br>5.88<br>x | 612<br>5.88<br>x | 615<br>5.04<br>x | 624<br>5.6<br>x | 594<br>5.55<br>x | 612<br>6.53<br>x |
| *rps4* | Length (bp)<br>GC content (%)<br>In-frame TAG<br>codons | 582<br>7.9<br>x | 606<br>8.74<br>x | 600<br>8.0<br>x | 603<br>8.95<br>x | 606<br>7.42<br>x | 591<br>9.47<br>72 | 591<br>9.47<br>72 | 591<br>9.47<br>72 | 615<br>7.8<br>x | 588<br>9.01<br>69 | 600<br>7.66<br>x | 582<br>8.59<br>x |
| *rps7* | Length (bp) | 402 | 378 | 378 | 414 | 420 | 414 | 414 | 414 | 432 | 429 | 423 | 411 |

| protein-coding gene | gene attributes | *B. reflexa* (KX784266) | *B. subcupularis* JAP | *B. subcupularis* (SRR16213604) | *B. tobiracola* JAP | *B. tobiracola* TWN | *B. yakushimensis* JAP | *B. yakushimensis* TWN | *B. yakushimensis* (NC_062820) | *Lophophytum leandri* (MT834848) | *Ombrophytum subterraneum* (MT834847) | *Rhopalocnemis phalloides* (MK036331] | *Sarcophyte sanguinea* (NC_080979) | *Thonningia sanguinea* [OQ810031] |
|---|---|---|---|---|---|---|---|---|---|---|---|---|---|---|
| *accD* | Length (bp)<br>GC content (%)<br>In-frame TAG<br>(TGA)[TGG]codons | 951<br>16.19<br>19;204 | 897<br>17.61<br>19;204 | 894<br>17.67<br>19;204 | 939<br>16.82<br>19;204 | 939<br>16.82<br>19;204 | 888<br>17.45<br>19;204 | 888<br>17.45<br>19;204 | 888<br>17.45<br>19;204 | 927<br>17.25<br>[204; 250] | 936<br>17.94<br>(204;250)<br>[21] | 915<br>20.43<br>[204; 250] | 1290<br>24.65<br>[204; 250] | 1062<br>23.72<br>19; 250<br>[204] |
| *clpP* | Length<br>GC content (%)<br>In-frame TAG<br>codons | 606<br>19.47<br>22 | 597<br>19.26<br>22;78;<br>[172] | 594<br>19.52<br>22;78;<br>[172] | 594<br>19.19<br>22;78;<br>172 | 594<br>19.19<br>22;78;<br>172 | 597<br>19.43<br>22;172 | 597<br>19.43<br>22;172 | 597<br>19.43<br>22;172 | 225;301;74<br>18.66;21.26;<br>22.97<br>(3pieces)<br>[22] | 225;292;71<br>22.22;22.26;<br>26.76<br>(3pieces)<br>(22) | 225;292;74<br>25.33;<br>24.31;32.43<br>(3pieces)<br>[22; 172] | 77;292;234<br>41.55;32.19;<br>30.34<br>(3pieces)<br>[22; 78] | 71;517<br>22.53; 28.23<br>22;55;172 |
| *matK* | Length<br>GC content (%)<br>In-frame TAG<br>codons | | | | | | | | | | | | 1446<br>9.75<br>[81;284;291<br>375;472] | |
| *rpl2* | Length (bp)<br>GC content (%)<br>In-frame TAG<br>codons | 768<br>13.93<br>247;28<br>4 | 744<br>14.38<br>35;247<br>;284 | 744<br>14.24<br>35;247<br>;284 | 738<br>15.04<br>247;28<br>4 | 738<br>15.04<br>247;28<br>4 | 777<br>13.77<br>35;247<br>;[284] | 786<br>13.61<br>35;247<br>;[284] | 777<br>13.77<br>35;247<br>;[284] | 261;417<br>12.26;20.38<br>(2pieces)<br>(284)[202] | 262;407<br>12.97;20.14<br>(2pieces)<br>17.33<br>(202;284) | 216;312<br>10.18;12.82<br>(2pieces)<br>[202] | 413;388<br>26.87;19.58<br>(2pieces)<br>[11;284] | 678<br>28.02<br>247;[284] |
| *rpl14* | Length (bp)<br>GC content (%)<br>In-frame TAG<br>codons | 345<br>10.43<br>x | 348<br>10.05<br>x | 345<br>10.14<br>x | 360<br>11.38<br>x | 360<br>11.38<br>x | 354<br>10.73<br>111 | 354<br>10.73<br>111 | 354<br>10.73<br>111 | | | | 369<br>24.11<br>x | 372<br>22.58<br>x |
| *rpl16* | Length (bp)<br>GC content (%) | | | | | | | | | | | 9;417<br>33.33;16.78<br>(2pieces)<br>[64;92] | 9;408<br>33.33;24.75<br>(2pieces)<br>[66;94] | 405<br>22.96<br>10;92 |
| *rpl36* | Length (bp)<br>GC content (%) | | 117<br>11.11<br>x | 114<br>11.4<br>x | 117<br>11.96<br>x | 117<br>11.96<br>x | | | | | 114<br>15.78<br>x | 126<br>11.11<br>x | 114<br>19.29<br>x | 111<br>18.01<br>x |
| *rps2* | Length (bp)<br>GC content (%)<br>In-frame TAG<br>codons | 549<br>4.0<br>18;110 | 504<br>3.57<br>x | 498<br>3.61<br>x | 594<br>5.72<br>110 | 594<br>5.72<br>110 | 562<br>4.62<br>18 | 561<br>4.81<br>18 | 562<br>4.62<br>18 | | | | 696<br>20.54<br>[17;219] | 654<br>17.73<br>14;16;40;110 |
| *rps3* | Length (bp)<br>GC content (%) | 624<br>4.96<br>x | 606<br>5.28<br>x | 603<br>5.14<br>x | 597<br>5.52<br>x | 597<br>5.52<br>x | 606<br>5.28<br>x | 606<br>5.28<br>x | 606<br>5.28<br>x | 636<br>8.8<br>[25;220] | 636<br>9.74<br>(25;186;220) | 639<br>10.17<br>[25;220] | 636<br>15.4<br>[25;186;220] | 618<br>14.88<br>186 |
| *rps4* | Length (bp)<br>GC content (%)<br>In-frame TAG<br>codons | 525<br>6.09<br>x | 603<br>7.79<br>x | 597<br>7.87<br>x | 555<br>8.1<br>x | 558<br>8.06<br>x | 594<br>8.24<br>x | 594<br>8.24<br>x | 594<br>8.24<br>x | 546<br>6.22<br>x | 540<br>5.92<br>(107) | | 603<br>13.75<br>x | 333<br>18.91<br>233 |
| *rps7* | Length (bp) | 408 | 378 | 375 | 420 | 420 | 426 | 426 | 426 | 414 | 360 | 438 | 444 | 468 |

| protein-coding gene | gene attributes | *B. fargesii* (SRR16214147) | *B. fungosa* ssp. *fungosa* JAP | *B. fungosa* ssp. *fungosa* TWN | *B. fungosa* var. *globosa* (MN414176) | *B. harlandii* (MN414177) | *B. japonica* 436 JAP | *B. japonica* 592 JAP | *B. japonica* TWN | *B. latisepala* (SRR16213603) | *B. laxiflora* (KX784265) | *B. mutinoides* TWN | *B. nipponica* JAP | *B. reflexa* (KX784266) | *B. subcupularis* JAP | *B. subcupularis* (SRR16213604) | *B. tobiracola* JAP | *B. tobiracola* TWN | *B. yakushimensis* JAP | *B. yakushimensis* TWN | *B. yakushimensis* (NC_062820) | *Lophophytum leandri* (MT834848) | *Ombrophytum subterraneum* (MT834847) | *Rhopalocnemis phalloides* (MK036331) | *Sarcophyte sanguinea* (NC_080979) | *Thonningia sanguinea* (OQ810031) |
|---|---|---|---|---|---|---|---|---|---|---|---|---|---|---|---|---|---|---|---|---|---|---|---|---|---|---|
| | GC content (%) | 5.22<br>x | 4.23<br>x | 4.23<br>x | 5.31<br>x | 5.71<br>x | 5.07<br>x | 5.07<br>x | 5.07<br>x | 4.39<br>x | 5.36<br>x | 5.43<br>x | 5.59<br>x | 4.9<br>x | 4.76<br>x | 4.8<br>x | 5.23<br>x | 5.23<br>x | 5.86<br>x | 5.86<br>x | 5.86<br>x | 11.35<br>x | 11.94<br>x | 10.73<br>[39] | 15.54<br>x | 19.44<br>148 |
| *rps8* | Length (bp)<br>GC content (%) | | | | | | | | | | | | | | | | | | | | | | | | 402<br>17.16<br>x | |
| *rps11* | Length (bp)<br>GC content (%) | 369<br>11.38<br>x | 366<br>12.02<br>x | 354<br>12.71<br>x | 366<br>12.84<br>x | 417<br>10.55<br>x | 369<br>11.38<br>x | 369<br>11.38<br>x | 372<br>11.29<br>x | 363<br>11.01<br>x | 372<br>11.82<br>x | 420<br>10.0<br>x | 366<br>11.74<br>x | 372<br>11.55<br>x | 357<br>11.48<br>x | 354<br>11.58<br>x | 357<br>11.48<br>x | 357<br>11.48<br>x | 363<br>11.84<br>x | 363<br>11.84<br>x | 363<br>11.84<br>x | | | | 501<br>18.76<br>[76] | 384<br>25.78<br>76 |
| *rps12a* | Length (bp)<br>GC content (%) | 114<br>14.91<br>x | 114<br>9.64<br>x | 99<br>12.12<br>x | 114<br>8.77<br>x | 123<br>15.44<br>x | 117<br>10.25<br>x | 117<br>10.25<br>x | 114<br>10.52<br>x | 99<br>11.11<br>x | 111<br>10.81<br>x | 111<br>18.01<br>x | 168<br>9.52<br>x | 114<br>9.64<br>x | 180<br>7.22<br>x | 111<br>9.0<br>x | 120<br>12.5<br>x | 111<br>13.51<br>x | 120<br>10.83<br>x | 114<br>11.4<br>x | 120<br>10.83<br>x | 105;23;232<br>15.23;13.04;<br>27.58<br>(3pieces)<br>x | 111;26;232<br>17.11;11.53;<br>27.15<br>(3pieces)<br>x | 96;29;235<br>15.62;10.34;<br>27.23<br>(3pieces)<br>x | 120;232<br>31.66;33.18<br>26<br>11.53<br>(3pieces)<br>x | 261;129<br>32.95;19.37<br>(2pieces)<br>x |
| *rps12b* | Length (bp)<br>GC content (%) | 258<br>24.41<br>x | 258<br>24.03<br>x | 258<br>24.03<br>x | 258<br>25.96<br>x | 258<br>25.58<br>x | 258<br>24.41<br>x | 258<br>24.41<br>x | 258<br>24.41<br>x | 261<br>23.75<br>x | 258<br>24.03<br>x | 258<br>26.35x | 258<br>24.41<br>x | 261<br>23.75<br>x | 261<br>23.37<br>x | 261<br>23.37<br>x | 261<br>24.52<br>x | 261<br>24.52<br>x | 258<br>24.03<br>x | 258<br>24.03<br>x | 258<br>24.03<br>x | | | | | |
| *rps14* | Length (bp)<br>GC content (%)<br>In-frame TAG codons | 201<br>6.46<br>x | 204<br>7.84<br>x | 201<br>7.96<br>x | 210<br>7.61<br>x | 222<br>6.3<br>x | 198<br>8.08<br>96 | 198<br>8.08<br>96 | 198<br>8.08<br>96 | 213<br>6.1<br>x | 210<br>8.09<br>96 | 201<br>6.46<br>x | 219<br>6.84<br>x | 204<br>6.86<br>x | 210<br>7.61<br>x | 204<br>7.35<br>x | 180<br>6.11<br>x | 180<br>6.11<br>x | 216<br>7.87<br>x | 216<br>7.87<br>x | 216<br>7.87<br>x | 267<br>9.36<br>x | 285<br>7.01<br>x | 258<br>11.62<br>x | 303<br>21.45<br>[6;46] | 204<br>17.64<br>x |
| *rps18* | Length (bp)<br>GC content (%)<br>In-frame TAG codons | 144<br>5.55<br>x | 171<br>3.5<br>x | 162<br>3.7<br>x | 171<br>4.67<br>x | 165<br>9.69<br>x | 171<br>4.09<br>x | 171<br>4.09<br>x | 171<br>4.09<br>x | 153<br>3.26<br>x | 171<br>4.09<br>x | 153<br>10.45<br>x | 174<br>5.17<br>x | 165<br>4.24<br>x | 159<br>6.28<br>10 | 156<br>6.41<br>10 | 159<br>6.91<br>x | 189<br>6.34<br>x | 165<br>4.24<br>x | 165<br>4.24<br>x | 165<br>4.24<br>x | 243<br>8.23<br>x | 207<br>9.66<br>x | 177<br>11.29<br>x | 231<br>16.88<br>x | 201<br>14.92<br>76 |
| *rps19* | Length (bp)<br>GC content (%)<br>In-frame TAG codons | 282<br>6.73<br>x | 267<br>7.49<br>x | 267<br>7.49<br>x | 237<br>8.01<br>x | 1026+<br>13.35<br>x | 249<br>7.22<br>x | 249<br>7.22<br>x | 249<br>7.22<br>x | 240<br>6.66<br>x | 249<br>6.42<br>x | 282<br>6.38<br>x | 222<br>9.45<br>x | 219<br>7.3<br>x | 240<br>7.91<br>x | 231<br>7.35<br>x | 294<br>6.12<br>x | 306<br>6.53<br>x | 249<br>6.82<br>x | 249<br>6.82<br>x | 249<br>6.82<br>x | 270<br>10.74<br>[38] | 243<br>14.4<br>[38] | 258<br>13.56<br>[38] | 276<br>17.39<br>[38] | 291<br>20.27<br>x |
| *ycf1* | Length (bp)<br>GC content (%)<br>In-frame TAG codons | 2889<br>5.36<br>39;754;831;883;1781 | 2760<br>5.18<br>687;754;831;883;890[1781] | 2670<br>5.24<br>687;754;831;883;890;1781 | 2943<br>5.02<br>754;831;883;890;1781 | 2850<br>6.07<br>39;691;710;831;883;1781 | 2994<br>4.67<br>831;883;890 | 2961<br>4.72<br>831;883;890 | 2904<br>4.82<br>831;883;890 | 2616<br>4.24<br>392;687;754;883;890;1781 | 2943<br>4.92<br>701;831;883;890 | 2868<br>6.27<br>39;369;691;710;831;883;1781 | 2979<br>5.33<br>831;883;890;1781 | 2691<br>5.16<br>9;174;687;754;883;890;1781 | 2598<br>5.35<br>754;831;883;890;1781 | 2631<br>5.28<br>754;831;883;890;1781 | 2913<br>6.0<br>39;714;754;831;883;1781 | 965<br>6.07<br>39;714;754;831;883;1781 | 2802<br>5.06<br>831;883;890 | 2793<br>5.11<br>831;883;890 | 2802<br>5.06<br>831;883;890 | 3543<br>7.28<br>(831;890)[883;921;960;1499] | 3018<br>7.58<br>(792;831;883;921)[890;960] | 3132<br>9.03<br>[41;883;921] | 4164<br>13.35<br>[426;477;589;687;831;890;921;941;1482;1499;1781;1796] | 3174<br>16.09<br>95;220;386;550;663;687;854;883;890;921;1113;1664;1781[733] |
| *ycf2* | Length (bp)<br>GC content (%)<br>In-frame TAG codons | 1035<br>2.99<br>x | | | | 858<br>2.91<br>1230 | 753<br>2.39<br>1227 | 747<br>2.4<br>1227 | 732<br>2.45<br>x | 732<br>1.5<br>x | 750<br>2.0<br>x | 885<br>2.48<br>x | | 771<br>2.2<br>386 | | | 1008<br>2.18<br>1024 | 1026<br>2.14<br>1024 | | | | 2010<br>5.42<br>(884;1399) | 2466<br>5.43<br>(1399)<br>[884] | 2475<br>7.35<br>[1398] | 4371<br>12.17<br>[558;610;625;1075;1112;1234;1296;1326] | 2253<br>12.07<br>82;92;145;521;674;735;1218;1407;1417;1435 |

**Table S5. Nucleotide identity (%) within the *Balanophora* plastomes aligned by MAFFT (E-INS-I algorithm).** Plastid genomes studied here and already known plastid genomes are accompanied by JAP/TWN and accession numbers, respectively. Identity exceeding 85% is highlighted.

| sample | *B. fargesii (SRR16214147)* | *B. fungosa fungosa JAP* | *B.fungosa fungosa TWN* | *B. fungosa* var. *globosa (MN414176)* | *B. harlandii (MN414177)* | *B. japonica436 JAP* | *B. japonica592 JAP* | *B. japonica TWN* | *B. latisepala (SRR16213603)* | *B. laxiflora (KX784265)* | *B. mutinoides TWN* | *B. nipponica JAP* | *B. reflexa (KX784266)* | *B. subcupularis JAP* | *B. subcupularis (SRR16213604)* | *B. tobiracola JAP* | *B. tobiracola TWN* | *B. yakushimensis JAP* | *B. yakushimensis TWN* | *B. yakushimensis (NC_062820)* |
|---|---|---|---|---|---|---|---|---|---|---|---|---|---|---|---|---|---|---|---|---|
| *B. fargesii (SRR16214147)* | | 69.21 | 69.68 | 68.73 | 71.48 | 70.35 | 70.31 | 70.76 | 70.26 | 71.79 | 73.56 | 69.40 | 70.38 | 69.01 | 67.53 | 76.69 | 76.77 | 69.00 | 68.99 | 69.00 |
| *B. fungosa fungosa JAP* | 69.21 | | 95.93 | 82.84 | 65.26 | 72.49 | 72.44 | 72.89 | 69.97 | 73.37 | 66.75 | 76.88 | 70.24 | 82.73 | 80.85 | 67.30 | 67.40 | 76.82 | 76.66 | 76.82 |
| *B.fungosa fungosa TWN* | 69.68 | 95.93 | | 82.41 | 65.41 | 72.79 | 72.76 | 73.01 | 70.89 | 73.57 | 66.61 | 76.44 | 71.30 | 83.38 | 80.96 | 67.38 | 66.91 | 77.06 | 76.95 | 77.06 |
| *B. fungosa* var. *globosa (MN414176)* | 68.73 | 82.84 | 82.41 | | 64.79 | 72.31 | 72.26 | 72.37 | 69.89 | 73.07 | 65.37 | 76.73 | 69.90 | 85.80 | 83.80 | 67.09 | 66.54 | 76.90 | 76.82 | 76.90 |
| *B. harlandii (MN414177)* | 71.48 | 65.26 | 65.41 | 64.79 | | 66.43 | 66.33 | 66.75 | 65.73 | 67.54 | 89.98 | 66.16 | 65.93 | 65.12 | 63.85 | 69.78 | 69.95 | 65.22 | 65.17 | 65.22 |
| *B. japonica436 JAP* | 70.35 | 72.49 | 72.79 | 72.31 | 66.43 | | 98.92 | 98.85 | 71.57 | 85.64 | 67.68 | 76.40 | 71.05 | 71.66 | 70.82 | 68.07 | 68.26 | 81.60 | 81.72 | 81.60 |
| *B. japonica592 JAP* | 70.31 | 72.44 | 72.76 | 72.26 | 66.33 | 98.92 | | 98.77 | 71.41 | 85.59 | 67.61 | 76.49 | 70.94 | 71.65 | 70.82 | 68.00 | 68.21 | 81.69 | 81.74 | 81.69 |
| *B. japonica TWN* | 70.76 | 72.89 | 73.01 | 72.37 | 66.75 | 98.85 | 98.77 | | 72.06 | 85.71 | 67.68 | 76.36 | 71.60 | 72.08 | 70.43 | 68.41 | 68.02 | 82.10 | 82.18 | 82.10 |
| *B. latisepala (SRR16213603)* | 70.26 | 69.97 | 70.89 | 69.89 | 65.73 | 71.57 | 71.41 | 72.06 | | 72.52 | 67.33 | 69.32 | 79.32 | 70.44 | 68.97 | 67.25 | 67.50 | 70.21 | 70.22 | 70.21 |
| *B. laxiflora (KX784265)* | 71.79 | 73.37 | 73.57 | 73.07 | 67.54 | 85.64 | 85.59 | 85.71 | 72.52 | | 68.93 | 77.83 | 71.57 | 73.00 | 72.27 | 69.48 | 69.74 | 82.46 | 82.39 | 82.46 |
| *B. mutinoides TWN* | 73.56 | 66.75 | 66.61 | 65.37 | 89.98 | 67.68 | 67.61 | 67.68 | 67.33 | 68.93 | | 66.75 | 67.76 | 66.67 | 64.57 | 71.47 | 71.21 | 66.54 | 66.50 | 66.54 |
| *B. nipponica JAP* | 69.40 | 76.88 | 76.44 | 76.73 | 66.16 | 76.40 | 76.49 | 76.36 | 69.32 | 77.83 | 66.75 | | 69.37 | 76.61 | 75.23 | 67.96 | 67.19 | 81.54 | 81.57 | 81.54 |
| *B. reflexa (KX784266)* | 70.38 | 70.24 | 71.30 | 69.90 | 65.93 | 71.05 | 70.94 | 71.60 | 79.32 | 71.57 | 67.76 | 69.37 | | 70.35 | 68.77 | 68.14 | 68.34 | 69.87 | 69.93 | 69.87 |
| *B. subcupularis JAP* | 69.01 | 82.73 | 83.38 | 85.80 | 65.12 | 71.66 | 71.65 | 72.08 | 70.44 | 73.00 | 66.67 | 76.61 | 70.35 | | 96.41 | 67.06 | 67.23 | 77.19 | 77.04 | 77.19 |
| *B. subcupularis (SRR16213604)* | 67.53 | 80.85 | 80.96 | 83.80 | 63.85 | 70.82 | 70.82 | 70.43 | 68.97 | 72.27 | 64.57 | 75.23 | 68.77 | 96.41 | | 65.74 | 65.16 | 75.42 | 75.41 | 75.42 |
| *B. tobiracola JAP* | 76.69 | 67.30 | 67.38 | 67.09 | 69.78 | 68.07 | 68.00 | 68.41 | 67.25 | 69.48 | 71.47 | 67.96 | 68.14 | 67.06 | 65.74 | | 98.54 | 66.81 | 66.78 | 66.81 |
| *B. tobiracola TWN* | 76.77 | 67.40 | 66.91 | 66.54 | 69.95 | 68.26 | 68.21 | 68.02 | 67.50 | 69.74 | 71.21 | 67.19 | 68.34 | 67.23 | 65.16 | 98.54 | | 66.80 | 66.78 | 66.80 |
| *B. yakushimensis JAP* | 69.00 | 76.82 | 77.06 | 76.90 | 65.22 | 81.60 | 81.69 | 82.10 | 70.21 | 82.46 | 66.54 | 81.54 | 69.87 | 77.19 | 75.42 | 66.81 | 66.80 | | 99.23 | 100.00 |
| *B. yakushimensis TWN* | 68.99 | 76.66 | 76.95 | 76.82 | 65.17 | 81.72 | 81.74 | 82.18 | 70.22 | 82.39 | 66.50 | 81.57 | 69.93 | 77.04 | 75.41 | 66.78 | 66.78 | 99.23 | | 99.23 |
| *B. yakushimensis (NC_062820)* | 69.00 | 76.82 | 77.06 | 76.90 | 65.22 | 81.60 | 81.69 | 82.10 | 70.21 | 82.46 | 66.54 | 81.54 | 69.87 | 77.19 | 75.42 | 66.81 | 66.80 | 100.00 | 99.23 | |

**Table S6. Pairwise genetic distance between *Balanophora* plastid genomes shown as the number of base substitutions per site.** Analyses were conducted in MEGA11 (Tamura et al. 2021) using the Jukes-Cantor model (Jukes and Cantor 1969) with 4 categories of a gamma distribution. Plastid genomes are accompanied by JAP/TWN and accession numbers, respectively.

| sample | *B. fargesii (SRR16214147)* | *B. fungosa fungosa JAP* | *B.fungosa fungosa TWN* | *B. fungosa* var. *globosa (MN414176)* | *B. harlandii (MN414177)* | *B. japonica436 JAP* | *B. japonica592 JAP* | *B. japonica TWN* | *B. latisepala (SRR16213603)* | *B. laxiflora (KX784265)* | *B. mutinoides TWN* | *B. nipponica JAP* | *B. reflexa (KX784266)* | *B. subcupularis JAP* | *B. subcupularis (SRR16213604)* | *B. tobiracola JAP* | *B. tobiracola TWN* | *B. yakushimensis JAP* | *B. yakushimensis TWN* | *B. yakushimensis (NC_062820)* |
|---|---|---|---|---|---|---|---|---|---|---|---|---|---|---|---|---|---|---|---|---|
| *B. fargesii (SRR16214147)* | | | | | | | | | | | | | | | | | | | | |
| *B. fungosa fungosa JAP* | 0.1904 | | | | | | | | | | | | | | | | | | | |
| *B.fungosa fungosa TWN* | 0.1874 | 0.0096 | | | | | | | | | | | | | | | | | | |
| *B. fungosa* var. *globosa (MN414176)* | 0.1986 | 0.1086 | 0.1050 | | | | | | | | | | | | | | | | | |
| *B. harlandii (MN414177)* | 0.1924 | 0.2199 | 0.2175 | 0.2252 | | | | | | | | | | | | | | | | |
| *B. japonica436 JAP* | 0.1951 | 0.1523 | 0.1471 | 0.1552 | 0.2215 | | | | | | | | | | | | | | | |
| *B. japonica592 JAP* | 0.1947 | 0.1519 | 0.1466 | 0.1553 | 0.2211 | 0.0011 | | | | | | | | | | | | | | |
| *B. japonica TWN* | 0.1943 | 0.1509 | 0.1474 | 0.1545 | 0.2206 | 0.0013 | 0.0006 | | | | | | | | | | | | | |
| *B. latisepala (SRR16213603)* | 0.2121 | 0.1725 | 0.1705 | 0.1798 | 0.2340 | 0.1819 | 0.1813 | 0.1812 | | | | | | | | | | | | |
| *B. laxiflora (KX784265)* | 0.1930 | 0.1499 | 0.1455 | 0.1510 | 0.2214 | 0.0839 | 0.0826 | 0.0825 | 0.1814 | | | | | | | | | | | |
| *B. mutinoides TWN* | 0.1778 | 0.2080 | 0.2056 | 0.2149 | 0.0572 | 0.2101 | 0.2097 | 0.2087 | 0.2253 | 0.2120 | | | | | | | | | | |
| *B. nipponica JAP* | 0.1863 | 0.1478 | 0.1442 | 0.1557 | 0.2101 | 0.1139 | 0.1133 | 0.1124 | 0.1759 | 0.1065 | 0.2006 | | | | | | | | | |
| *B. reflexa (KX784266)* | 0.2115 | 0.1748 | 0.1715 | 0.1779 | 0.2332 | 0.1786 | 0.1780 | 0.1781 | 0.1232 | 0.1780 | 0.2237 | 0.1717 | | | | | | | | |
| *B. subcupularis JAP* | 0.1887 | 0.1047 | 0.1008 | 0.0825 | 0.2174 | 0.1530 | 0.1535 | 0.1518 | 0.1724 | 0.1461 | 0.2048 | 0.1459 | 0.1705 | | | | | | | |
| *B. subcupularis (SRR16213604)* | 0.1902 | 0.1058 | 0.1026 | 0.0837 | 0.2190 | 0.1539 | 0.1544 | 0.1535 | 0.1739 | 0.1465 | 0.2072 | 0.1475 | 0.1725 | 0.0019 | | | | | | |
| *B. tobiracola JAP* | 0.1471 | 0.1996 | 0.1957 | 0.2083 | 0.1972 | 0.2038 | 0.2030 | 0.2026 | 0.2168 | 0.2028 | 0.1849 | 0.1920 | 0.2156 | 0.1980 | 0.1997 | | | | | |
| *B. tobiracola TWN* | 0.1470 | 0.1991 | 0.1955 | 0.2073 | 0.1974 | 0.2043 | 0.2034 | 0.2024 | 0.2165 | 0.2031 | 0.1869 | 0.1926 | 0.2142 | 0.1981 | 0.2002 | 0.0030 | | | | |
| *B. yakushimensis JAP* | 0.1869 | 0.1473 | 0.1438 | 0.1510 | 0.2115 | 0.0714 | 0.0713 | 0.0707 | 0.1745 | 0.0701 | 0.2021 | 0.1082 | 0.1711 | 0.1428 | 0.1440 | 0.1955 | 0.1962 | | | |
| *B. yakushimensis TWN* | 0.1863 | 0.1472 | 0.1438 | 0.1514 | 0.2114 | 0.0719 | 0.0715 | 0.0710 | 0.1744 | 0.0702 | 0.2021 | 0.1085 | 0.1713 | 0.1423 | 0.1433 | 0.1956 | 0.1962 | 0.0012 | | |
| *B. yakushimensis (NC_062820)* | 0.1869 | 0.1473 | 0.1438 | 0.1510 | 0.2115 | 0.0714 | 0.0713 | 0.0707 | 0.1745 | 0.0701 | 0.2021 | 0.1082 | 0.1711 | 0.1428 | 0.1440 | 0.1955 | 0.1962 | 0.0000 | 0.0012 | |

**Table S7. DNA sequencing and plastome assembly details for all studied species.** Question marks refer to SPAdes assemblies which were identical but incomplete. A short sequence comprising *rrn5* and the start of *ycf1* were missing from these three SPAdes assemblies.

| Sample | Voucher no. | Sequencing method | Number of reads | Plastome size (bp) | Number of nodes in SPAdes assembly | Identity with NOVOPlasty assembly (%) |
|---|---|---|---|---|---|---|
| *B. fungosa ssp. fungosa JAP* | X | Illumina NovaSeq 6000 | 68,565,854 | 14,968 | 4 | 99-100 |
| *B. japonica 436 JAP* | X | Illumina NovaSeq 6000 | 72,239,842 | 15,624 | >1 | 100? |
| *B. japonica 592 JAP* | X | Illumina NovaSeq 6000 | 64,340,476 | 15,646 | 1 | 100 |
| *B. nipponica JAP* | X | Illumina NovaSeq 6000 | 81,566,116 | 15,189 | >1 | 100? |
| *B. subcupularis JAP* | X | Illumina NovaSeq 6000 | 77,737,272 | 14,259 | 1 | 100 |
| *B. tobiracola JAP* | X | Illumina NovaSeq 6000 | 75,742,761 | 16,255 | >1 | 100? |
| *B. yakushimensis JAP* | X | Illumina NovaSeq 6000 | 104,551,614 | 14,631 | 1 | 100 |
| *B. fungosa* ssp. *fungosa TWN* | Su139 | Illumina MiSeq | 17,105,928 | 14,583 | >10 | 95-98 |
| *B. japonica TWN* | Su031 | Illumina MiSeq | 19,917,154 | 15,515 | >9 | 95-100 |
| *B. mutinoides TWN* | Su180 | Illumina MiSeq | 20,296,346 | 15,889 | 4 | 99 |
| *B. tobiracola TWN* | Su166 | Illumina MiSeq | 17,586,552 | 16,290 | 1 | 100 |
| *B. yakushimensis TWN* | Su099 | Illumina MiSeq | 1,218,295 | 14,628 | 1 | 100 |

**Table S8. RNA sequencing and transcriptome and proteome assembly details for studied species (JAP) and re-assembled publicly available transcriptomic data accompanied by accession numbers.**

| Sample | Sequencing method | Number of reads | Number of transcripts | Number of predicted proteins | Number of complete predicted proteins |
|---|---|---|---|---|---|
| *B. fungosa ssp. fungosa JAP* | Illumina NovaSeq 6000 | 37,467,431 | 222,041 | 46,439 | 27,109 |
| *B. japonica 436 JAP* | Illumina NovaSeq 6000 | 37,091,893 | 168,475 | 43,237 | 18,232 |
| *B. japonica 592 JAP* | Illumina NovaSeq 6000 | 46,817,996 | 291,323 | 103,553 | 60,788 |
| *B. nipponica JAP* | Illumina NovaSeq X | 29,114,553 | 113,070 | 34,972 | 20,723 |
| *B. subcupularis JAP* | Illumina NovaSeq 6000 | 52,214,335 | 170,916 | 60,650 | 37,805 |
| *B. tobiracola JAP* | Illumina NovaSeq 6000 | 46,724,888 | 160,944 | 47,903 | 32,858 |
| *B. yakushimensis JAP* | Illumina NovaSeq 6000 | 41,057,661 | 136,710 | 24,094 | 9,311 |
| *B. fungosa ssp. indica (SRR12009646)* | Illumina HiSeq 3000 | 20,377,191 | 102,767 | 31,946 | 21,074 |
| *B. fungosa (ERR2040275)* | Illumina HiSeq 2000 | 13,333,334 | 89,578 | 20,361 | 11,675 |
| *B. sp. (misidentified as B. harlandii; SRR12009647)* | Illumina HiSeq 3000 | 14,020,422 | 85,610 | 26,360 | 19,013 |
| *Lophophytum mirabile (SRR10883507)* | Illumina HiSeq 2500 | 295,533,285 | 562,212 | 107,735 | 65,559 |
| *Rhopalocnemis phalloides (SRR14800310)* | Illumina NovaSeq 6000 | 32,777,552 | 250,632 | 68,370 | 32,635 |

**Table S9. The number of nuclear-encoded plastid-targeted proteins across the *Balanophoraceae* proteomes predicted by Deeploc 2.0 (Thumuluri et al. 2022).** The second column includes all proteins with localization in the plastid (even when (an) additional localization(s) was/were predicted). Conservative filtering includes only proteins with "Chloroplast transit peptide" or "Thylakoid luminal transit peptide" as targeting signals. Plastid-targeted proteins (soft filtering) were clustered by CD-HIT (Li and Godzik 2006) at 98% and 95% sequence identities to identify the number of protein isoforms and recent paralogs. Proteomes predicted from re-assembled publicly available transcriptomic data are accompanied by accession numbers.

| Sample | Plastid-targeted (conservative filtering) | Plastid-targeted (all predicted) | Plastid-targeted protein clusters (98% identity) | | Plastid-targeted protein clusters (95% identity) | |
|---|---|---|---|---|---|---|
| *B. fungosa ssp. fungosa JAP* | 553 | 1,252 | 1,100 | 87.86% | 1,046 | 83.55% |
| *B. japonica 436 JAP* | 824 | 1,972 | 1,784 | 90.47% | 1,745 | 88.49% |
| *B. japonica 592 JAP* | 891 | 2,173 | 1,488 | 68.48% | 1,368 | 62.95% |
| *B. nipponica JAP* | 681 | 1,508 | 1,187 | 78.71% | 1,096 | 72.68% |
| *B. subcupularis JAP* | 1,457 | 3,079 | 2,532 | 82.23% | 2,405 | 78.11% |
| *B. tobiracola JAP* | 684 | 1,640 | 1,351 | 82.38% | 1,267 | 77.26% |
| *B. yakushimensis JAP* | 366 | 897 | 847 | 94.43% | 824 | 91.86% |
| *B. fungosa ssp. indica (SRR12009646)* | 614 | 1,407 | 1,162 | 82.59% | 1,093 | 77.68% |
| *B. fungosa (ERR2040275)* | 493 | 1,127 | 1,025 | 90.95% | 991 | 87.93% |
| *B. sp. (misidentified as B. harlandii; SRR12009647)* | 519 | 1,246 | 1,123 | 90.13% | 1,085 | 87.08% |
| *Lophophytum mirabile (SRR10883507)* | 1,397 | 3,134 | 2,354 | 75.11% | 2,092 | 66.75% |
| *Rhopalocnemis phalloides (SRR14800310)* | 931 | 2,103 | 1,704 | 81.03% | 1,605 | 76.32% |
| ***Averages*** | **784.17** | **1,794.83** | **1,471.42** | **83.70%** | **1,384.75** | **79.22%** |

### Supplementary Material 1: Phylogenetic trees

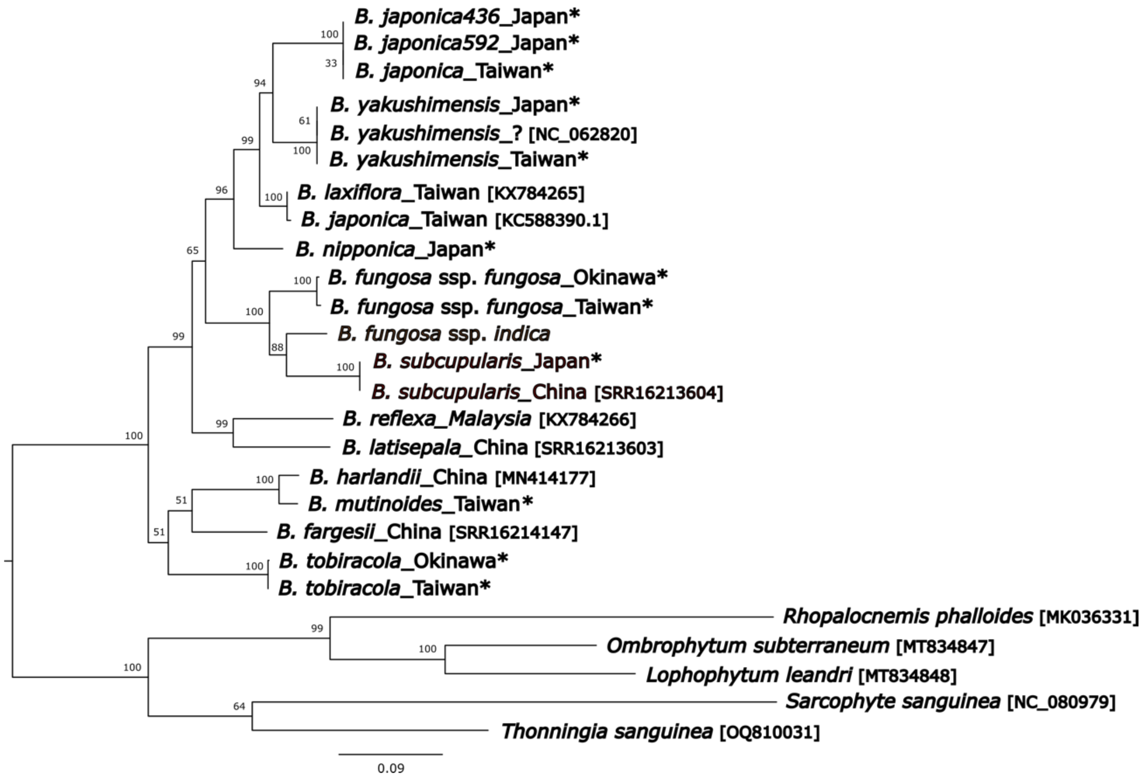


**SupplMat1A. Maximum likelihood phylogeny of Balanophoraceae inferred by IQTREE from plastid 16S rRNA gene (TVM+F+I+G4 model; 1,000 bootstraps).** The phylogenetic tree is rooted and bootstrap values are displayed next to branches. Sequences newly generated by this study are highlighted by asterisks. GenBank accession numbers for previously published data are indicated in the brackets.

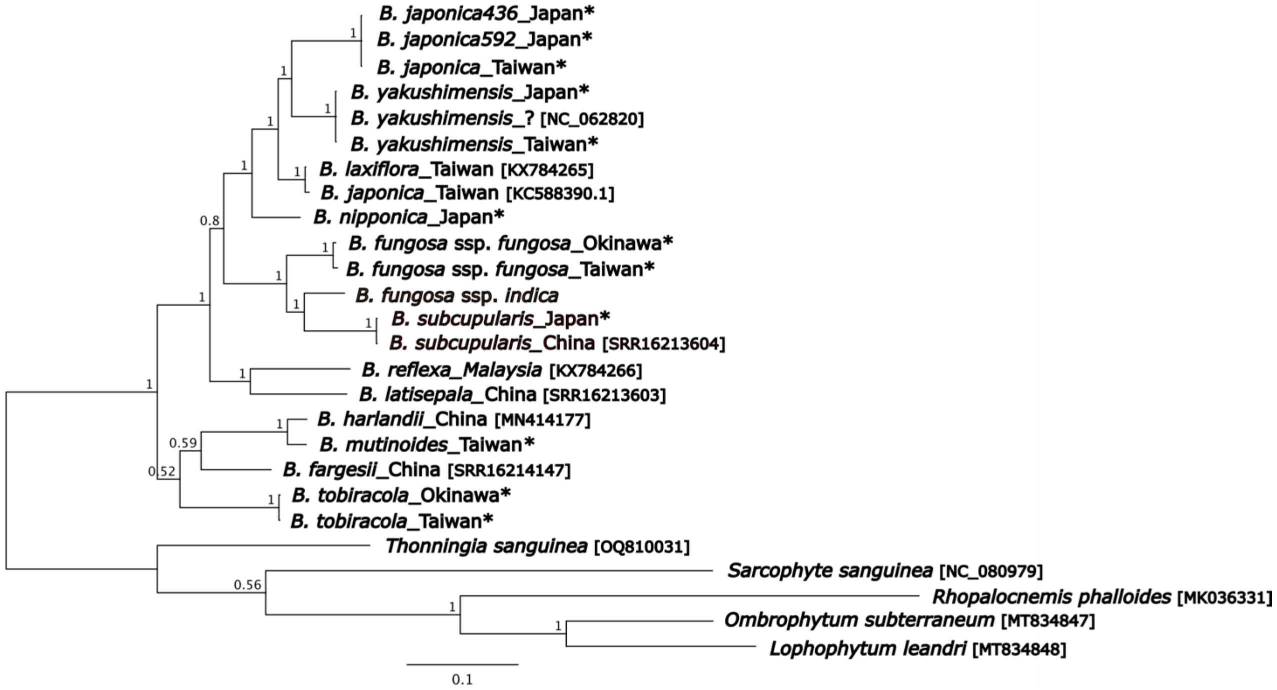


**SupplMat1B. Bayesian phylogeny of Balanophoraceae inferred from plastid 16S rRNA gene (GTR+G model; 1,000,000 generations).** Posterior probabilities are displayed next to branches. The phylogenetic tree is rooted. Sequences newly generated by this study are highlighted by asterisks. GenBank accession numbers for sequences obtained from NCBI are indicated in the brackets.

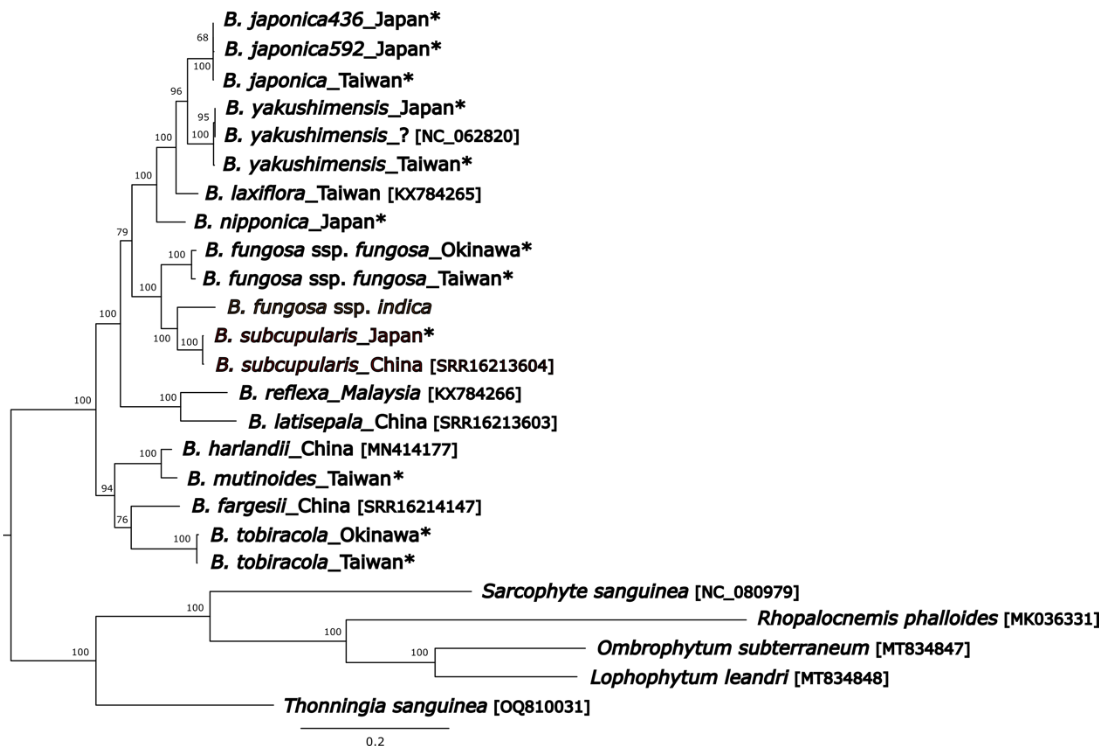


**SupplMat1C. Maximum likelihood phylogeny of Balanophoraceae inferred by IQTREE from plastid 23S rRNA gene (GTR+F+R3 model; 1,000 bootstraps).** The phylogenetic tree is rooted and bootstrap values are displayed next to branches. Sequences newly generated by this study are highlighted by asterisks. GenBank accession numbers for previously published data are indicated in the brackets.

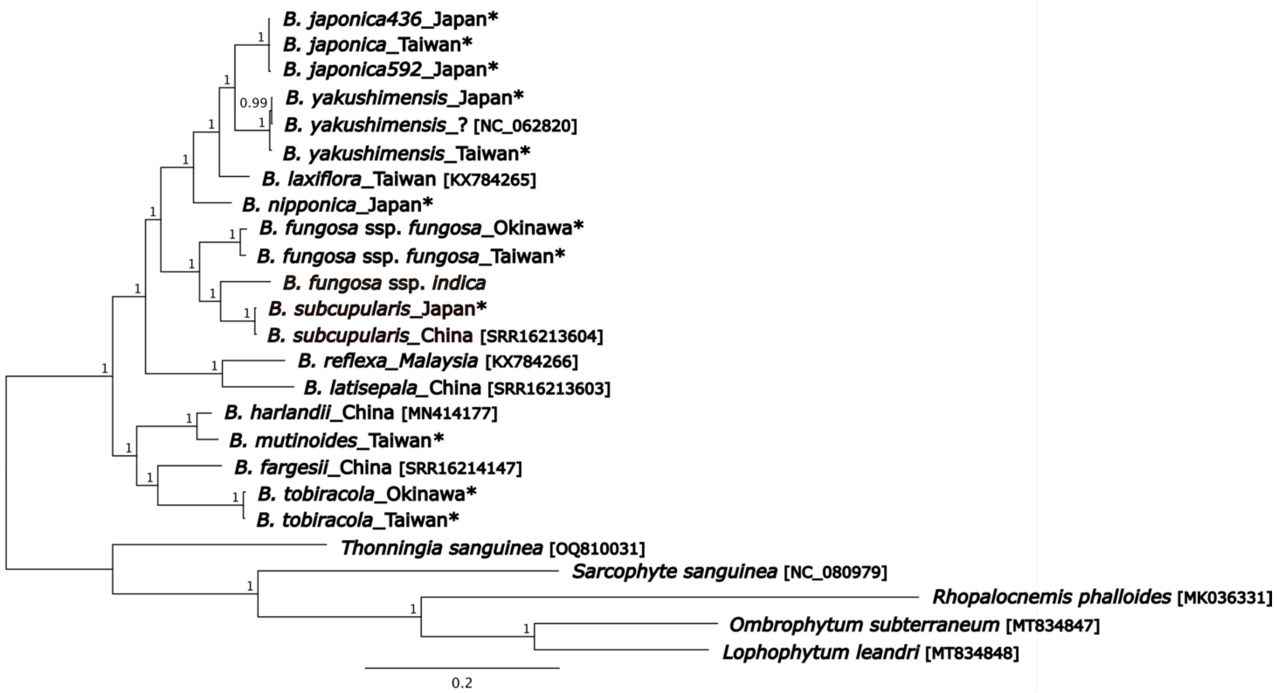


**SupplMat1D. Bayesian phylogeny of Balanophoraceae inferred from plastid 23S rRNA gene (GTR+G model; 1,000,000 generations).** Posterior probabilities are displayed next to branches. The phylogenetic tree is rooted. Sequences newly generated by this study are highlighted by asterisks. GenBank accession numbers for sequences obtained from NCBI are indicated in the brackets.

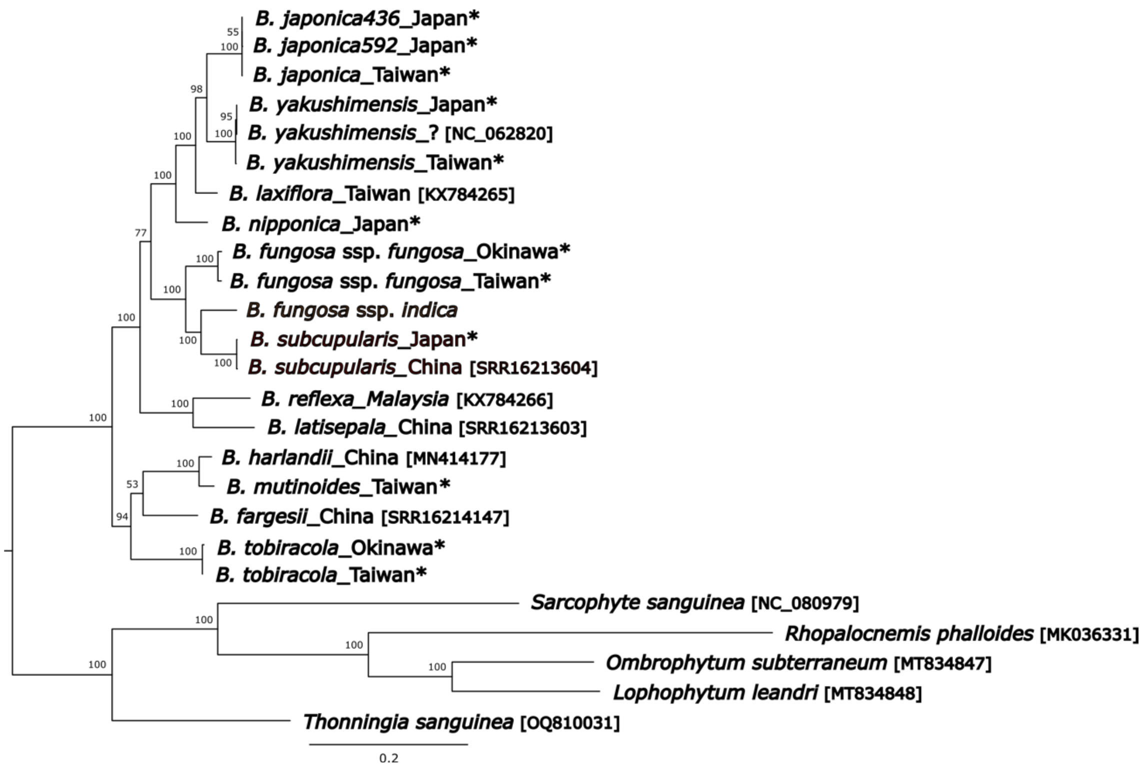


**SupplMat1E. Maximum likelihood phylogeny of Balanophoraceae inferred by IQTREE from concatenated plastid 16S and 23S rRNA genes (GTR+F+R3 model; 1,000 bootstraps).** The phylogenetic tree is rooted and bootstrap values are displayed next to branches. Sequences newly generated by this study are highlighted by asterisks. GenBank accession numbers for previously published data are indicated in the brackets.

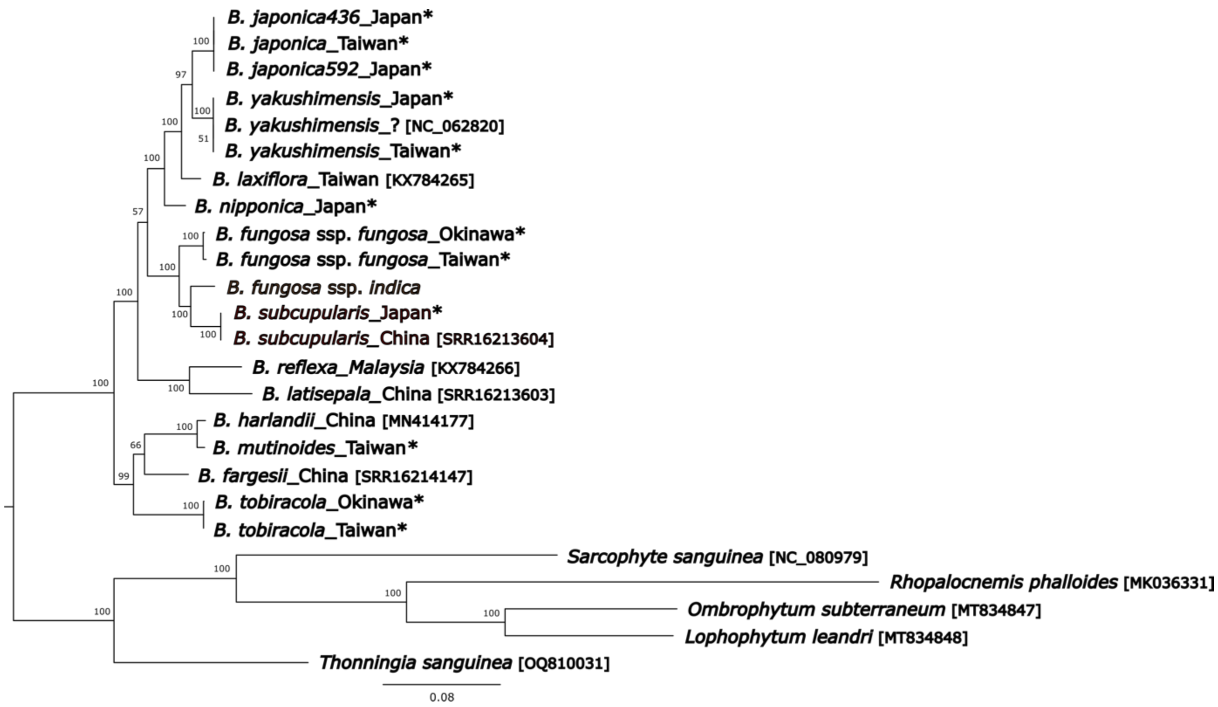


**SupplMat1F. Maximum likelihood phylogeny of Balanophoraceae inferred by IQTREE from concatenated plastid 16S and 23S rRNA genes (GTR+F+R3 model; 1,000 bootstraps).** The phylogenetic tree is rooted and bootstrap values are displayed next to branches. Sequences newly generated by this study are highlighted by asterisks. GenBank accession numbers for previously published data are indicated in the brackets. The dataset was trimmed by Gblocks before constructing the tree.

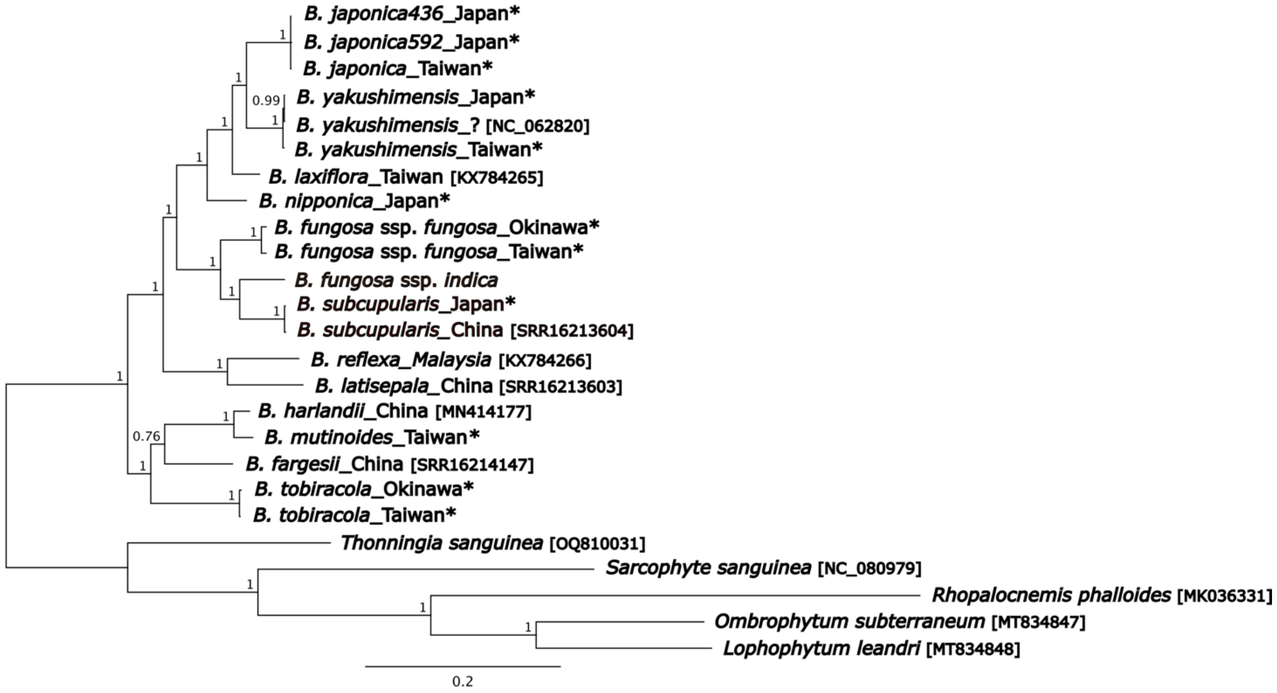


**SupplMat1G. Bayesian phylogeny of Balanophoraceae inferred from concatenated plastid 16S and 23S rRNA genes (GTR+G model; 10,000,000 generations).** Posterior probabilities are displayed next to branches. The phylogenetic tree is rooted. Sequences newly generated by this study are highlighted by asterisks. GenBank accession numbers for sequences obtained from NCBI are indicated in the brackets.

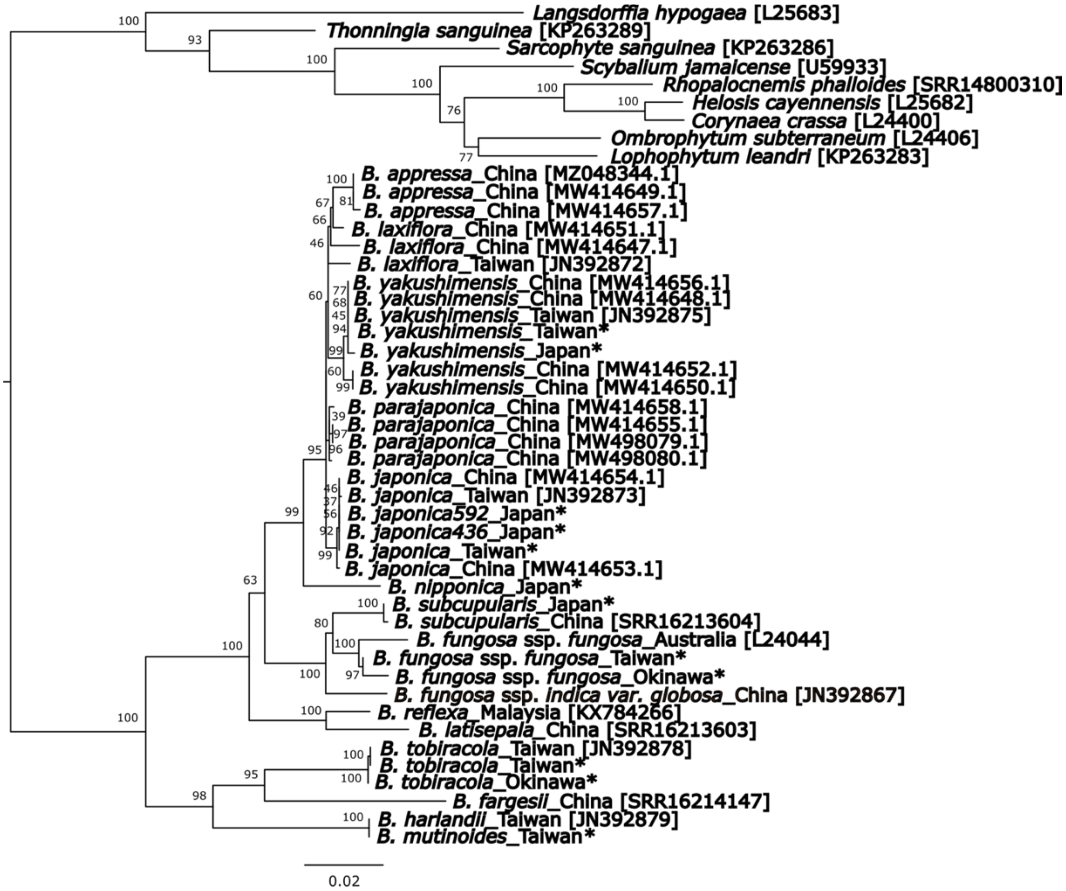


**SupplMat1H. Maximum likelihood phylogeny of Balanophoraceae inferred by IQTREE from nuclear 18SrRNA gene (GTR+F+I+G4 model; 1,000 bootstraps).** The phylogenetic tree is rooted and bootstrap values are displayed next to branches. Sequences newly generated by this study are highlighted by asterisks. GenBank accession numbers for previously published data are indicated in the brackets.

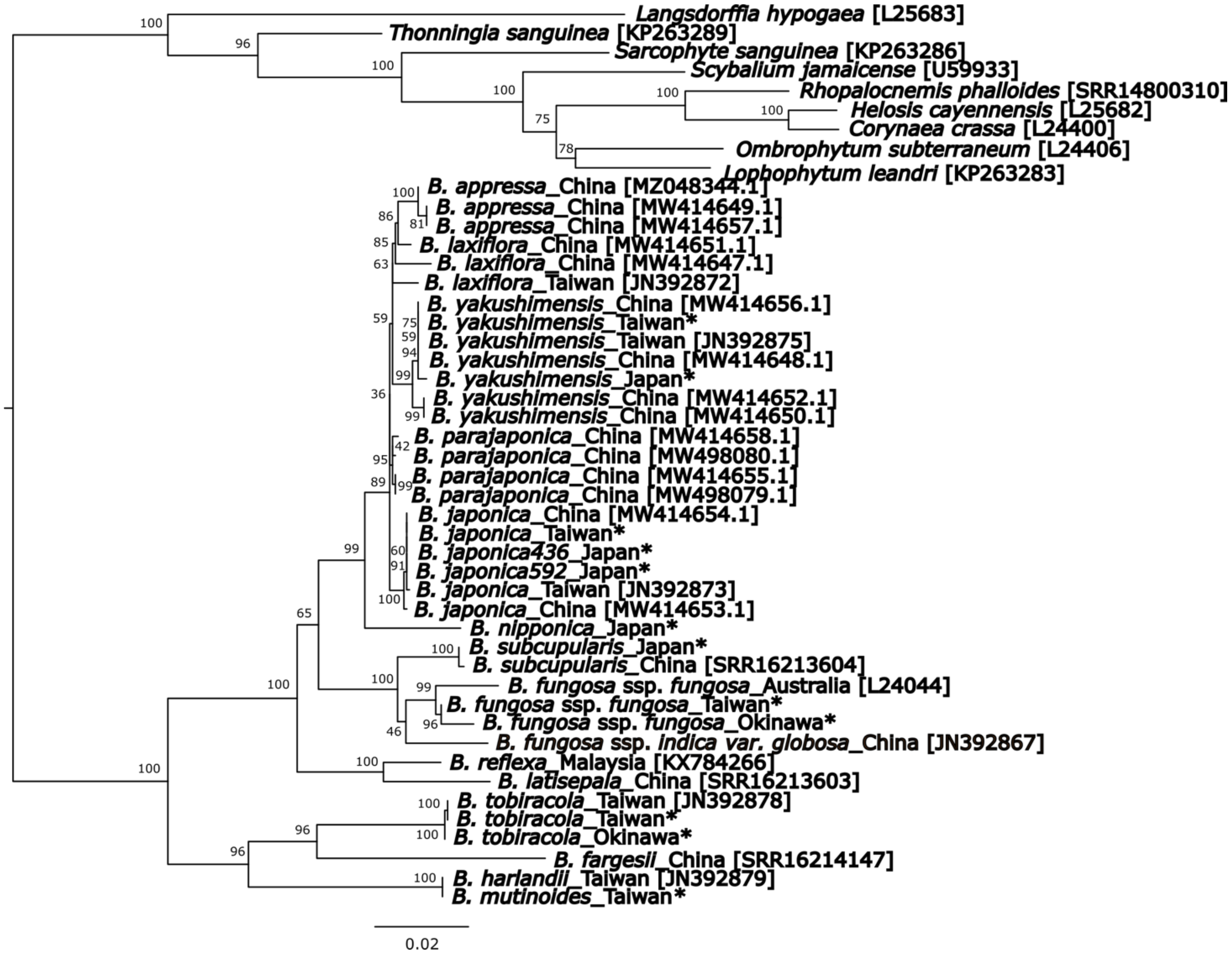


**SupplMat1I. Maximum likelihood phylogeny of Balanophoraceae inferred by IQTREE from nuclear 18SrRNA gene (GTR+F+I+G4 model; 1,000 bootstraps).** The phylogenetic tree is rooted and bootstrap values are displayed next to branches. Sequences newly generated by this study are highlighted by asterisks. GenBank accession numbers for previously published data are indicated in the brackets. The dataset was trimmed by Gblocks before constructing the tree.

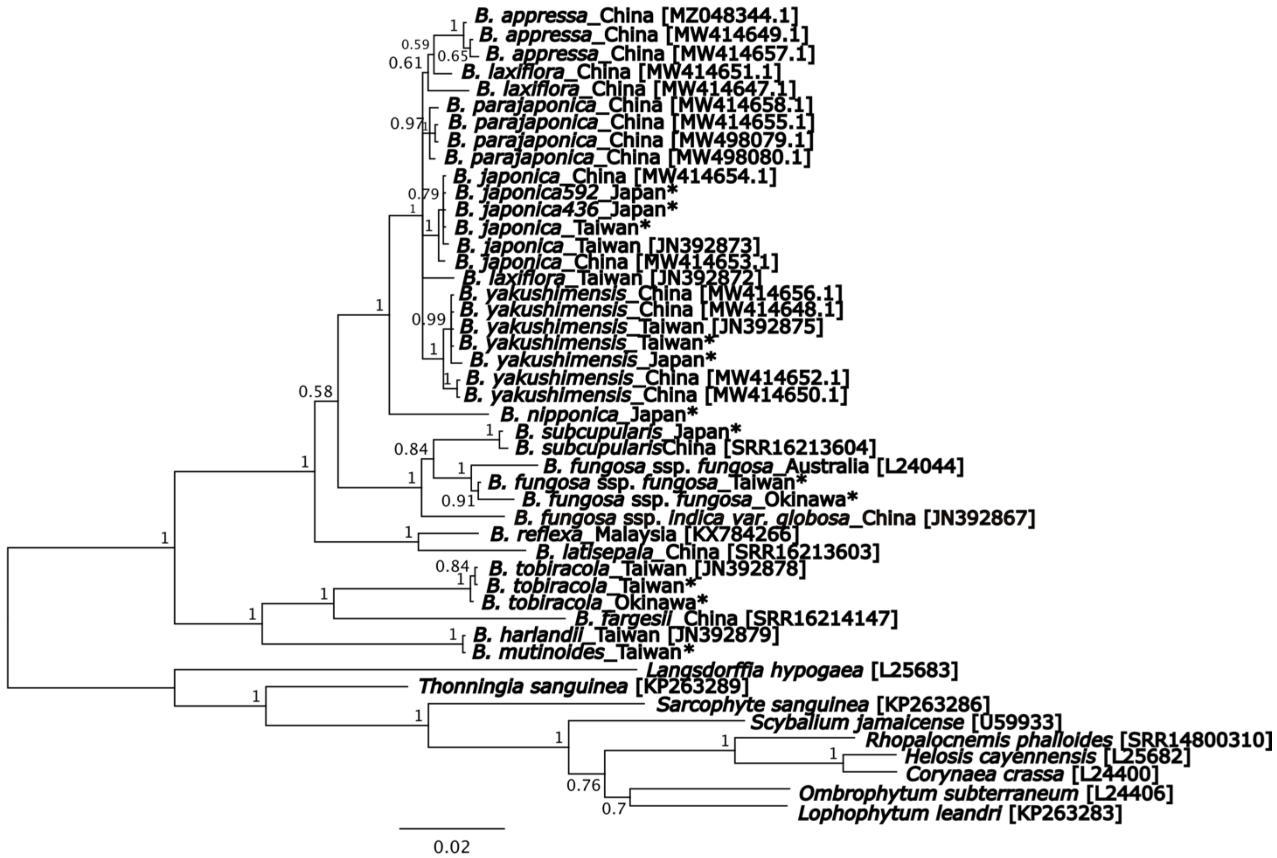


**SupplMat1J. Bayesian phylogeny of Balanophoraceae inferred from nuclear 18S rRNA gene (GTR+G model; 10,000,000 generations).** Posterior probabilities are displayed next to branches. The phylogenetic tree is rooted. Sequences newly generated by this study are highlighted by asterisks. GenBank accession numbers for sequences obtained from NCBI are indicated in the brackets.

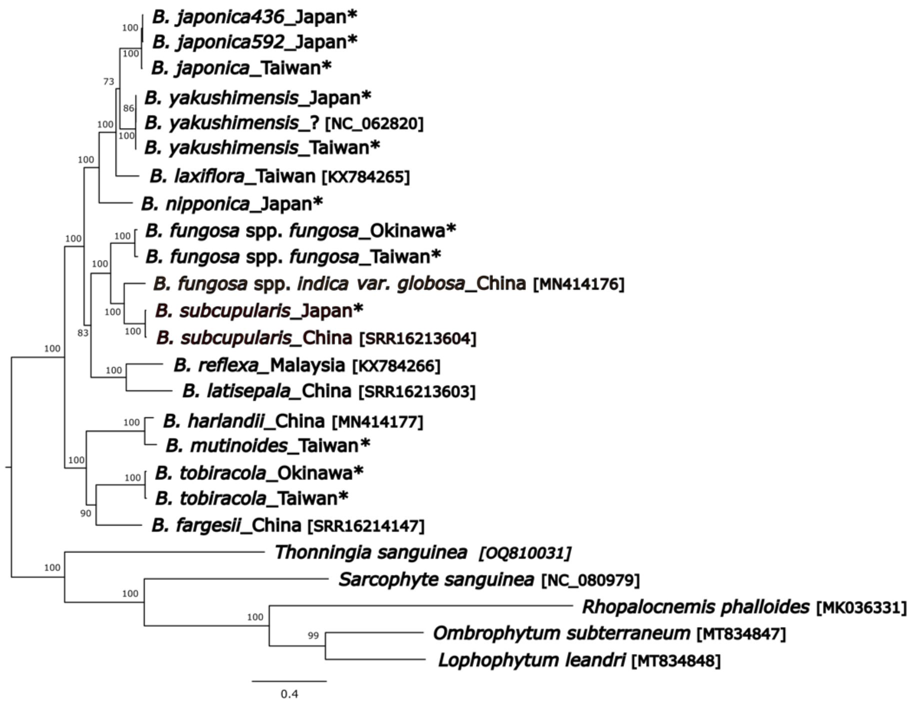


**SupplMat1K. Maximum likelihood phylogeny of Balanophoraceae inferred by IQTREE from a concatenated alignment of 15 protein-coding plastid genes (mtInv+F+I+G4 model; 1,000 bootstraps).** The phylogenetic tree is rooted and bootstrap values are displayed next to branches. Sequences newly generated by this study are highlighted by asterisks. GenBank accession numbers for previously published data are indicated in the brackets.

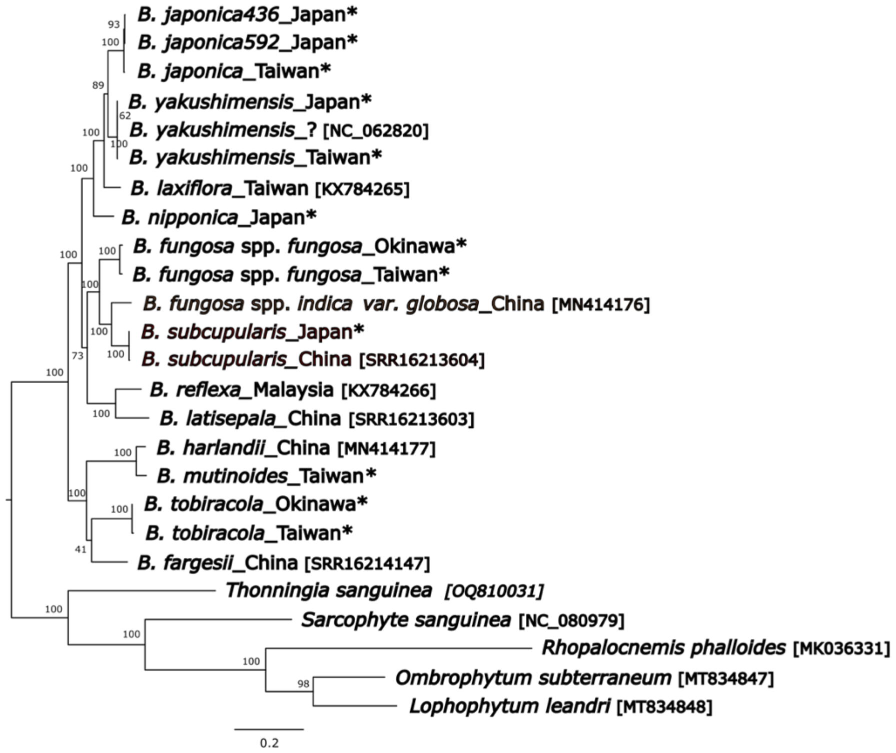


**SupplMat1L. Maximum likelihood phylogeny of Balanophoraceae inferred by IQTREE from a concatenated and trimmed alignment of 15 protein-coding plastid genes (JTT+F+I+G4 model; 1,000 bootstraps).** The phylogenetic tree is rooted and bootstrap values are displayed next to branches. Sequences newly generated by this study are highlighted by asterisks. GenBank accession numbers for previously published data are indicated in the brackets. The alignment was trimmed by by Gblocks before constructing the tree.

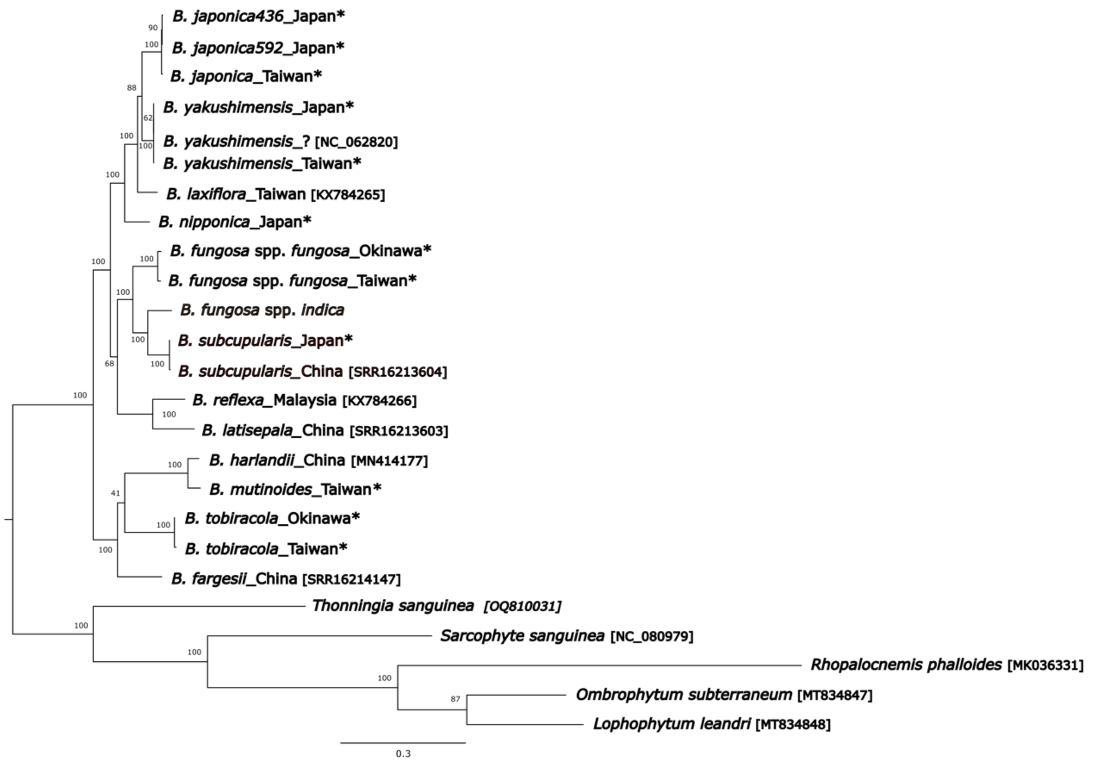


**SupplMat1M. Maximum likelihood phylogeny of Balanophoraceae inferred from a concatenated and trimmed alignment of 15 protein-coding plastid genes (LG+C20+F+G model; 10,000,000 generations).** The phylogenetic tree is rooted and bootstrap values are displayed next to branches. Sequences newly generated by this study are highlighted by asterisks. GenBank accession numbers for sequences obtained from NCBI are indicated in the brackets. The alignment was trimmed by Gblocks before constructing the tree.

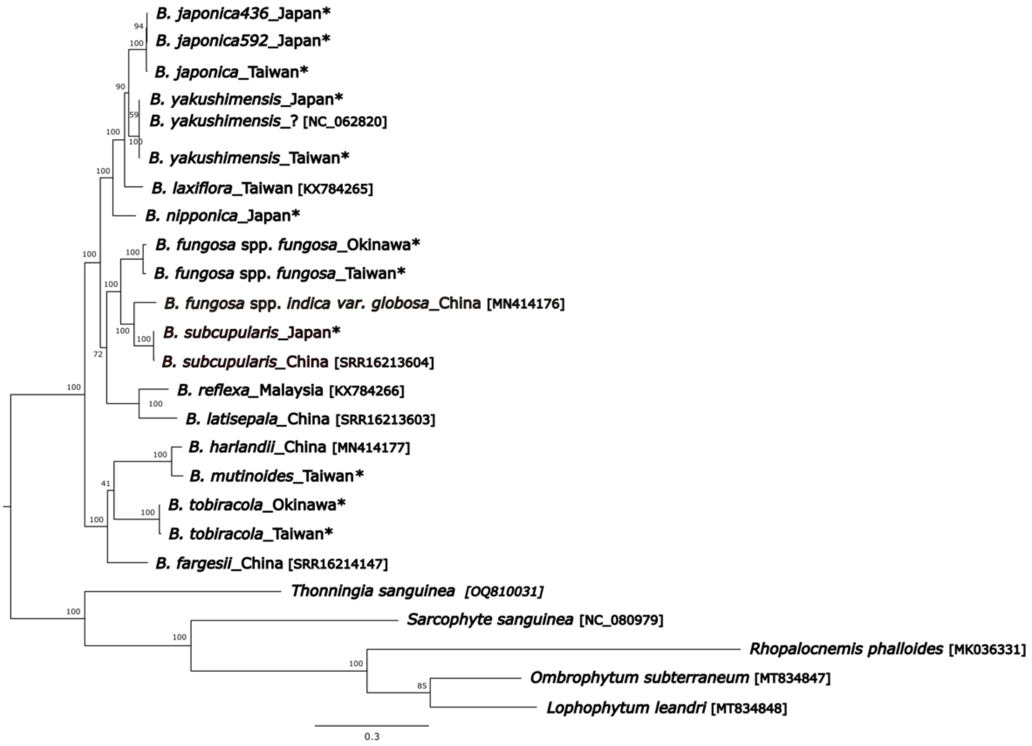


**SupplMat1N. Maximum likelihood phylogeny of Balanophoraceae inferred from a concatenated and trimmed alignment of 15 protein-coding plastid genes (LG+C40+F+G model; 10,000,000 generations).** The phylogenetic tree is rooted and bootstrap values are displayed next to branches. Sequences newly generated by this study are highlighted by asterisks. GenBank accession numbers for sequences obtained from NCBI are indicated in the brackets. The alignment was trimmed by Gblocks before constructing the tree.

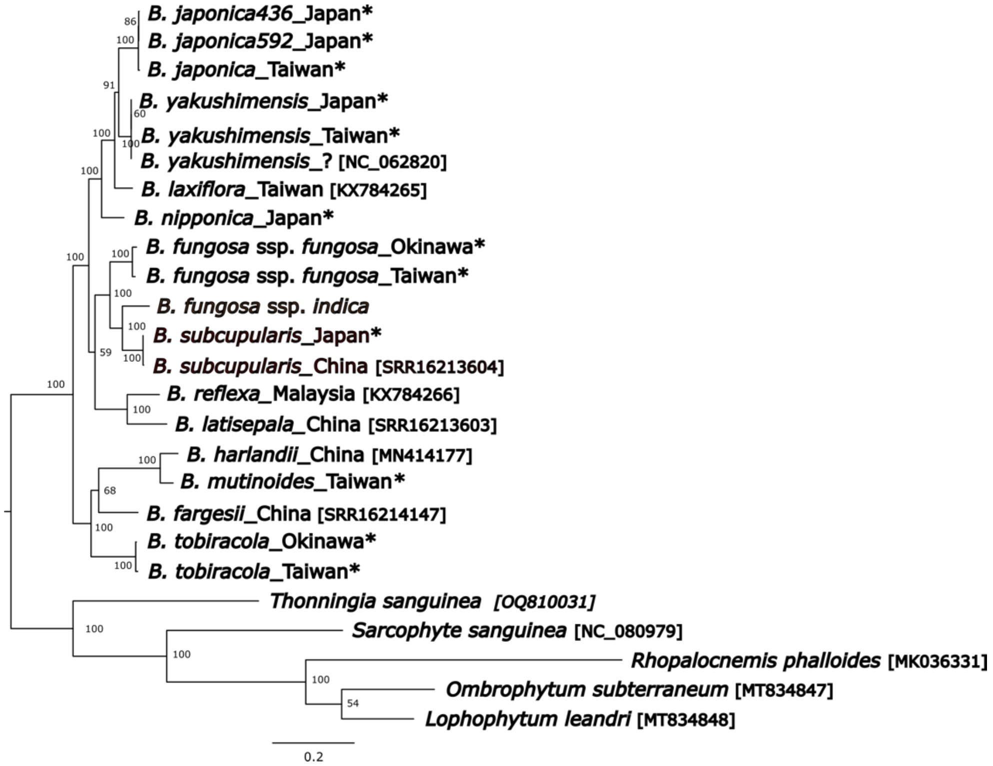


**SupplMat1O. Maximum likelihood phylogeny of Balanophoraceae inferred from a concatenated and trimmed alignment of 15 protein-coding plastid genes (LG+C60+F+G model; 10,000,000 generations).** The phylogenetic tree is rooted and bootstrap values are displayed next to branches. Sequences newly generated by this study are highlighted by asterisks. GenBank accession numbers for sequences obtained from NCBI are indicated in the brackets. The alignment was trimmed by Gblocks before constructing the tree.

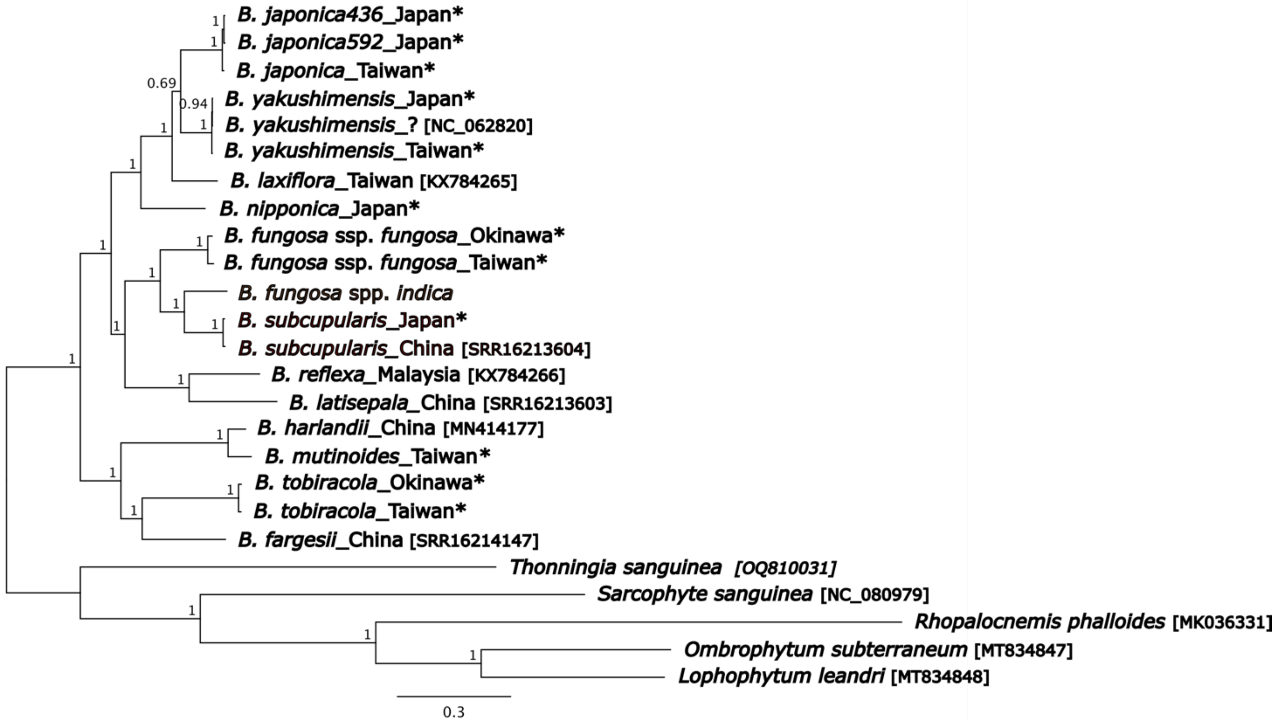


**SupplMat1P. Bayesian phylogeny of Balanophoraceae inferred from a concatenated alignment of 15 protein-coding plastid genes (LG+G model; 10,000,000 generations).** Posterior probabilities are displayed next to branches. The phylogenetic tree is rooted. Sequences newly generated by this study are highlighted by asterisks. GenBank accession numbers for sequences obtained from NCBI are indicated in the brackets.
